# Climate change is predicted to disrupt patterns of local adaptation in wild and cultivated maize

**DOI:** 10.1101/549675

**Authors:** Jonás A Aguirre-Liguori, Santiago Ramírez-Barahona, Peter Tiffin, Luis E Eguiarte

## Abstract

Climate change is one of the most important threats to biodiversity and crop sustainability. The impact of climate change is often evaluated on the basis of expected changes in species’ geographical distributions. Genomic diversity, local adaptation, and migration are seldom integrated into projections of species’ responses to climate change. Here we predict that climate change will impact populations of two wild relatives of maize, the teosintes *Zea mays* ssp. *mexicana* and *Z. mays* ssp. *parviglumis*, by altering patterns of local adaptation and decreasing migration probabilities. These alterations appear to be geographically heterogeneous across populations, suggesting that the possible impacts of climate change will vary considerably among populations. This in spite that most populations exhibit high levels of genetic diversity and are predicted to lie within future suitable areas. The heterogeneous patterns of local adaptation uncovered in teosintes are also evident across maize landraces, which suggests that climate change may give way to future maladaptation of several landraces within currently cultivated areas, possibly leading to increased chances of production shocks under increased temperatures. The predicted alterations to habitat distribution, migration potential, and patterns of local adaptation in wild and cultivated maize, raises a red flag for the future of populations. This underscores the need for continued integration of agronomical practices, genomic data, and climate models to better understand the impacts of a rapidly changing climate on cultivated and wild species.

## Introduction

Climate change is having detrimental effects on the geographic distribution, local abundance and levels of genetic diversity of species^1–6^. In particular, climate change induced population declines are one of the major contributors to the current loss of biodiversity. In cultivated species, these impacts will have significant economic and alimentary repercussions at local and global scales^2,4,7^. The development of strategies to mitigate the impacts of climate change on cultivated species has been a high priority, with recent proposals highlighting the importance of genetic diversity and local adaptation of wild relatives for crop improvement and mitigation^4,8–12^.

Maize (*Zea mays* ssp. *mays*) was domesticated in Mexico from its wild relatives, the teosintes, nearly 9,000 years ago^13^. More than 200 Latin-American landraces of maize have been described, which are the result of past and on-going sociocultural processes that shape and maintain extraordinary biodiversity in this species^14,15^. Teosintes grow wild under a wide range of climatic conditions, from very hot and humid coastal environments to temperate and dry inland regions^10,16,17^. Maize land races have an even wider geographic and environmental range^5,18,19^. Evidence of gene flow from teosintes into maize and the presence of fertile hybrids^20,21^ indicates that gene introgression into landraces is possible and could be important to the continued evolution of maize to respond to new environmental challenges^16,22^.

Both the teosintes and several maize landraces are threatened by habitat loss and climate change^4,7,19,23^, primarily increasing temperature and aridity^4,6^. Local adaptation to warm and dry climates is common in teosintes^16,24–26^. Populations locally adapted to warm and dry environments are likely to contain alleles that could reduce the negative effects of global warming and increased aridity. Thus, the local adaptation of teosintes populations, which has not been directed by cultural and agronomical practices, might prove useful as a blueprint of local adaptation for crop improvement and mitigation in maize landraces.

Ecological niche modeling has become a powerful tool to assess the impacts of climate change on species’ geographic distribution and persistence^27–29^. Commonly used ecological niche models assume that populations within a species are identical and share the same environmental tolerance limits, disregarding differences in the local adaptive response of populations to climate^30,31^. However, given that local adaptation is common, among-population variation in climatic tolerances and varying levels of climate-gene relationships across the genome are expected^25,30^. To our knowledge only a couple of studies have explicitly integrated ecological niche modeling and genomic data to more realistically assess species’ response to climate change^30,32,33^.

Here we examined how climate change is expected to affect the geographic distribution and the frequency of single nucleotide polymorphisms (SNPs) associated with climate-related traits in two teosintes species: *Z. mays* ssp. *mexicana* and *Z. mays* ssp. *parviglumis* (hereafter referred to as *mexicana* and *parviglumis*, respectively). These teosintes thrive under distinct environmental conditions: *mexicana* is native to the highland regions of Central Mexico, *parviglumis* is native to the lowlands of South-western Mexico^10,16,17,25^.

## Materials and Methods

### Ecological niche models

We used present day climatic variables to build species distribution models for the two teosintes separately as implemented in Maxent v.3.3.3^34^. We used 254 (*mexicana*) and 329 (*parviglumis*) occurrence data points and the bioclimatic variables available in WorldClim^35^ at a resolution of 30arc/sec (~1 Km^2^), with an area extent of 90°-120° W, 10°-30° N, and background sampling points randomly chosen across the study area (SI Appendix). The eight climate models included: two general circulation models (Community Climate System Model (CCSM), Model for Interdisciplinary Research on Climate, MIROC), two greenhouse gas concentration models adopted by the Intergovernmental Panel on Climate Change (Representative Conservation Pathways: RCP 4.5, RCP 8.5) and two future projections (2050, 2070). After validating the models, we generated binary presence/absence maps to quantify the degree of change in the geographic distribution of the two teosintes species (SI Appendix).

### Population genomic data

We merged the datasets of ref.^24^ and ref.^25^ (available at: http://dx.doi.org/10.5061/dryad.8m648 and https://doi.org/10.5061/dryad.tf556), both obtained using the MaizeSNP50 Illumina Bead chip, to construct a dataset of 33,454 high quality SNPs distributed along the ten chromosomes of maize^24,25^. The dataset comprises 23 populations for *mexicana* and 24 populations for *parviglumis*, covering the entire geographic and environmental distribution of teosintes^25^. The sets of locally adapted SNPs used here are described in detail by ref.^25^, who used *Bayescenv*^36^ and *Bayenv*^37^ to identify locally adapted SNPs with significant genetic differentiation among populations (*F*_*ST*_) and with significant associations with temperature and precipitation after controlling for geographic clines (SI Appendix). To define the set of candidate SNPs (canSNPs) and putative adaptive SNPs (paSNPs), we performed a Discriminant Analyses on Principal Components (DAPC) using climatically defined groups of populations and the locally adapted SNPs for each species separately using the *adegenet* package^38^ in R^39^ (SI Appendix). We define the set of paSNPs as those with a high power to discriminate populations (*i.e.*, values greater than the third quartile of distribution of loadings) (see SI Appendix, fig. S1), and the set of canSNPs as those with a lower discrimination power.

We estimated levels of genetic diversity (*Hs*) for refSNPs using the *adegenet* package^37^ and *hierfstat* package^40^ in R^39^. We characterized genetically poor populations as those with an *Hs* lower than 0.2 for refSNPs, because lower *Hs* appears to be the probable result of recent bottlenecks^24^.

### Gradient Forest (GF)

Recently, species ecological distribution modeling techniques have been developed to consider intraspecific genetic variation in the assessment of the impact of climate change^29,30,32,33^. Among these, GF^41^ is a regression tree-based approach that implements non-linear regressions to estimate patterns of turnover in species (or allelic) composition as a function of environmental variables^30^ (SI Appendix). The algorithm identifies regions along an environmental gradient associated with a high rate of change in allele frequencies, irrespective of the underlying allele frequencies. GF uses this information to construct allele turnover functions^30^, which can be used to model the change in allele frequencies throughout a species’ range as a function of climatic variables.

We used the *gradientforest* package^41^ in R^39^ to construct turnover functions for each SNP dataset separately (e.g., refSNPs, canSNPs, paSNPs) with the 19 bioclimatic variables used for the species distribution models (see above). We estimated the allelic frequencies of all SNPs in each dataset and implemented GF using 100 trees and a correlation threshold of 0.5^30^. Based on the best allele turnover models for each set of SNPs, we constructed landscape models for the present and for the eight future climate models using the *raster* package^42^ in R^39^. We compared present-day and future landscape models to identify regions (and populations) where the current gene–environment relationships will be most disrupted (genomic offset) (SI Appendix), which measures the relative magnitude of allele frequency changes expected as result of climate change^30,32,33^.

To gauge the robustness of the GF models to the use of different sets of SNPs, we ran additional analyses using: (1) a set of reference SNPs with an allele frequency distribution similar to those of canSNPs and paSNPs; (2) different sets of outlier SNPs identified by ref.^25^; and (3) on the complete set of 33,454 SNPs (SI Appendix).

### Allele distribution models

We predicted the geographic distribution for putative adaptive alleles at paSNPs using Maxent v.3.3.3^34^. Since we were interested in modeling the distribution of the putatively warm-adapted alleles (SI Appendix), we used the corresponding populations’ geographic coordinates where these alleles were recorded as input. We used the same settings and validation procedures used for the species distribution models (see above).

We generated binary presence/absence distribution models for each paSNP for the present and for future climate models (SI Appendix). We estimated the geographic overlap between the present model and each future model to define suitable areas for local adaptation as those with at least five paSNPs predicted to occur. We cross validated the allele distribution models using the GF models for paSNPs, basically inspecting the genomic offset of populations for three sets of regions: present-only, future-only, and overlap.

### Barriers to migration

Based on the overlapped distribution models for the paSNPs, we tested the potential capacity of populations to migrate into the new regions predicted under future climate scenarios using circuit theory^43^. This represents a simplified model of population migration based on the distribution of paSNPs and assumes that migration would be mostly limited by local adaptation. We constructed maps of potential migration using the present and future distribution models for paSNPs to determine landscape resistance (environmental distances) as a proxy of limitation to the successful migration, where increasing resistance indicates decreasing probabilities of migration (SI Appendix). For each sampled population, we constrained our analyses to a 1°x1° degree grid-cell centered on each sampled population. We used the 10-percentile of resistance values as a minimum threshold to estimate migration potential, which can be interpreted as the resistance to successful migration into at least 10% of the future areas of potential settlement (SI Appendix).

### Identifying putative adaptive SNPs in maize landraces

We explored whether the paSNPs identified for teosintes have been documented in particular landraces of maize^44^. For this, we downloaded the genotype data for maize described by ref.^44^, consisting of 36,931 SNPs across 46 landraces in Mexico, with 1 to 16 accessions per landrace. After identifying putative adaptive alleles at paSNPs across landraces, we estimated a relative ‘adaptive’ score by adding up individual allele frequencies in each landrace. Due to the varying number of accessions per land race, we corrected the adaptive score by the maximum number of alleles possible in each landrace (i.e., for each locus, four alleles for two accessions) (SI Appendix).

We evaluated the correlation between the adaptive scores and temperature, genomic offset and the geographic extension of each landrace (SI Appendix). For this, we employed simple linear regressions using the adaptive score as the response variable and climatic and geographic data for landraces as the predictors. We used the estimated genomic offset for the two teosintes under the less extreme climate change models (CCSM_2050_RCP4.5) to extract mean genomic offset values from the geographic occurrences of each landrace. To assess the possibility of uncovering positive correlations between predictors and the adaptive score of landraces irrespective of their potential for adaptation, we tested the correlations using adaptive scores estimated from 1000 different sets of randomly sampled refSNPs (13 per set).

## Results and Discussion

We employed ecological niche modeling and climate averages from 1960 to 1990^35^ to build geographic distribution models for the two teosintes species under eight projections of climate change. As expected due to geographic constraints^6,19,45^, the highland teosintes (*mexicana*) are predicted to be more susceptible to climate change than the lowland teosintes (*parviglumis*)^17,19^. These models predict that the potentially suitable geographic range for *mexicana* will be reduced by 30-39% in the next 30 to 40 years, whereas *parviglumis* range is predicted to be reduced by 4-9% (see SI Appendix, fig. S2, table S1). Consistent with the predicted contraction in geographic range, 29% and 15% of populations in *mexicana* and *parviglumis*, respectively, are predicted to fall outside of suitable habitat by 2050 (45% and 29% by 2070).

Ecological niche models can be used to predict suitable geographic ranges^27,29^. Intuitively, the predicted shifts and shrinkage in the teosintes’ geographic range represent a potential threat to their persistence under climate change^19^. However, these models do not provide direct information on the potential effects that climate change will have on the genetic composition of local populations^31^. To examine the potential effect of climate change on the genetic composition of local populations we applied an extension of Gradient Forest^41^ developed by ref.^30^ to predict how climate change will alter the frequencies of three sets of SNPs selected from a dataset of 33,454 SNPs (see SI Appendix) previously identified to be segregating within a sample of 23 populations of *mexicana* and 24 populations of *parviglumis* ^24,25^. The sampled populations were collected from throughout the teosintes’ distributions.

We subdivided the genomic dataset into three sets of SNPs. The first set is comprised by nine *parviglumis* and eight *mexicana* SNPs with frequencies that differ strongly among populations living either in warm or cold environments, a signal of local adaptation. Several of these SNPs are in or near genes involved in responses to heat, drought, and other environmental stress and we hereafter referred to these as putatively adaptive SNPs, or paSNP (see SI Appendix, table S2). The second set, hereafter referred to as candidate SNPs or canSNPs, include 24 *parviglumis* and 25 *mexicana* SNPs that also are significantly associated with climate, but not as strongly as paSNPs (see SI Appendix). The third set is comprised of 500 reference SNPs (refSNPs) that we randomly selected from the 33,454 SNPs that showed some, but not particularly strong population differentiation (*F*_*ST*_ values within the 10% and 90% percentiles of the distribution of per-SNP pairwise population *F*_*ST*_). Our expectations are that the paSNPs will not only be most strongly affected by climate change, but also will have a greater contribution to population adaptation to new climatic conditions than can or refSNPs.

Using the extension of Gradient Forest, we estimated environment-SNP associations by modeling allele frequency changes along present climatic gradients^41^. These models, in combination with projections of future climate change, can then be used to predict temporal patterns of within-population genetic turnover throughout a species geographic range^30,41^. The extent of genetic turnover of the paSNPs and the canSNPs relative to refSNPs, which we hereafter refer to as genomic offset^30,32,33^, provides insight into the disruptive effect of climate change on the frequencies of alleles that bear the strongest signatures of local adaptation among contemporary populations. Higher genomic offset is indicative of stronger shifts in frequencies of locally adapted alleles than populations having a lower genomic offset.

Although a more pronounced geographic contraction is predicted for *mexicana* than *parviglumis*, genomic offset at paSNPs and canSNPs is higher in *parviglumis* than in *mexicana* (mean = 0.27 vs. 0.06 across all models, respectively; see SI Appendix, tables S3, S4, S5). The higher genome-wide offset in *parviglumis* (see SI Appendix, figs. S3, S4) likely reflects greater population structure in *parviglumis* than *mexicana*^25^ and might also indicate that the genetic composition of *parviglumis* populations will be more affected by climate change than *mexicana* populations. The levels of genomic offset are, however, highly variable among populations (fig. 1a-b), with genomic offset lower for populations either with higher frequencies (> 0.5) of the adaptive alleles at paSNPs or having little expected change in climatic conditions (see SI Appendix, fig. S5). The among-population variation underscores that both the selective impact of climate change, and the genomic consequences of subsequent evolutionary response, may both vary among populations and are affected by past local adaptation.

**Figure 1.**
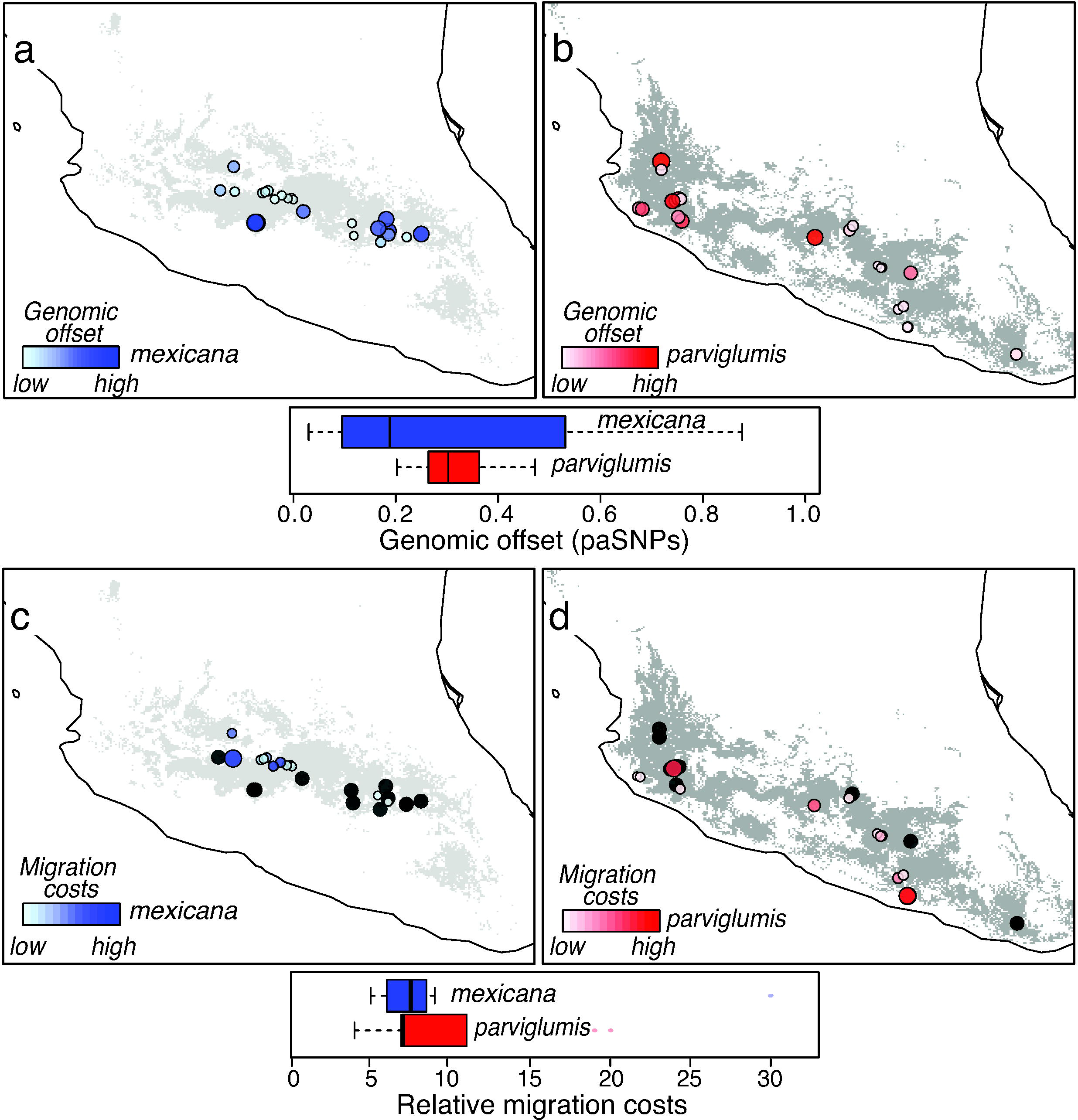
Genomic offset in climate-related SNPs and limited migratory potential for sampled populations of two teosintes in Mexico: *Zea mays* spp. *mexicana* and *Zea mays* spp. *parviglumis*. **a-d**, Distribution of genomic offset (a-b) and relative migration costs (c-d) for sampled populations of the two species of teosintes estimated under the climate change model CCSM_2050_RCP4.5. Light and dark gray areas represent the species distribution models for *mexicana* and *parviglumis*, respectively. Circle size is proportional to the estimated genomic offset and migration costs, with white circles representing populations with the lowest genomic offset and migration costs. Black circles in panels c and d represent populations with no migration potential (infinite migration costs) according the allele distribution models.

The considerable levels of genomic offset predicted in most populations have the potential to greatly alter patterns of local adaptation, or at least the genetic basis of local adaptation. The genomic offset may also limit populations’ adaptive potential, thereby leading to reduced population sizes and possibly higher probabilities of local extinction^3,25,30,32,33,45^. This points towards more complex impacts of climate change on wild populations than those predicted using geographical distribution models alone^31^.

To complement the use of Gradient Forest, we used the same modeling tools used to predict species distributions to predict the present and future geographic distribution of the putatively adaptive alleles at each of the paSNPs (see SI Appendix). This modeling showed that by 2050, the geographic distribution of adaptive alleles is expected to contract by ~ 45% in *mexicana* and by 36% *parviglumis* (table S6). The geographic areas not suitable for adaptive alleles in the future also show higher levels of genomic offset (see SI Appendix, fig. S6), indicating that areas outside the predicted distribution of adaptive alleles are expected to have big shifts in climate-gene relationships, either due to a lower starting frequency of adaptive alleles or to a larger change in climate.

Adaptive alleles at paSNPs are expected to confer selective advantages to expanding warmer and drier environments, yet populations lacking these adaptive alleles might follow a different evolutionary trajectory to adapt to climate change. In this context, standing ‘neutral’ genetic diversity could act as an important source of variation for future local adaptation^1,5,46,47^. We found a significantly negative association between genetic diversity at refSNPs (*Hs*) and the predicted genomic offset for paSNPs (see SI Appendix, fig. S7). These results suggest that, on average, genetically rich populations will be less vulnerable to climate change, probably because the adaptive potential of populations is maximized with intermediate allele frequencies. This result is consistent with the importance usually given to standing genetic variation to allow responding to a changing climate^46,47^. However, more than two-thirds of teosintes populations (71% in *parviglumis* and 91% in *mexicana*) show high levels of genetic diversity (*Hs* > 0.2, ref. ^24^), yet even under the most conservative climate model, these populations will experience widely varying levels of genomic offset (fig. 1a-b).

While some populations might adapt to the conditions of a changing climate, other populations might migrate to track the climatic conditions to which they are currently adapted^48,49^. With a focus on the alleles of major effect, we evaluated the migratory potential^43,50^ for sampled populations from their current location into sites with climatic conditions that are predicted to be suitable for the current frequencies of adaptive alleles. To do this, we used the distribution models for adaptive alleles to estimate environmental costs (i.e., lower probabilities of populations successfully migrating^43,50^) along all potential migration routes radiating from each sampled population.

Depending on the climate model used, as the climate changes, 46-54% of *parviglumis* populations and 22-52% of *mexicana* populations are expected to have the possibility of migrating into sites in which the alleles of major effect will remain advantageous (see SI Appendix, table S7). Although more *mexicana* populations are predicted to have no migration capabilities (*i.e.*, infinite migration costs) into climatically suitable sites, probably due to isolation from areas predicted to have suitable conditions in the future, the remaining populations of *mexicana* actually showed lower migration costs than populations of *parviglumis* (fig. 1c-d). Thus, although migration costs are expected to be higher for *parviglumis* than for *mexicana*, populations inhabiting the Central Highlands of Mexico are expected to have the highest migratory constraints (fig. 1c). The variation in migrations costs might result from the differing rates of temperature change estimated between highland environments compared to lowland regions^51^. The predicted likelihood of successful migration into a climatically suitable location was positively correlated with within-population frequency of adaptive alleles (see SI Appendix, fig. S8).

Under the most conservative model of climate change (CCSM_4.5_2050), our species distribution models predict 88% and 84% of sampled populations within future suitable areas for *parviglumis* and *mexicana*, respectively (see SI Appendix, table S1). In addition, most of these populations exhibit high levels of genetic diversity^25^. However, the heterogeneous geographical patterns of genomic offset and migration potential suggest that the impacts of climate change will vary considerably among populations. Overall, eastern populations of *mexicana* and western populations of *parviglumis* show high levels of genomic offset and lower migration potential (fig. 1), thus we expect these populations will experience stronger pressures to adapt to climate change than other teosintes populations. From this simple comparison, we conclude that although highly useful, distribution models are an oversimplification of the complex ecological and evolutionary consequences of climate change^28,29^.

Locally adapted populations of teosintes can be used as a source of advantageous genetic diversity for landraces of maize^4,17,22^, through the introduction of ‘new wild alleles’ into cultivated populations^8,9,11,12^. Experimental evidence has proven that teosintes germplasm can be successfully transferred into maize and can increase crop yields and drought resistance^14,15,21,52^. Alternatively, maize improvement can be achieved more easily by selecting adaptive alleles already present in maize landraces. We explored the frequency at which paSNPs identified for teosintes have been documented in some Mexican landraces of maize^44^, finding adaptive alleles in 46 landraces of maize (see SI Appendix, table S8). This search allowed us to estimate a relative ‘adaptive’ score by adding up the individual frequencies of adaptive alleles for available accessions in each landrace (SI Appendix).

Expanding on the varying levels of geographic contraction predicted for different maize landraces^19^, we found an uneven presence of adaptive alleles across distinct landraces in Mexico (fig. 2a), with a positive trend of increasing frequency of adaptive alleles (*i.e.*, adaptive score) in larger geographic ranges and higher mean annual temperatures (fig. 2c-d). Although the positive trend uncovered for range size is non-significant, several landraces with narrow distribution ranges, such as the almost extinct ‘Tehua’ landrace, and other highly restricted landraces^14^ (depicted as open circles in fig. 2b-e), tend to have adaptive alleles at fewer paSNPs and with lower frequencies. In turn, we found a negative relationship between adaptive scores and the genomic offset (fig. 2e), but this was statistically significant only for the CCSM_2070_RCP8.5 model. However, adaptive scores estimated with refSNPs showed a weaker association with temperature and genomic offset than those estimated for paSNPs, with less than 0.5% of the refSNPs subsamples showing a stronger correlation (see SI Appendix, fig. S9).

**Figure 2.**
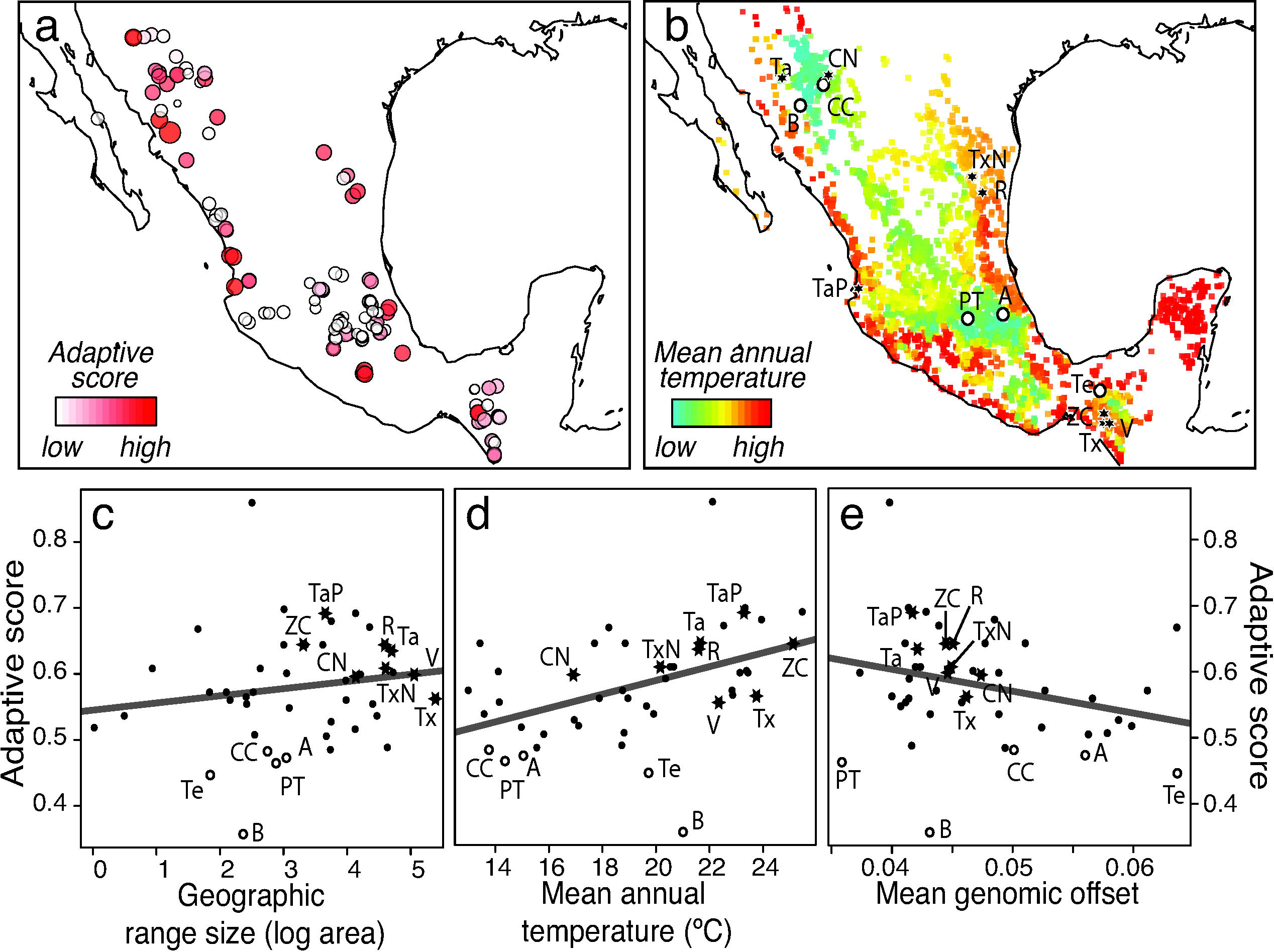
Adaptive scores and genomic offset in climate-related SNPs among landraces of maize accessions in Mexico. **a**, Distribution of adaptive scores for accessions of 46 maize landraces in Mexico estimated from the climate-SNP models for the two teosintes species under the climate change model CCSM_2050_RCP4.5. Circle size is proportional to the estimated adaptive scores, with white circles representing accessions with the adaptive scores. **b**, Known geographic occurrences for the 46 maize landraces in Mexico colored by mean annual temperature. Black stars represent accessions for landraces with high adaptive scores that have been used for maize improvement and white circles represent accessions for restricted landraces with low adaptive potential. **c-e**, Association of adaptive scores versus the geographic range size, mean annual temperature and genomic offset estimated for 46 maize landraces. Regression coefficients for range size: *β* = 0.02, *F* = 1.1, *d.f.* = 45, *p* = 0.31; for temperature: *β* = 0.01, *F* = 17.7, *d.f.* = 45, *p* < 0.001, R^2^ = 0.27; for genomic offset: *β* = −3.31, *F* = 3.76, *d.f.* = 45, *p* = 0.06. ‘Conico Norteño’ (CN), ‘Ratón’ (R), ‘Tabloncillo’ (Ta), ‘Tabloncillo Perla’ (TaP), ‘Tuxpeño’ (Tx), ‘Tuxpeño Norteño’ (TxN), ‘Vandeño’ (V), ‘Zapolote Chico’ (ZC), ‘Arrocillo’ (A), ‘Bofo’ (B), ‘Cristalino de Chihuahua’ (CC), ‘Palomero Toluqueño’ (PT) and ‘Tehua’ (Te).

The statistically significant relationships between paSNPs identified in teosintes and the climatic environment in which maize landraces are grown, could result from past introgression of adaptive genes into maize via gene flow from teosintes^16,18,22,53^. However, hybridization between maize and teosintes appears to be limited to areas of sympatry^22,53^ and thus it seem more likely that the higher adaptive scores for landraces growing in warmer climates reflect past adaptation to these conditions^4^. The low regression coefficients recovered for paSNPs in maize may partially result from the transferability of models of teosintes into maize, or from the quantitative nature of phenotypic traits associated with adaptation to climate^54–57^. Nonetheless, our results emphasize that climate change might give way to future maladaptation of landraces within currently cultivated areas^4^, possibly leading to increased chances of yield declines in many of these landraces under increase temperatures^7^.

Based on our analyses, we identified several landraces (depicted as black stars in fig. 2b-e) with moderate to high adaptive scores and known to be drought resistant^14^ (see SI Appendix, table S9). Indeed, several of these landraces have been used to improve maize lines with enhanced heat and drought resistance, with increased yields^14,52^. We acknowledge that adaptation to climate very likely involves quantitative traits^46,54–57^, yet identifying alleles with small effects on polygenic traits remain a major challenge^57^, especially in species with complex genetic structures, such as teosintes. Nonetheless, the paSNPs and canSNPs we identified are promising candidates to test the response of maize and teosintes to changing climate under experimental settings. The convergence between the adaptive score of landraces, together with historical agronomical practices, opens up the opportunity to fully integrate the vast cultural knowledge on maize landraces with analyses of genomic data in an ecological context.

In an era in which population-level genomic data is more accessible for an increasing number of non-model species, approaches integrating genomic and environmental data are powerful tools to better understand the impacts of climate change on the viability of natural populations^5,30–33^. Predictions of the impact of climate change on species based solely on projected shifts in species’ ranges, although valuable, only provide partial information^28,29^. By building spatiotemporal models of climate-gene relationships, we show that the future of wild populations of teosintes and possibly of several landraces of maize appears to be much harsher than previously estimated^19^.

Although it is tempting to directly relate our results to future declines in population fitness^33^, the association between climate and fitness should be properly validated using experimental or simulation approaches^32,58^. Nevertheless, the fact that future climate change predicts significant alterations to habitat distribution, migration potential, and patterns of local adaptation, raises a red flag for the future of wild and cultivated populations of maize.

Our results represent an important advance moving beyond the standard species distribution models to assess climate change impacts^30–33^, yet our approach still remains an oversimplification of the complex evolutionary and ecological processes affecting populations^31,45,46,57^. Our methodological proposal is a step forward in proving how the analysis of genomic data can identify relevant genetic resources to aid wildlife conservation and crop sustainability under a rapidly changing climate.

## Acknowledgements

We thank Eria A. Rebollar Caudillo, Erika Aguirre Planter and Laura Espinosa Asuar for technical and logistic support during this project. This study was funded by CONACYT Investigación Científica Básica CB2011-167826 and SEP-CONACYT-ANUIES-ECOS France M12-A03-CONACYT-ANUIES 207571 granted to L.E.E; and postdoctoral fellowship from DGAPA-UNAM granted to S.R.-B. This article was written during a sabbatical leave of L.E.E. in the Dept. of Plant and Microbial Biology at the Univ. of Minnesota, with support from the program PASPA-DGAPA, UNAM.

## Author contributions

J.A-L., S.R.-B. and L.E.E. conceived and designed the framework for the analyses. J.A-L. and S.R.-B. collected the data, conducted analyses and drafted the manuscript. L.E.E. secured the funding. All authors discussed results and wrote the manuscript.

## Data availability

Genomic data are available at the Dryad Digital Repository: http://dx.doi.org/10.5061/dryad.8m648, https://doi.org/10.5061/dryad.tf556, and https://doi.org/10.5061/dryad.4t20n. The list of candidate SNPs at https://doi.org/10.1111/mec.14203. Geographic occurrences of teosintes and maize landraces at http://www.biodiversidad.gob.mx. R data and code at https://github.com/spiritu-santi/teosintes.

## References

1. Allendorf FW, Hohenlohe PA, Luikart G. 2010. Genomics and the future of conservation genetics. Nat Rev Genet 11: 697–709.

2. Dai A. 2012. Increasing drought under global warming in observations and models. Nat Clim Change 1633: 52–57.

3. Pauls SU, Nowak C, Bálint M, Pfenninger M. 2012. The impact of global climate change on genetic diversity within populations and species. Mol Ecol 22: 925–946.

4. Hellin J, Bellon MR, Hearne SJ. 2014. Maize landraces and adaptation to climate change in Mexico. J Crop Improv 28: 482–501.

5. Schafer ABA, et al. 2015. Genomics and the challenging translation into conservation practice. Trends Ecol Evol 30: 78–87.

6. Pacifici M, et al. 2017. Species’ traits influenced their response to recent climate change. Nat Clim Change 3223: 1–4.

7. Tigcheelar M, Battisti DS, Naylor RL, Ray DK. 2018. Future warming increases probability of globally synchronized maize production shocks. Proc Natl Acad Sci 115: 6644–6649.

8. Palmgren MG, et al. 2014. Are we ready for back-to-nature crop breeding□? Trends Plant Sci. 20: 155–164.

9. Warschefsky E, Penmetsa RV, Cook DR, Wettberg EJ. 2014. Back to the wilds: Tapping evolutionary adaptations for resilient crops through systematic hybridization with crop wild relatives. Am J Bot 101: 1791–1800.

10. Aguirre-Liguori JA, Aguirre-planter E, Eguiarte LE. 2016. Genetics and ecology of wild and cultivated maize: domestication and introgression. In: Ethnobotany of Mexico. New York: Springer, pp 403–416

11. Zhang H, Mittal N, Leamy LJ, Barazani O, Song BH. 2016. Back into the wild — Apply untapped genetic diversity of wild relatives for crop improvement. Evol Appl 10: 5–24.

12. Dempewolf H, et al. 2017. Past and future use of wild relatives in crop breeding. Crop Sci 57: 1070–1082.

13. Matsuoka Y, et al. 2002. A single domestication for maize shown by multilocusmicrosatellite genotyping. Proc Natl Acad Sci 99: 6080–6084.

14. CONABIO. 2011. Base de datos del proyecto global “Recopilación, generación, actualización y análisis de información acerca de la diversidad genética de maíces y sus parientes silvestres en México”. Comisión Nacional para el Conocimiento y Uso de la Biodiversidad. México, DF.

15. Sánchez GJJ. 2011. Diversidad del maíz y teocintle. Informe preparado para el proyecto global “Recopilación, generación, actualización y análisis de información acerca de la diversidad genética de maíces y sus parientes silvestres en México” de la Comisión Nacional para el Conocimiento y Uso de la Biodiversidad. Comisión Nacional para el Conocimiento y Uso de la Biodiversidad. México, DF.

16. Hufford MB, Martínez-Meyer E, Gaut BS, Eguiarte LE, Tenaillon MI. 2012. Inferences from the historical distribution of wild and domesticated maize provide ecological and evolutionary insight. PLoS ONE 11: e47659.

17. Sánchez González JJ, et al. 2018. Ecogeography of teosinte. PLoS ONE 13: e0192676.

18. van Heerwaarden J, et al. 2011. Genetic signals of origin, spread, and introgression in a large sample of maize landraces. Proc Natl Acad Sci 108: 1088–1092.

19. Ureta C, Martínez-Meyer E, Perales HR, Álvarez-Buylla ER. 2012. Projecting the effects of climate change on the distribution of maize races and their wild relatives in Mexico. Glob Change Biol 18: 1073–108.

20. Baltazar BM, Sanchez-Gonzalez JJ, de la Cruz-Larios L, Schoper J B. 2005. Pollination between maize and teosinte: an important determinant of gene flow in Mexico. Theor Appl Genet 110: 519–526.

21. Kato TAY, Sánchez JJG. 2002. Introgression of chromosome knobs from Zea diploperennis into maize. Maydica 47: 33–50.

22. Warburton ML, et al. 2011. Gene flow among different teosinte taxa and into the domesticated maize gene pool. Genet Resour Crop Ev 58:1243–1261.

23. Wilkes HG. 2006. Urgent notice to all maize researchers: disappearance and extinction of the last wild teosinte populations is more than half completed. A modest proposal for teosinte evolution and conservation in situ: the Balsas, Guerrero, Mexico. Maydica 52:49–58

24. Pyhäjärvi T, Hufford MB, Mezmouk S, Ross-Ibarra J. 2013. Complex patterns of local adaptation in teosinte. Genome Biol Evol 5: 1594–1609.

25. Aguirre-Liguori JA, et al. 2017. Connecting genomic patterns of local adaptation and niche suitability in teosintes. Mol Ecol 26: 4226–4240.

26. Fustier M-A, et al. 2017. Signatures of local adaptation in lowland and highland teosintes from whole genome sequencing of pooled samples. Mol Ecol 26: 2738–D2756.

27. Peterson AT, et al. 2011. Ecological niches and geographic distributions. Princeton University Press, New Jersey.

28. Pagel J, Schurr FM. 2012. Forecasting species ranges by statistical estimation of ecological niches and spatial populations dynamics. Glob Ecol Biogeogr 21: 293–304.

29. Gotelli NJ, Stanton-Geddes J. 2015. Climate change, genetic markers and species distribution modelling. J Biogeogr 42: 1577–1585.

30. Fitzpatrick MC, Keller SR. 2015. Ecological genomics meets community-level modelling of biodiversity: mapping the genomic landscape of current and future environmental adaptation. Ecol Lett 18: 1–16.

31. Peterson ML, Doak DF, Morris WF. 2019. Incorporating local adaptation into forecasts of species’ distribution and abundance under climate change. Global Change Biol early view: https://doi.org/10.1111/gcb.14562.

32. Exposito-Alonso M, et al. 2017. Genomic basis and evolutionary potential for extreme drought adaptation in Arabidopsis thaliana. Nat Ecol Evol 2: 352–358.

33. Bay RA, et al. 2018. Genomic signals of selection predict climate-driven populations declines in a migratory bird. Science 359: 83–86.

34. Phillips SJ, Anderson RP, Schapire RE. 2006. Maximum entropy modeling of species geographic distributions. Ecol Model 190: 231–259.

35. Hijmans RJ, Cameron SE, Parra JL, Jones PG, Jarvis A 2005. Very high-resolution interpolated climate surfaces for global land areas. Int J Climatology, 25: 1965–1978.

36. Villemereuil P, Gaggiotti OE. 2015. A new FST-based method to uncover local adaptation using environmental variables. Methods Ecol Evol 6: 1248–1258.

37. Coop G, Witonsky D, Di Rienzo A, Pritchard JK. 2010. Using environmental correlations to identify loci underlying local adaptation. Genetics 185: 1411–1423.

38. Jombart T. 2008. adegenet: a R package for the multivariate analysis of genetic markers. Bioinformatics 24: 1403–1405.

39. R Core Team. 2018. R: A language and environment for statistical computing. R Foundation for Statistical Computing, Vienna, Austria.

40. Goudet J. 2005. Hierfstat, a package for R to compute and test variance components and F-statistics. Mol Ecol Notes 5: 184–186.

41. Ellis N, Smith SJ, Pitcher CR. 2012. Calculating importance gradients on physical predictors. Ecology 93: 156–168.

42. Hijmans RJ. 2017. raster: Geographic Data Analysis and Modeling. R package version 2.6–7. https://CRAN.R-project.org/package=raster

43. McRae BH, Dickson BG, Keitt T. 2008. Using circuit theory to model connectivity in ecology, evolution, and conservation. Ecology 89: 2712–2724.

44. Arteaga MC, et al. 2016. Genomic variation in recently collected maize landraces from Mexico. Genomics Data 7: 38–45.

45. Hoffmann AA, Sgrò CM. 2011. Climate change and evolutionary adaptation. Nature 470: 479–485.

46. Siol M, Wright SI, Barrett SC. 2010. The population genomics of plant adaptation. New Phytol 188: 313–332.

47. Frankham R. 2010. Challenges and opportunities of genetic approaches to biological conservation. Biol Conserv 143: 1919–1927.

48. Jezkova T, Wiens JJ. 2016. Rates of change in climatic niches in plant and animal populations are much slower than projected climate change. Proc Royal Soc B 283: 20162104.

49. Ackerly DD, et al. 2010. The geography of climate change: implications for conservation biogeography. Divers Distrib 16:476–487.

50. Manel S, Schwartz MK, Luikart G, Taberlet P. 2003. Landscape genetics: combining landscape ecology and population genetics. Trends Ecol Evol 18: 189–197.

51. Loarie SR, et al. 2009. The velocity of climate change. Nature 462: 1052–1055.

52. Wang L, Yang A, He C, Qu M, Zhang J. 2008. Creation of new maize germplasm using alien introgression from Zea mays ssp. mexicana. Euphytica 164: 789–801.

53. Moreno-Letelier A, et al. 2017. Was maize domesticated in the Balsas Basin? Complex patterns of genetic divergence, gene flow and ancestral introgressions among Zea subspecies suggest an alternative scenario. bioRxiv: 239707.

54. Yoder JB, et al. 2014. Genomic signature of adaptation to climate in Medicago truncatula. Genetics 196: 1263–1275.

55. Tiffin P, Ross-Ibarra J. 2014. Advances and limits of using population genetics to understand local adaptation. Trends Ecol Evol 29: 673–680.

56. Kawecki TJ, Ebert D. 2004. Conceptual issues in local adaptation. Ecol Lett 7: 1225–1241.

57. Yeaman S. 2015. Local adaptation by alleles of small effect. Am Nat 186: S74–S89.

58. Fitzpatrick MC, Keller SR, Lotterhos KE. 2018. Comment on “Genomic signals of selection predict climate-driven population declines in a migratory bird”. Science 361: eaat7279.

