## Supplementary material for "Climate change is predicted to disrupt patterns of local adaptation in wild and cultivated maize": SI Appendix

**This PDF file includes:**

Supplementary Methods

Supplementary References

Supplementary Figures 1 to 12

Supplementary Tables 1 to 9

**Supplementary Methods**

***Genomic dataset***

We combined the SNPs datasets of ref.[1] and ref.[2] to obtain a dataset of 33,454 SNPs distributed across all chromosomes of maize. Genotyping quality was set with a threshold of 0.15 for the $GC_{50}$ score and by removing monomorphic SNPs and clusters with call rates below 0.85. See ref.[1] and [2] for more details.

*Defining putative adaptive SNPs (paSNPs).* The dataset including the locally adapted SNPs identified by ref.[2] was subdivided based on the power of individual SNPs to discriminate among climatically defined groups of populations. Based on the values of the first environmental principal component, associated with temperature[2], we categorized populations for each species into four groups based on the quartile distribution of temperature values across sampled populations: Group 1-cold (6 *parviglumis* and 7 *mexicana* populations); Group 2-normal cold (6 *parviglumis* and 6 *mexicana* populations); Group 3-normal warm (6 *parviglumis* and 6 *mexicana* populations); and Group 4-warm (6 *parviglumis* and 6 *mexicana* populations).

We tested the power of locally adapted SNPs to discriminate among the above-mentioned groups of populations by performing a Discriminant Analyses on Principal Components (DAPC)[3] for each teosintes species separately using ten principal components (summarizing 25% of the genetic variability across populations) and retaining the first four discriminant functions. These first four discriminant functions explained ~ 67% of the among-population environmental variance for *parviglumis* and ~ 77% for *mexicana.* We plotted the contribution of individual SNPs to the discriminant functions using the *loadingplot* function of the *adegenet* package[3] in R[4] with the default parameters. We selected the locally adapted SNPs with the highest discrimination power (paSNPs) using the default parameters (*i.e.*, the third quartile of the distribution of loadings (fig. S1).

***Predictive models using Gradient Forest***

Gradient Forest (GF) uses a machine-learning algorithm to divide biological data into bins (allele frequencies) occurring at numerous split values along a given environmental gradient[5]. Moving throughout an environmental gradient, the algorithm estimates the amount of variation in the data (in this case allele frequencies) explained by the different split values (split importance). In principle, the estimation of split importance along a gradient does not depend on the initial allele frequency in any given population, but on the amount of change observed among populations located along different regions of the gradient. GF cumulatively sums the split importance along a gradient, which then reflects the overall association of an environmental gradient with allele frequency changes, using this information to construct allele turnover functions[6] along the environmental gradient. Given the correspondence between environmental and geographic space[7], these functions can be directly projected into geographic space, allowing the visualization of differences in genetic composition throughout the species' distribution ranges, even in places were no populations have been sampled.

To estimate the genomic offset under different future scenarios of climate change, Euclidian distances between present and future models are used to identify regions with a more pronounced disruptive effect of climate change, where larger distances are indicative of higher populations vulnerability[6]. More specifically, the Euclidean distances are estimated as follows,

$$\sqrt{\sum_{i}^{n}(present_i - future_i)^2}$$

where *n* represents the number of bioclimatic variables used in the analyses, and *i* represents the contribution of a grid (coordinate) to the allelic turnover function for a given environment.

For each subspecies and each set of SNPs (paSNPs, canSNPs and refSNPs), we estimated the allelic turnover models using the *gradientforest*[5] package in R[4]. These allelic turnover models were used to predict the genomic offset across the teosintes distribution. Values of genomic offset were standardized relative to the maximum observed Euclidian distance[6] estimated across known teosintes occurrences[8], setting areas with a standardized value > 1 to NA. The genomic offset values across the distribution of teosintes were transformed into a raster grid and the estimated values of genomic offset were extracted for all known teosintes occurrences[8] and sampled populations using the *raster* package[9] in R[4].

In addition to the GF analyses using the three sets of SNPs, we estimated allele turnover functions using the complete set of 33,454 SNPs identified in teosintes[1,2]. Due to computational limitation, we only estimated allele turnover and not genomic offset for the complete set of SNPs and classified them based on their overall contribution to the model (i.e., gradient forest's $R^2$). Based on the individual contribution of each SNP to the model ($R^2$), we assessed whether the canSNPs and paSNPs are a representative sample of SNP-climate association across the genome. For SNPs with significant contribution to the turnover model we used the quartile distribution to categorize SNPs as having low, moderate, high, and very high climate-frequency associations. We compared the contribution of different SNPs categories to the contribution estimated by the GF analyses on the canSNPs and paSNPs, showing that these two sets of SNPs are representative of strong genome-wide climate-frequency associations (figs. S3, S4).

*Validation of allele turnover models*. GF analyses can be sensitive to the presence of habitat heterogeneity masking the presence of climate-frequency associations. In order to test the predictive ability of the resulting allele turnover models, we used simple Mantel correlations to test for significant associations between predicted allele turnover and pairwise genetic differentiation between populations ($F_{ST}$). We also tested for the correlation between allele turnover and environmental distances among populations.

We used the *bedassle*[10] package in R[4] to estimate the pairwise genetic differentiation ($F_{ST}$) between populations for paSNPs, canSNPs and refSNPs, separately. To estimate environmental differentiation, we extracted environmental information of the 19 bioclimatic layers[11] used in the species distribution models for each pixel predicted by the species distribution models. We then performed a principal component analyses (PCA) on these data to reduce environmental variation down to six principal components and obtain values for each sampled population of teosintes. We used the data for the six principal components to calculate Euclidian distances between populations (environmental differentiation) using the *stats* package in R[4]. Finally, for each GF model (e.g., paSNPs, canSNPs, refSNPs) we modified the original script of ref.[6] to perform a PCA and reduce the contribution of the 19 bioclimatic variables to the allelic turnover function into six principal components. These principal components were used to estimate Euclidian distances between populations, which reflect the predicted allelic dissimilarity between populations (predicted genetic differentiation).

For each SNPs set, we evaluate the correlation between: (1) predicted genetic dissimilarity and genetic differentiation; and (2) predicted genetic dissimilarities and environmental differentiation. For both teosintes species we found strong associations (all *p-values* $< 0.05$) between predicted genetic dissimilarity and environmental differentiation between populations (fig. S10). In addition, we found strong significant associations between genetic differentiation and predicted genetic dissimilarities for paSNPs and canSNPs, after controlling for environmental distances. We observed a low, yet significant association between genetic differentiation and predicted genetic dissimilarities for refSNPs (fig. S10).

The lower power of GF to predict pairwise genetic differentiation using refSNPs is expected because these SNPs have an overall low contribution to the model (fig. S3). Accordingly, we found that allele turnover functions have a stronger predictive power in *mexicana* than in *parviglumis*, the latter having a higher environmental variation and a stronger genetic structure than the former[1,2]. This indicates that habitat heterogeneity can have significant impacts on the results of GF. However, in the case of teosintes the allele turnover functions for canSNPs and paSNPs were robust to the effects of habitat heterogeneity.

*Allele frequency differences among SNP sets.* The allelic turnover models show that paSNPs and canSNPs have significantly higher contribution to model construction than refSNPs (fig. S3). Although this is expected and reflects the stronger frequency-environment associations (i.e., local adaptation) seen for paSNPs and canSNPs, these differences can be biased by significant discrepancies in allele frequencies for refSNPs compared to locally adapted SNPs. In the present case, paSNPs and canSNPs were defined using methods that rely on identifying outlier $F_{ST}$ values among SNPs[2]. If higher predictability of allelic turnover is affected by the stronger differences in allele frequencies and not local adaptation, then using refSNPs with marked differences in allelic frequencies among populations should generate significantly higher allelic turnover models.

To assess the impacts of allele frequencies on allele turnover models using refSNPs, we performed additional GF analyses using a set of refSNPs for which the among-population frequencies more closely matched those observed for canSNPs and paSNPs. For this, we estimated the standard deviation of allele frequencies among populations for the canSNPs and paSNPs. Then, we selected the set of refSNPs (hereafter n_refSNPs) showing a standard deviation higher than the median standard deviation observed for canSNPs (very few SNPs had standard deviation within the range observed for paSNPs) and showing no significant associations with environmental variables.

We performed a GF analysis using the paSNPs, canSNPs, refSNPs, and n_refSNPs to assess the contribution of each SNP to the turnover function. For both teosintes species we found that the n_refSNPs had a lower predictability than paSNPs. However, we found that the model contribution of n_refSNPs was similar to that estimated for refSNPs only in *mexicana*, whereas for *parviglumis* there was a greater variance in the predictive power of n_refSNPs with some overlap with canSNPs (fig. S3). Although this might indicate a possible bias in the signal recovered for canSNPs due to genetic structure in *parviglumis*, it is important to recall that canSNPs were selected by employing outlier tests (*e.g.*, *bayescenv*, *bayenv*) that controlled for possible gene surfing[2]. The estimated allele turnover for paSNPs appears to be robust to differences in among-population allele frequencies in both species.

*Identification of candidate SNPs.* Another important source of bias for the modeling of allele turnover is the proper identification of genes responsible for local adaptation[12]. In this context, modeling and interpreting the results for only a handful of SNPs (e.g., paSNPs) may be an over-simplification of the genomic processes affecting local adaptation to climate[13]. Thus, we assessed the impact of including varying number of SNPs for the construction of allele turnover models and the estimation of genomic offset under future climate change scenarios. For this, we performed GF analyses to estimate the genomic offset based on different sets of outlier SNPs identified by ref.[2]: (1) canSNPs + paSNPs (USNPs); (2) outlier SNPs detected by bayescenv (scenvSNPs); (3) outlier SNPs detected by bayenv (baySNPs); and (4) outlier SNPs detected by bayenv and bayescenv (bay∩scenvSNPs). For details on the procedures for outlier detection see ref.[2].

Overall, these analyses show that many outlier SNPs can be informative about allele turnover and thus have elevated contribution to the genomic offset of populations (fig. S3). For instance, as expected the scenvSNPs and baySNPs showed a lower genomic offset than that observed for paSNPs, yet the estimated values are greater than those estimated for refSNPs. These results highlight two important points. First, the method to identify outlier SNPs is crucially important for the estimation of genomic offset[12], particularly in situation where populations have pronounced patterns of genetic and environmental differentiation, such as *parviglumis*. Second, the estimated genomic offset for populations using a handful of locally adapted SNPs appears to be robust to the number of SNPs included in the model (fig. S3, S4). Increasing the number of SNPs did not produce significantly distinct patterns of estimated genomic offset than those observed using the reduced set of canSNPs and paSNPs. Although there is variation in the range of estimated genomic offset among different sets of SNPs (fig. S3), expected due to varying levels of SNP-climate associations (*i.e.*, allele effects), we found that the per-population genomic offset was highly correlated among all outlier SNP sets (fig. S3i). This suggest that unseen candidate SNPs or new candidate SNPs entering a population (either by migration, mutation or pre-adaptation) will most probably have small impact on the observed patterns of local adaptation and genomic offset estimated across populations (information on individual SNPs can be found at https://github.com/spiritu-santi/teosintes).

*Initial allele frequencies and climate change.* To gauge the influence of initial allele frequencies on the resulting levels of genomic offset, we tested the correlation between the genomic offset estimated for each population and the frequency of putative adaptive alleles at paSNPs within populations. Since we are interested in the response of populations to increasing temperature, we used the frequencies of the warm-adapted alleles at paSNPs, identified as the alleles with higher frequencies in populations growing at the warm-end of the species' climatic niche (climatic groups 3 and 4, see above). For simplicity, we estimated an adaptive score for each population by summing up adaptive allele frequencies over all paSNPs (see below) and tested the relationship with genomic offset only under the two most extreme models: CCSM 4.5 2050 and CCSM 8.5 2070 (fig. S5).

We also tested the correlation between genomic offset and the estimated change in climatic conditions expected for each population, which were measured as the sum of the absolute differences between the present and future values for the 19 bioclimatic variables[11] used for the species distribution modeling and GF analyses. We employed simple linear regressions using the *stats* package in R[4] (fig. S5).

#### *Ecological niche modeling*

*Species distribution models.* We used 254 (*mexicana*) and 329 (*parviglumis*) occurrence data points (available at: www.biodiversidad.gob.mx/genes/proyectoMaices.html) and performed the modeling using previously described settings and validation procedures[14]. The resulting species models showed a good performance, with AUC values of 0.982 and 0.972 for *mexicana* and *parviglumis*, respectively.

To generate binary presence/absence maps for the present and future, we applied a minimum presence logistic threshold (*i.e.*, the minimum model value observed among species occurrences) to the models. The change in the predicted geographic distribution of the species was simply estimated as the number of grid-cells predicted in the future relative to the present (models available at https://github.com/spiritu-santi/teosintes). We also estimated overlap between present and future models using the corresponding binary maps.

Given that the sum of two binary maps (presence = 1, absence = 0) does not distinguish between present-only and future-only regions (both with values of 1 in the overlap map), we arbitrarily set a presence value of two for the present models and a value of four for the future models. These values were treated as categorical variables. We summed the values of the two models and defined three sets of regions: (1) regions uniquely predicted for the present (present-only grid-cells, value of two); (2) regions uniquely predicted for the future (future-only grid-cells, value of four); (3) regions of overlap between the two time models (overlap grid-cells, value of six).

*Allele distribution models.* We predicted the geographic distribution for putative adaptive alleles segregating at paSNPs using Maxent v.3.3.3[15]. Since we were interested in modeling the distribution of the putatively warm-adapted alleles, we used as input the populations' geographic coordinates where these alleles were present. Populations that were fixed with the non-adaptive allele did not contribute to the models. We used the same climatic variables; setting and validation procedures used for the species distribution modeling (see above). Overall, the resulting models for the paSNPS showed a good performance, with the lowest AUC values being 0.972 and 0.976 for *mexicana* and *parviglumis*, respectively.

For each adaptive allele distribution model (8 for *mexicana* and 9 for *parviglumis*) we created binary presence/absence maps as depicted above for the species distribution models. Then, for each grid-cell we estimated the sum across allele models constructed for the present-day and under each of the eight future climate change scenarios. We transformed the resulting *raster* grids into binary maps by setting areas with five or more alleles present to 1 and areas with fewer than five alleles to 0.

For each overlapped map, we extracted values for the bioclimatic variables for every grid-cell and generated a data frame combining the climatic conditions for areas predicted in present and future. We used these data to perform a Principal Component Analysis (PCA) with the *prcomp* function in R[4], retaining the first four principal components that explained more than 85% of the total variance in climatic variables.

Because we were interested in obtaining the future distribution of warm-adapted alleles, we estimated the present-day PC climatic range for the normal-warm and warm populations for each subspecies separately, essentially obtaining the current environmental range defining the warm-end niche of the two teosintes (based in the range estimated from the first principal components). Subsequently, we selected areas within the present and future predicted distributions that had climatic values within the present-day environmental range. Basically, this allowed us to define the geographic regions that presently lay within the environmental range of warm-adapted populations and those that will show the same range in future scenarios.

Based on the genomic offset of populations, we cross validated the final adaptive allele distribution models using the Gradient Forests models described above, basically inspecting the genomic offset of populations estimated with Gradient Forests for the three sets of regions (i.e., present-only, future-only, overlap). These analyses show that in general areas laying outside the predicted distribution for paSNPs have a higher mean and variance for genomic offset than areas predicted by either future or present paSNPs models, particularly in *mexicana* (fig. S6).

*Maxent's suitability and allele frequencies.* For each paSNPs and canSNPs, we tested for the correlation between the mean allele frequencies in each population and the mean suitability estimated by Maxent (*e.g.,* logistic output) across models for individual SNPs. Overall, we found that suitability values are not correlated with mean allele frequencies (fig. S11), which is consistent with previous analyses showing the inability of Maxent to predict species abundance[16]. Although some individual paSNPs showed significant associations with suitability, particularly in *parviglumis*, these were non-significant after correcting for multiple comparisons using the Bonferroni test. All analyses were performed using the *raster*[9] and *stats* packages in R[4].

#### Migration analyses

We first identified areas with climatic conditions outside the present environmental range estimated for populations (see above) and treated these areas as hard barriers to migration by setting their value to NA. This assumes that teosintes populations are not capable of establishing in regions outside particular climatic conditions, namely regions with higher temperatures than currently inhabited.

We defined routes of potential migration by identifying regions predicted in the overlapped present and future distribution models for paSNPs, corresponding to the transition surfaces used in circuit theory[17]. These surfaces are built by assigning different resistance values to migration (i.e., environmental costs) to distinct environmental elements[18]. In the present case, different resistance values were assigned across the landscape based on the predicted distribution of paSNPs. Then, we defined areas of potential settlement corresponding to sites where populations could eventually establish in the future, by identifying areas predicted only in the overlapped future models for paSNPs, corresponding to the geographic coordinates into which potential migration would be favored by local adaptation.

We used the *Gdistance* package[19] in R[4] to build transition layers around each focal population, defining a transition function of 1/mean(x) for each pixel (*i.e.*, grid-cell) between the eight neighboring grid-cells (*i.e.*, rooks move). Then we used the geographic coordinates of potential settlement sites and implemented a cost-distance function to determine the resistance values (environmental distances) from the focal population to all sites, as a proxy of migration limitation. To estimate the overall migration limitation of every population, we estimated the overall distribution of environmental distances across all potential settlement sites. We then identified settlement sites with estimated resistance values below the 10, 20, 30, 40 and 50 percentiles of the distribution of environmental distances. These percentiles can be interpreted as the overall resistance to the successful migration into 10, 20, 30, 40, and 50% of the future areas of potential settlement. We identified populations with the least probabilities of migrating (migration unlikely) as those with undefined environmental costs at the 10-precentile thresholds, namely those populations for which the transition matrices become infinity due to the presence of hard barriers to migration. All other populations with positive environmental costs were treated as having high chances of migrating (migration likely) into suitable areas in the future (results of migration analyses under all climate change models are available at https://github.com/spiritu-santi/teosintes).

*Migration potential.* Ultimately, the likelihood of successful migration of a particular population would be proportional, not only to the estimated migration costs, but also to the initial frequencies of adaptive alleles present in the population. Thus, for each population we evaluated the combined frequencies of adaptive alleles at paSNPs, by estimating an adaptive score defined as the sum of individual allele frequencies weighted by the total number of adaptive alleles present in a population. This adaptive score ranges between 0 and 1, with higher values indicating a higher frequency of adaptive alleles. We tested for differences in adaptive score between populations with likely and unlikely migration using an

Analysis of Variance (ANOVA) as implemented in the *stats* package in R[4], but only under the two most extreme climate change models: CCSM_2050_RCP4.5 and CCSM_2070_RCP8.5. In general, populations predicted to have increased migration potential have significantly higher frequencies of adaptive alleles (measured by the adaptive score) than population predicted with low chances of migrating (fig. S8). Accordingly, the estimated genomic offset was higher for populations without migration potential than for those with migration potential, a pattern that is more pronounced in *mexicana* than in *parviglumis* (fig. S8).

Besides the possibility of tracking climate change by migrating into new locations, populations can evolve to respond to future climate conditions by receiving adaptive variants via gene flow from other populations. For *Zea mays* in particular, pollen is wind-dispersed[20], yet studies have shown that the viability of pollen decreases significantly over short scales[21-23]. To assess the potential contribution of adaptive introgression to the response of teosintes populations to climate change, we used the geographic distances and allele frequencies as proxies for the potential for gene flow among populations[24]. These analyses do not consider the presence of known teosintes populations that remain unsampled and with no genetic data[8,25], which may otherwise serve as sources of adaptive alleles or bridges to gene flow between sampled populations.

Basically, for each population we estimated the minimum distances to populations that could serve as sources of adaptive alleles through gene flow. In any given population, incoming gene flow can only increase the frequency of an adaptive allele if the initial frequency of that allele in the source population is greater than in the target population. Thus, we calculated the minimum distance to a population with an adaptive score greater than 0.5 and higher than the adaptive score of the focal population. All else being equal, as geographic distance between two populations increases, the probabilities for gene flow between these populations would decrease. In addition, for each population we identified the set of neighboring populations (arbitrarily set to four populations, corresponding to ~15% of populations) and then evaluated whether these had a higher adaptive score than the focal population. We used ANOVA tests as implemented in the *'stats'* package in R[4] to determine whether geographic distances differed between populations identified as having migration potential and those without migration potential.

We estimated the probability of population migration and gene flow under the two more extreme climate change models (*i.e.,* CCSM-RCP4.5-2005, CCSM-RCP8.5-2070); the results of all tests are summarized in table S7. Populations that are predicted to have increased chances of migrating also appear to be closer to populations that could serve as sources of adaptive alleles via gene flow, than populations with low chances of migrating (fig. S12).

#### *Identifying putative adaptive SNPs in maize landraces*

The MaizeSNP50 BeadChip used by ref.[1] and ref.[2] to genotype teosintes, was originally designed to detect genetic diversity among maize landraces. We explored whether the paSNPs identified for teosintes have been documented in particular landraces of maize[26]. For this, we downloaded the genotype data for maize[26], consisting of 36,931 SNPs across 46 landraces in Mexico, with 1 to 16 accessions per landrace.

Briefly, we obtained the adaptive score for each landrace by adding the frequency of the putative adaptive alleles segregating at paSNPs recorded in each accession, *i.e.*, 1.0 for the adapted homozygous genotype, 0.5 for a heterozygous genotype, and 0.0 for the non-adapted homozygous genotype. The 'adaptive' score relates to the frequency of adaptive alleles, where a lower score indicates that adaptive alleles tend to be less represented in a particular landrace. A higher adaptive score would suggest that adaptive alleles are near fixation in particular landraces, probably resulting from past adaptation to warm and dry environments or the probable introgression of adaptive alleles from teosintes[27]. On the contrary, lower adaptive scores indicate that landraces have the potential to adapt to changing environmental conditions, with the maximum adaptive potential at intermediate allele frequencies. In principle, the adaptive score is not related to the genetic diversity within a population, because both homozygous states have the same genetic diversity but contrasting adaptive potential scores (i.e., genetic diversity would be greatest at intermediate adaptive potential scores), whereas the heterozygous state has the highest genetic diversity, but intermediate adaptive potential scores.

We acknowledge two limitations of our approach to identifying adaptive alleles in maize landraces. First, although gene-climate associations can be analyzed using single individuals per population[28], the direct application in maize of ecological models using allele frequency data, such as Gradient Forests, is prevented because the available SNPs dataset for maize is limited to one or a few accessions per landrace. Accordingly, we might expect a non-negligible number of unseen adaptive alleles across landraces purely due to sampling artifacts. Second, extrapolating from wild relatives to maize is not straightforward because the latter are highly dependent on humans for reproduction and usually grow under conditions that act as possible buffers against local- and regional-level climatic trends[29]. Nevertheless, ecological analyses indicate similar trends of shrinking habitat in both teosintes and maize landraces in response to rising temperature and increasing aridity[30].

For each landrace of maize, we approximated the geographic range by estimating a spatial convex hull distribution model[31] around the reported occurrences[8] as implemented in the *dismo* package[32] in R[4]. Unlike species distribution modeling, the convex hull estimates species geographic ranges using spatial data alone and can be easily implemented for multiple species[31]. We chose to use convex hulls to estimate the geographic range of maize landraces, because these are easy to apply and are independent from environmental data. Basically, a convex hull model predicts that a species (landrace) is present inside the convex hull of a set of geographic occurrences and absent outside this area, yet the model assumes that a species is present throughout the entire hull[31].

### SI References


1. Pyhäjärvi T, Hufford MB, Mezmouk S, Ross-Ibarra J. 2013. Complex patterns of local adaptation in teosinte. *Genome Biol Evol* **5:** 1594-1609.
2. Aguirre-Liguori JA, et al. 2017. Connecting genomic patterns of local adaptation and niche suitability in teosintes. *Mol Ecol* 26: 4226-4240.
3. Jombart T. 2008. *adegenet*: a R package for the multivariate analysis of genetic markers. *Bioinformatics* 24: 1403-1405.
4. R Core Team. 2018. R: A language and environment for statistical computing. R Foundation for Statistical Computing, Vienna, Austria.
5. Ellis N, Smith SJ, Pitcher CR. 2012. Calculating importance gradients on physical predictors. *Ecology* 93: 156-168.
6. Fitzpatrick MC, Keller SR. 2015. Ecological genomics meets community-level modelling of biodiversity: mapping the genomic landscape of current and future environmental adaptation. *Ecol Lett* **18:** 1-16.
7. Peterson AT, et al. 2011. *Ecological niches and geographic distributions*. Princeton University Press, New Jersey.
8. CONABIO. 2011. *Base de datos del proyecto global "Recopilación, generación, actualización y análisis de información acerca de la diversidad*

*genética de maíces y sus parientes silvestres en México"*. Comisión Nacional para el Conocimiento y Uso de la Biodiversidad. México, DF.

**9.** Hijmans RJ. 2017. *raster*: Geographic Data Analysis and Modeling. R package version 2.6-7. https://CRAN.R-project.org/package=raster

**10.** Bradburd GS, Ralph PL, Coop GM. 2013. Disentangling the effects of geographic and ecological isolation on genetic differentiation. *Evolution* **67**: 3258-3273.

**11.** Hijmans RJ, Cameron SE, Parra JL, Jones PG, Jarvis A 2005. Very high-resolution interpolated climate surfaces for global land areas. *Int J Climatology*, **25**: 1965–1978.

**12.** Fitzpatrick MC, Keller SR, Lotterhos KE. 2018. Comment on "Genomic signals of selection predict climate-driven population declines in a migratory bird". *Science* 361: eaat7279.

**13.** Kawecki TJ, Ebert D. 2004. Conceptual issues in local adaptation. *Ecol Lett* 7: 1225-1241.

**14.** Hufford MB, Martínez-Meyer E, Gaut BS, Eguiarte LE, Tenaillon MI. 2012. Inferences from the historical distribution of wild and domesticated maize provide ecological and evolutionary insight. *PLoS ONE* 11: e47659.

**15.** Phillips SJ, Anderson RP, Schapire RE. 2006. Maximum entropy modeling of species geographic distributions. *Ecol Model* **190:** 231-259.

**16.** Yañes-Arenas C, Martinez-Meyer E, Mandujano S, Rojas-Soto O. 2012. Modelling geographic patterns of population density of the white-tailed deer in central Mexico by implementing ecological niche theory. *Oikos* 121: 2081-2089.

**17.** McRae BH, Dickson BG, Keitt T. 2008. Using circuit theory to model connectivity in ecology, evolution, and conservation. *Ecology* **89:** 2712-2724.

**18.** Manel S, Schwartz MK, Luikart G, Taberlet P. 2003. Landscape genetics: combining landscape ecology and population genetics. *Trends Ecol Evol* **18:** 189-197.

**19.** van Etten J. 2017. R Package *gdistance*: Distances and routes on geographical grids*. J Stat Softw* **76:** 1-21.

**20.** Aguirre-Liguori JA, Aguirre-planter E, Eguiarte LE. 2016. *Genetics and ecology of wild and cultivated maize: domestication and introgression. In*: Ethnobotany of Mexico. New York: Springer, pp 403-416

**21.** Luna VS, et al. 2001. Maize pollen longevity and distance isolation requirements for effective pollen control. *Crop Sci* **41**: 1551–1557.

**22.** Aylor DE, Baltazar BM, Schoper JB. 2005. Some physical properties of teosinte (*Zea mays* subsp. *parviglumis*) pollen. *J Exp Bot* 56; 2401-2407.

**23.** Hufford MB, Gepts P, Ross-Ibarra. 2011. Influence of cryptic population structure on observed mating patterns in the wild progenitor of maize (*Zea mays* ssp. *parviglumis*). *Mol Ecol* **20**: 46-55.

**24.** Slatkin M. 1985. Gene flow in natural populations. *Ann Rev Ecol Syst* **16**: 393-430.

**25.** Sánchez González JJ, et al. 2018. Ecogeography of teosinte. *PLoS ONE* 13: e0192676.

**26.** Arteaga MC, et al. 2016. Genomic variation in recently collected maize landraces from Mexico. *Genomics Data* **7:** 38-45.

**27.** Hufford MB, et al. 2013. The genomic signature of crop-wild introgression in maize. *PLoS Genet* **9**: e1003477.

**28.** Yoder JB, et al. 2014. Genomic signature of adaptation to climate in *Medicago truncatula*. *Genetics* **196**: 1263-1275.

**29.** Sánchez GJJ. 2011. *Diversidad del maíz y teocintle.* Informe preparado para el proyecto global "Recopilación, generación, actualización y análisis de información acerca de la diversidad genética de maíces y sus parientes silvestres en México" de la Comisión Nacional para el Conocimiento y Uso de la Biodiversidad. Comisión Nacional para el Conocimiento y Uso de la Biodiversidad. México, DF.

**30.** Ureta C, Martínez-Meyer E, Perales HR, Álvarez-Buylla ER. 2012. Projecting the effects of climate change on the distribution of maize races and their wild relatives in Mexico. *Glob Change Biol* 18: 1073-108.

**31.** Meyer L, Diniz-Filho JAF, Lohmann L. 2018. A comparison of the hull methods for estimating species ranges and richness maps. *Plant Ecol Div*, **DOI: 10.1080/17550874.2018.1425505**

**32.** Hijmans RJ, Phillips S, Leathwick J, Elith J. 2017. *dismo*: Species distribution modeling. R package version 1.1-4. https://CRAN.R-project.org/package=dismo

**Supplementary Figure 1.** Diffrent candidate SNPs have varying power to discriminate among climatically defined groups of sampled populations of two species of teosintes in Mexico (*Zea mays* spp. *parviglumis* and *Z. mays* spp. *mexicana*). Bars represent the loadings for the first discriminant function of the Discriminant Analysis of Principal Components (DAPC) estimated for candidate SNPs. Red bars represent the loadings for the putative adaptive SNPs selected for each species.

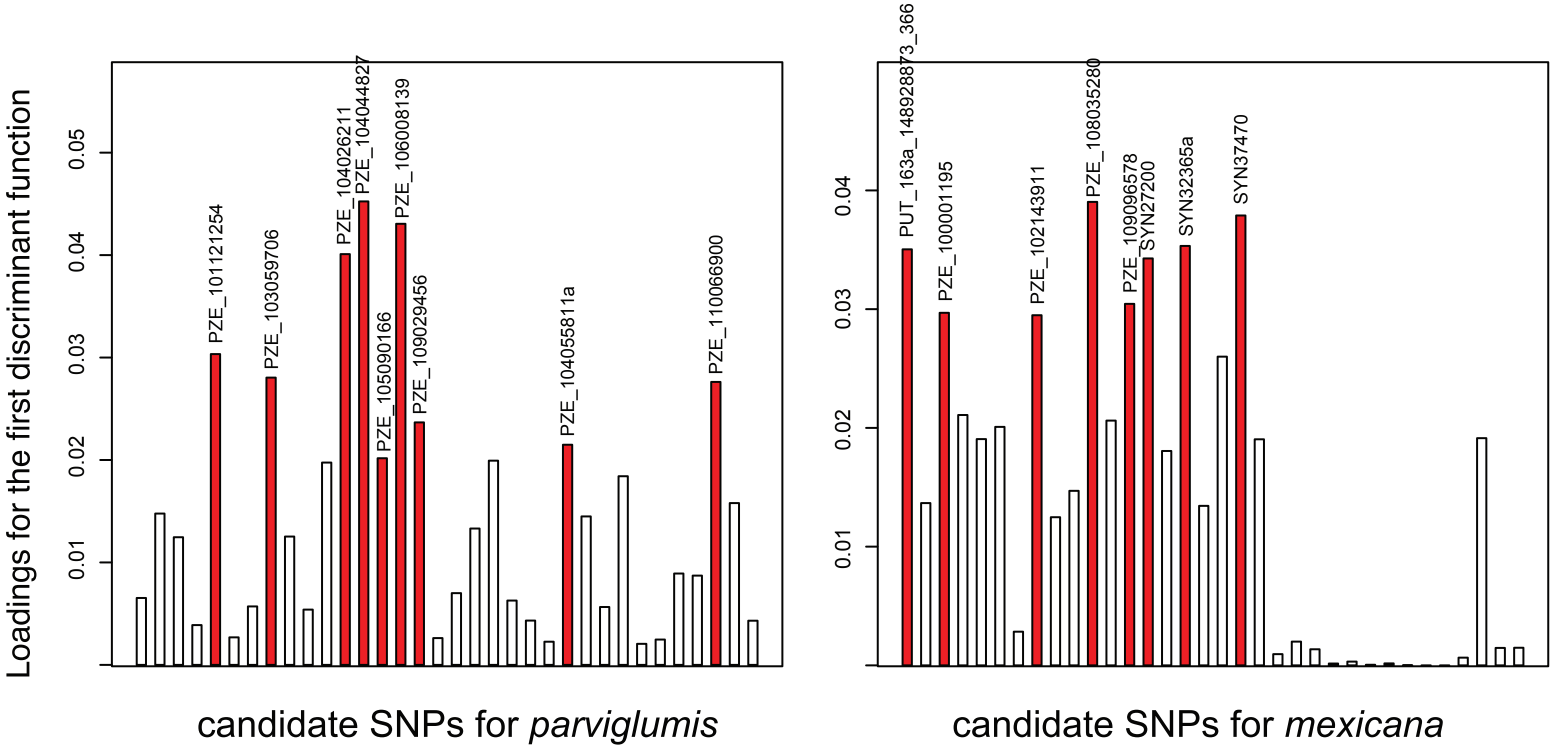

**Supplementary Figure 2.** Geographic distribution of two species of teosintes in Mexico: (*Zea mays* spp. *parviglumis* and *Z. mays* spp. *mexicana*) as predicted by ecological niche modeling. Overlap between the present-day and future distirbution of teosintes under the eight models of climate change. Ecological niche models were constructed using all known ocurrence records for the two teosinte species in Mexico. Distribution areas predicted only in the present-day are shown in light blue, areas predicted only in future models are shown in yellow, and areas predicted to overlap by present and future models ares shown in black (models available in ASCII format at https://github.com/spiritu-santi/teosintes).

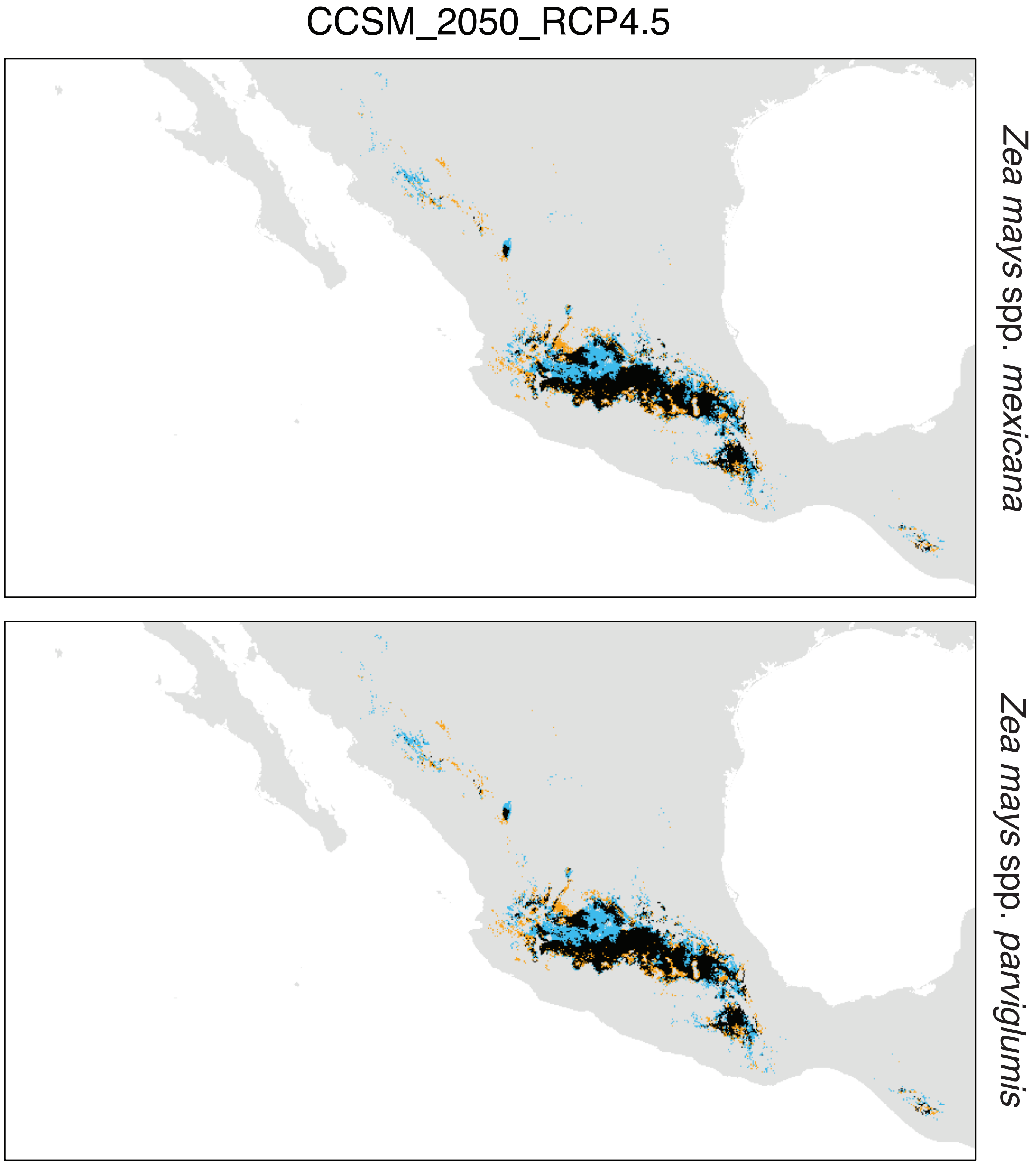

**Supplementary Figure 2.** Continued

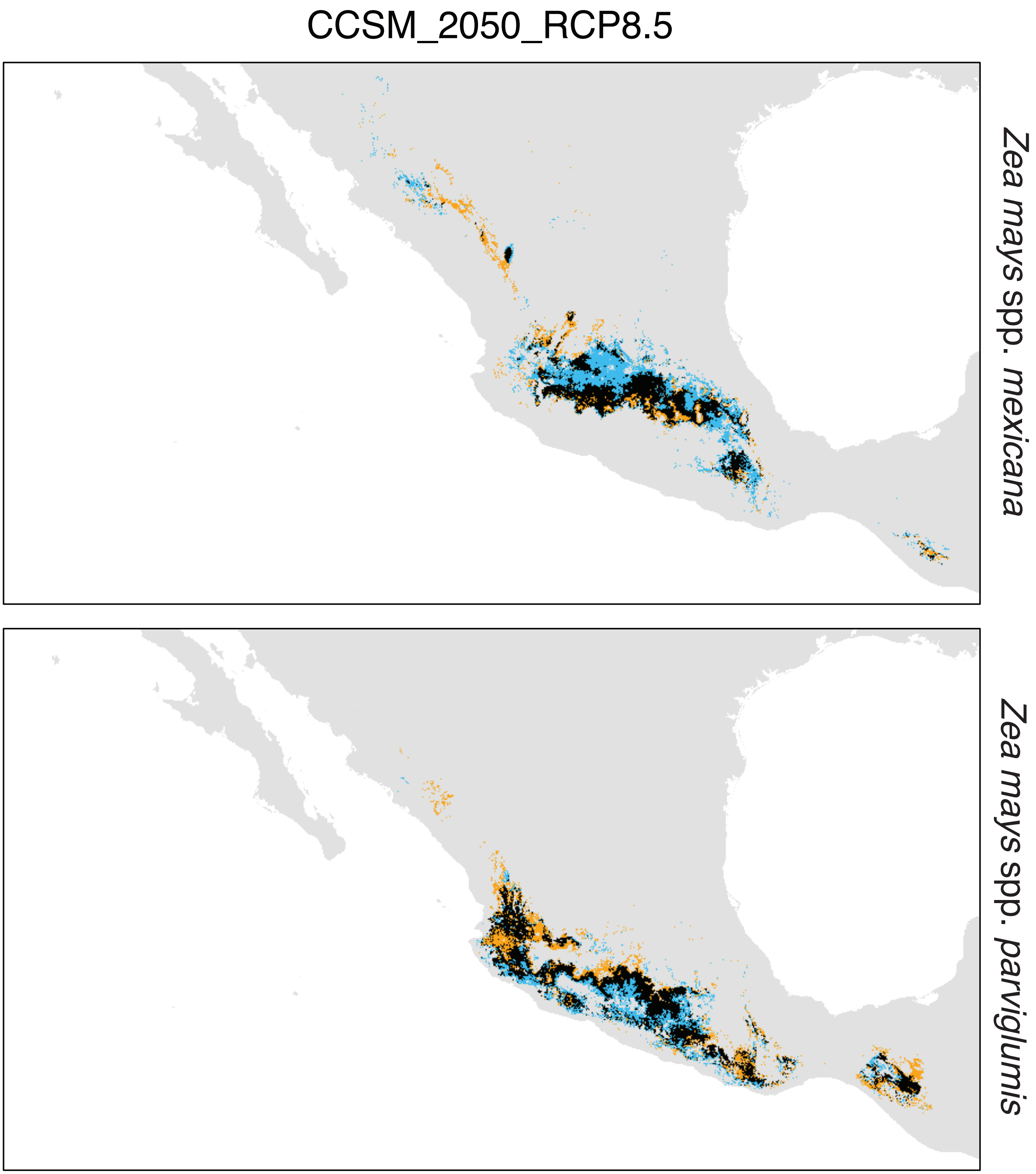

**Supplementary Figure 2.** Continued

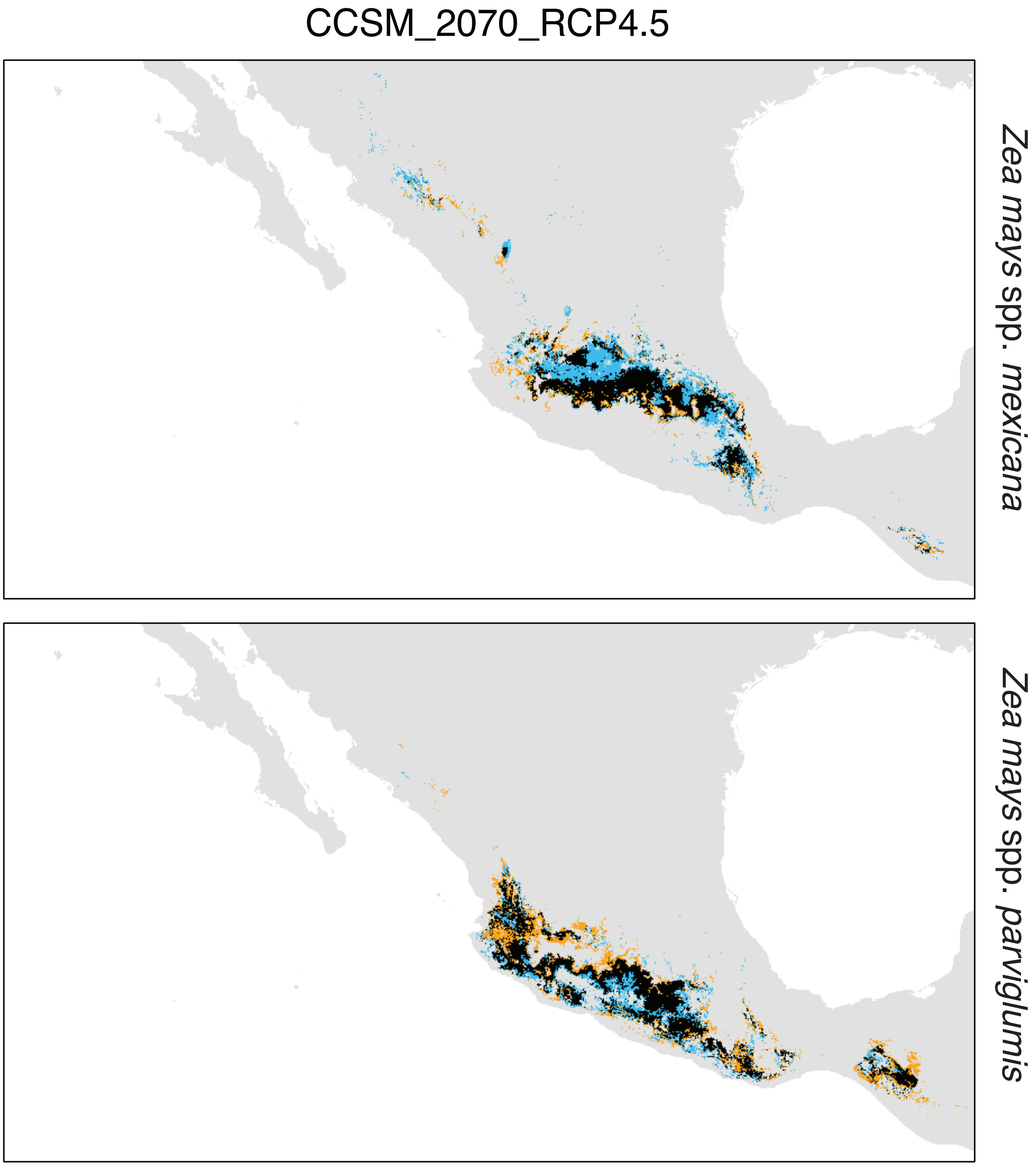

**Supplementary Figure 2.** Continued

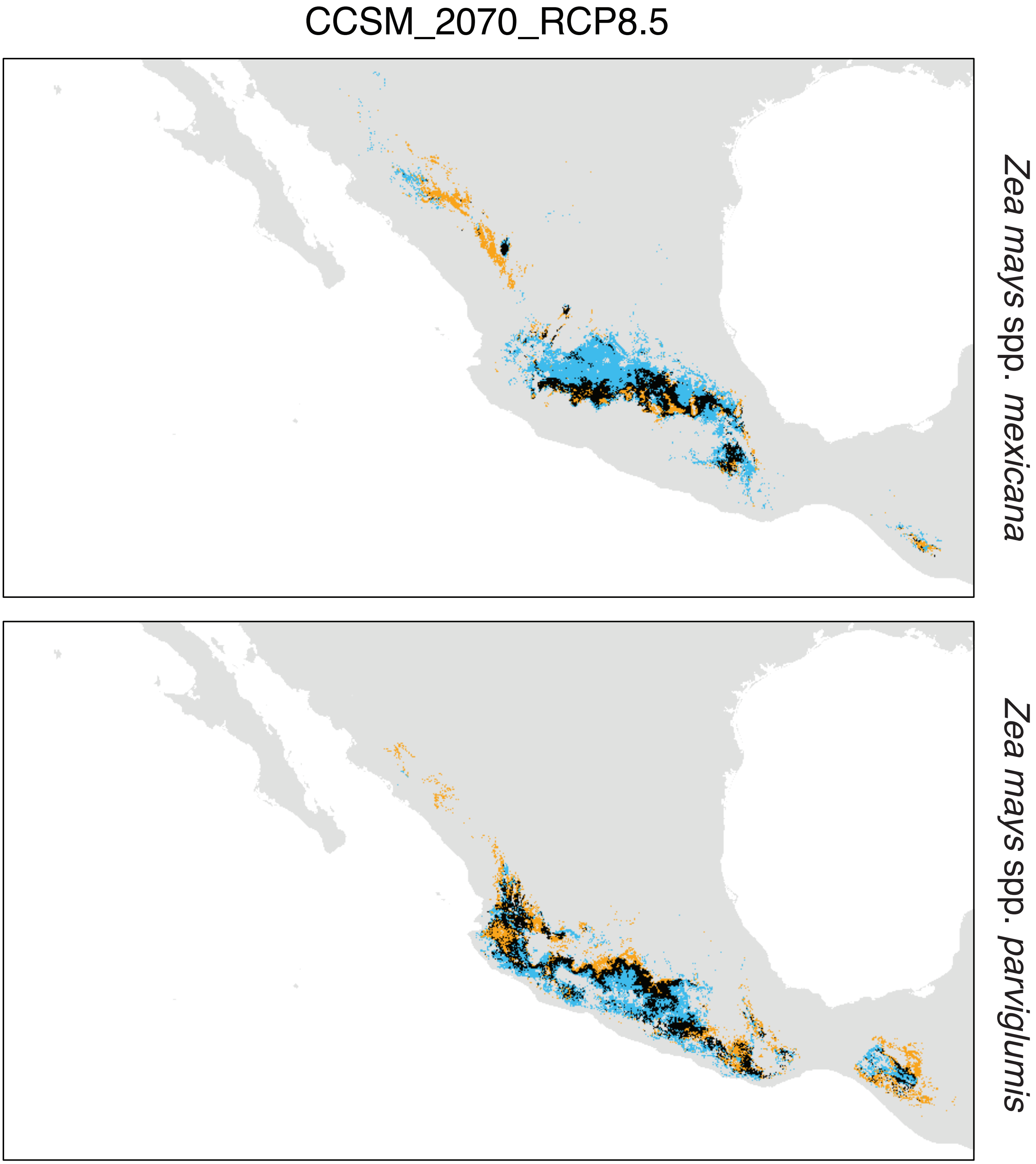

**Supplementary Figure 2.** Continued

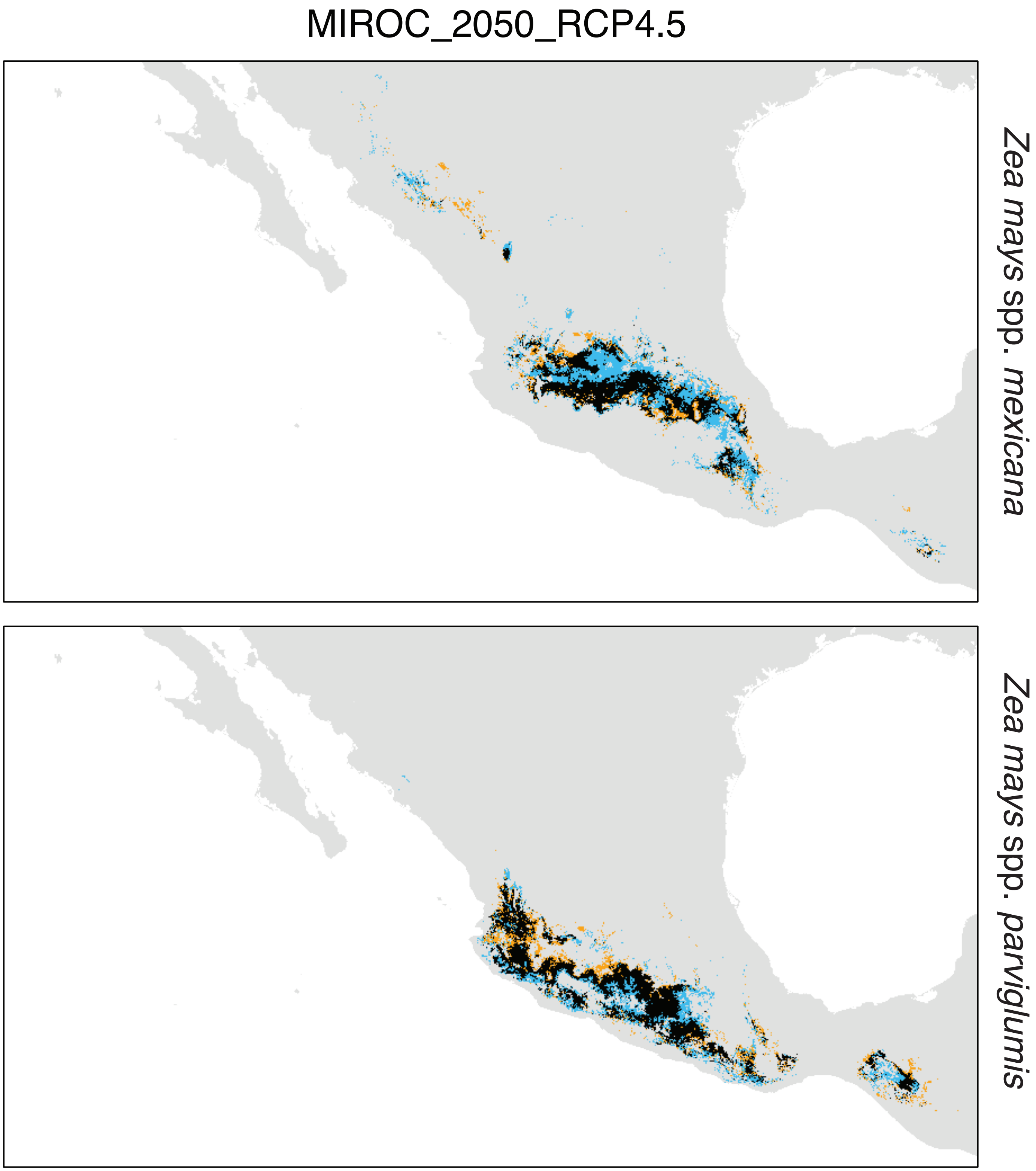

**Supplementary Figure 2.** Continued

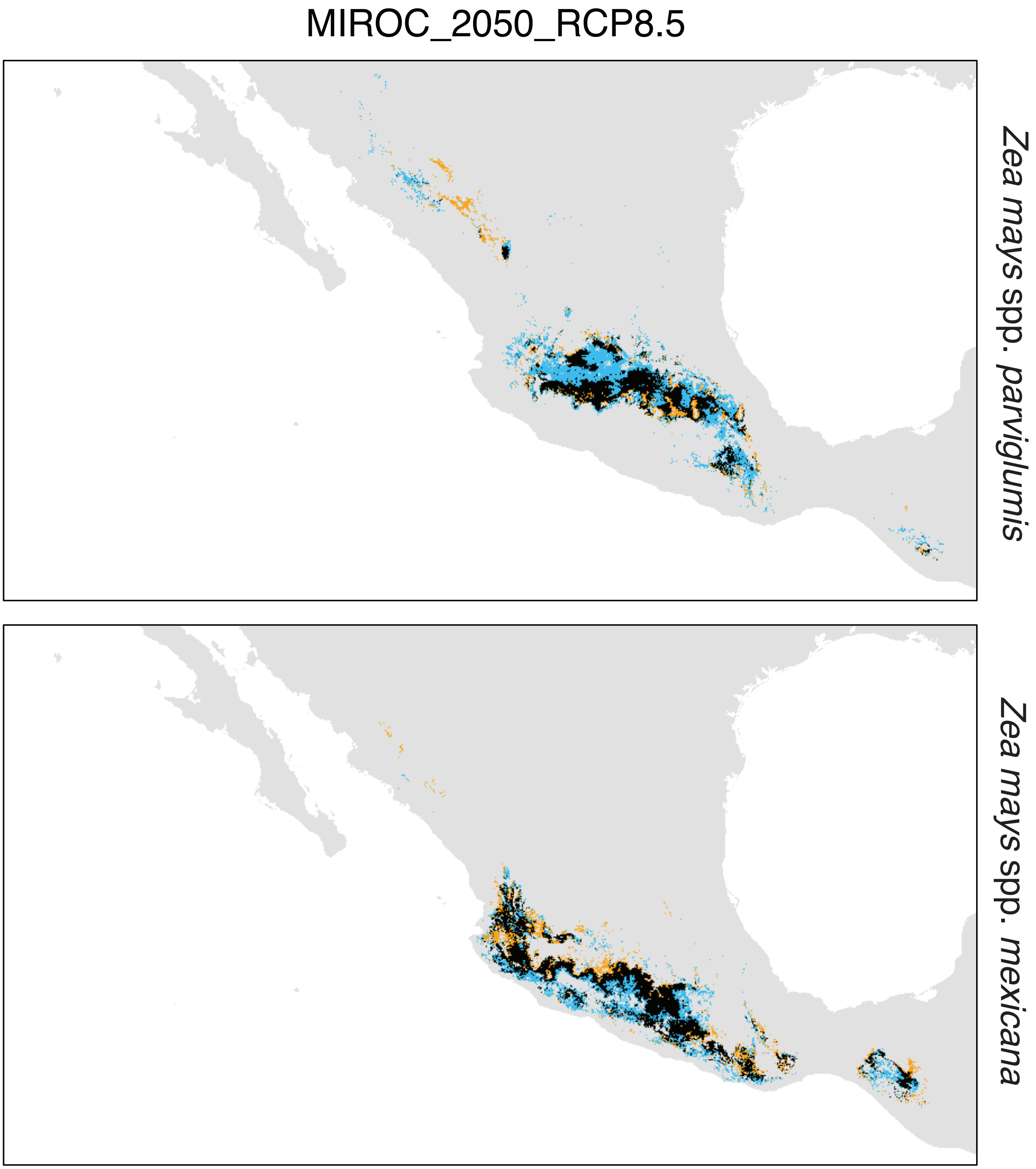

**Supplementary Figure 2.** Continued

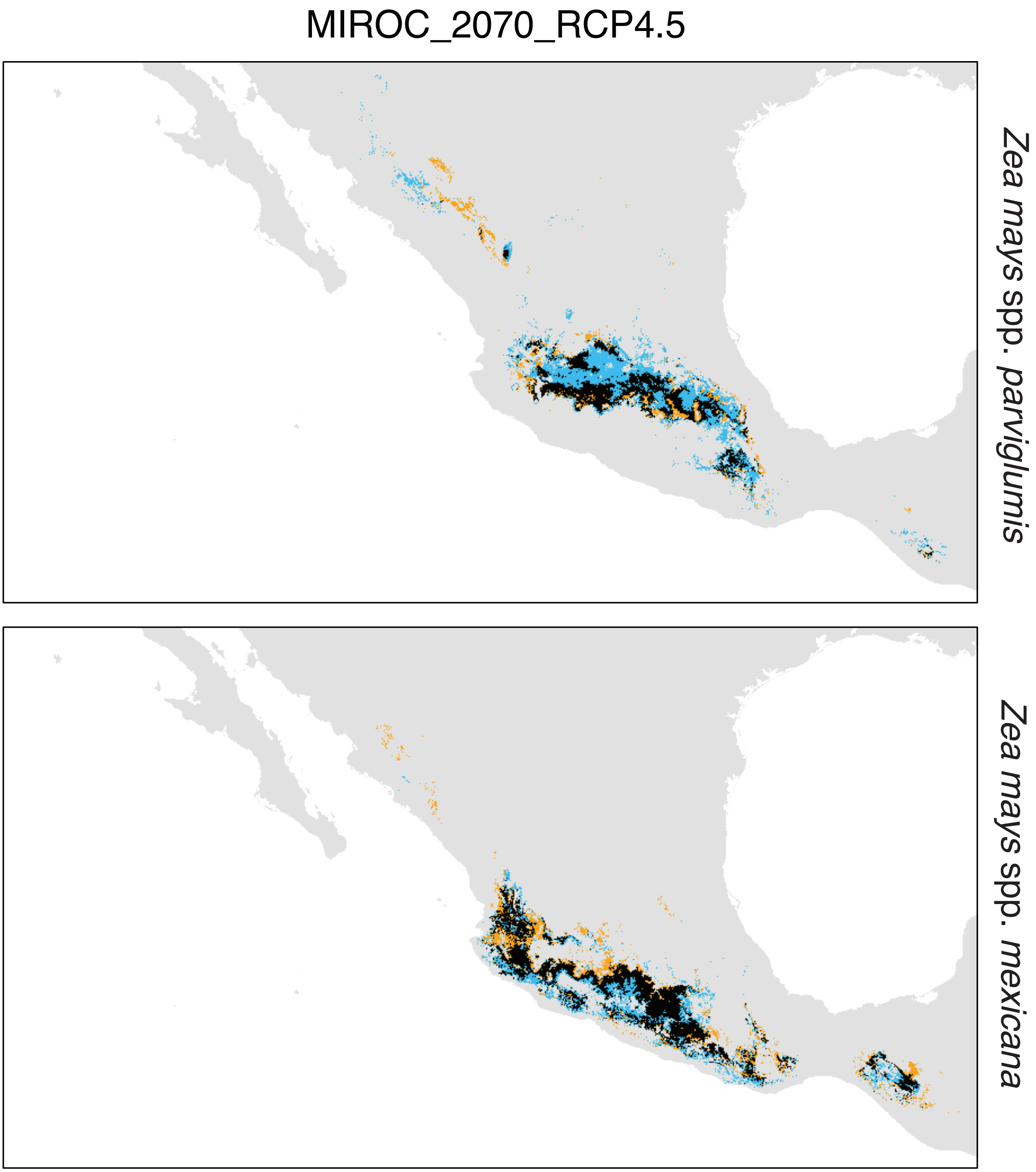

**Supplementary Figure 2.** Continued

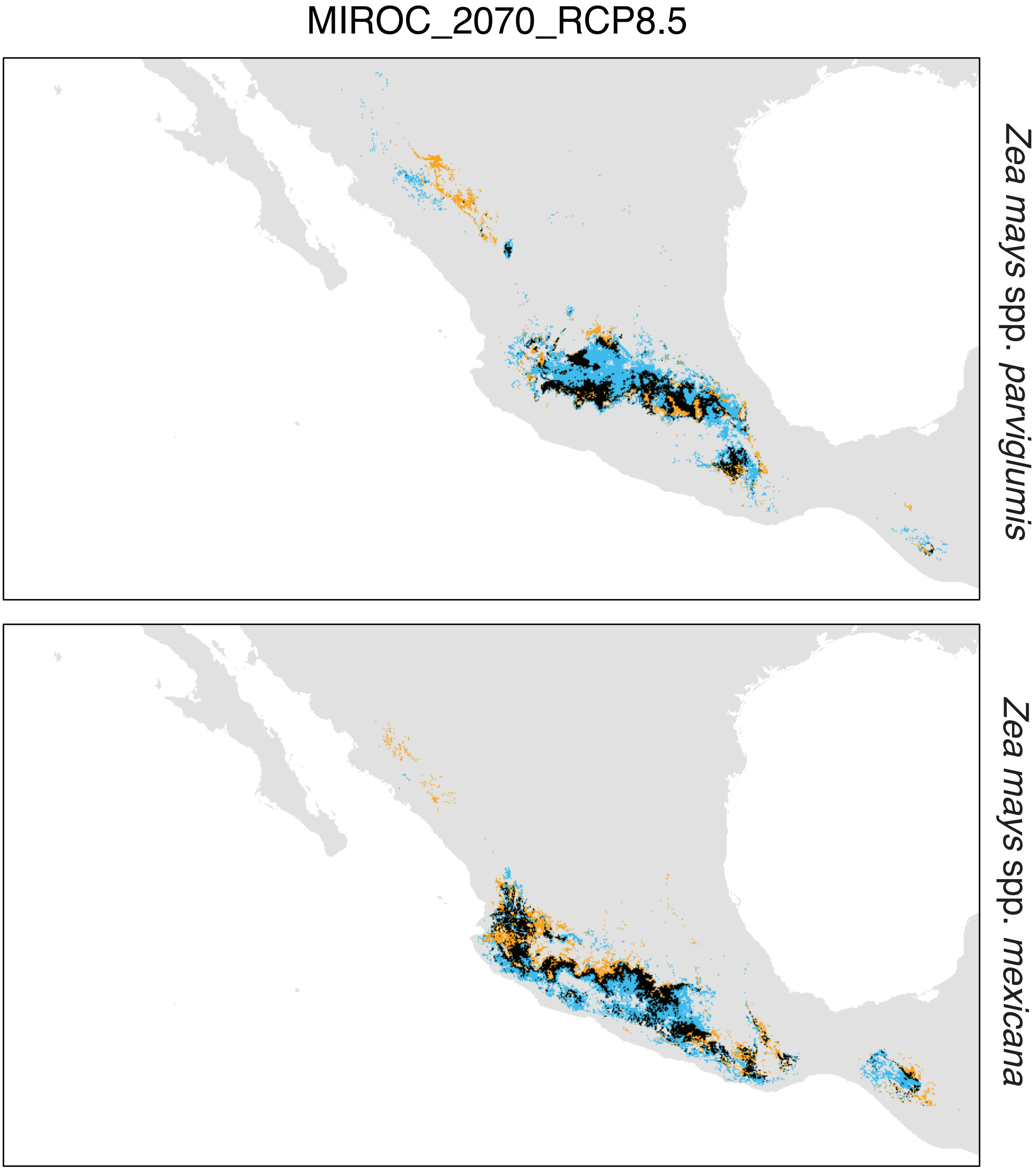

**Supplementary Figure 3. a-h,** Genomic offset estimated with Gradient Forest analyses under the eight models of climate change for sampled populations of two species of teosintes in Mexico (*Zea mays* spp. *parviglumis* and *Z. mays* spp. *mexicana*). **i**, Correlation between per-population genomic offset estimated with paSNPs (red) and canSNPs (orange) versus the rest of the SNPs sets across the eight models of climate change. paSNPs: putative adaptive; canSNPs: candidate; refSNPs: reference; n_refSNPS: reference controled for allele frequencies; USNPs: putative adaptive plus candidate;baySNPs: outlier detected with bayenv; scenvSNPs: outlier detected with bayescenv; bay∩scenvzSNPs: outlier detected with bayenv and bayescenv. Total number of SNPs in each category with significant contribution to the model are given in parenthesis.

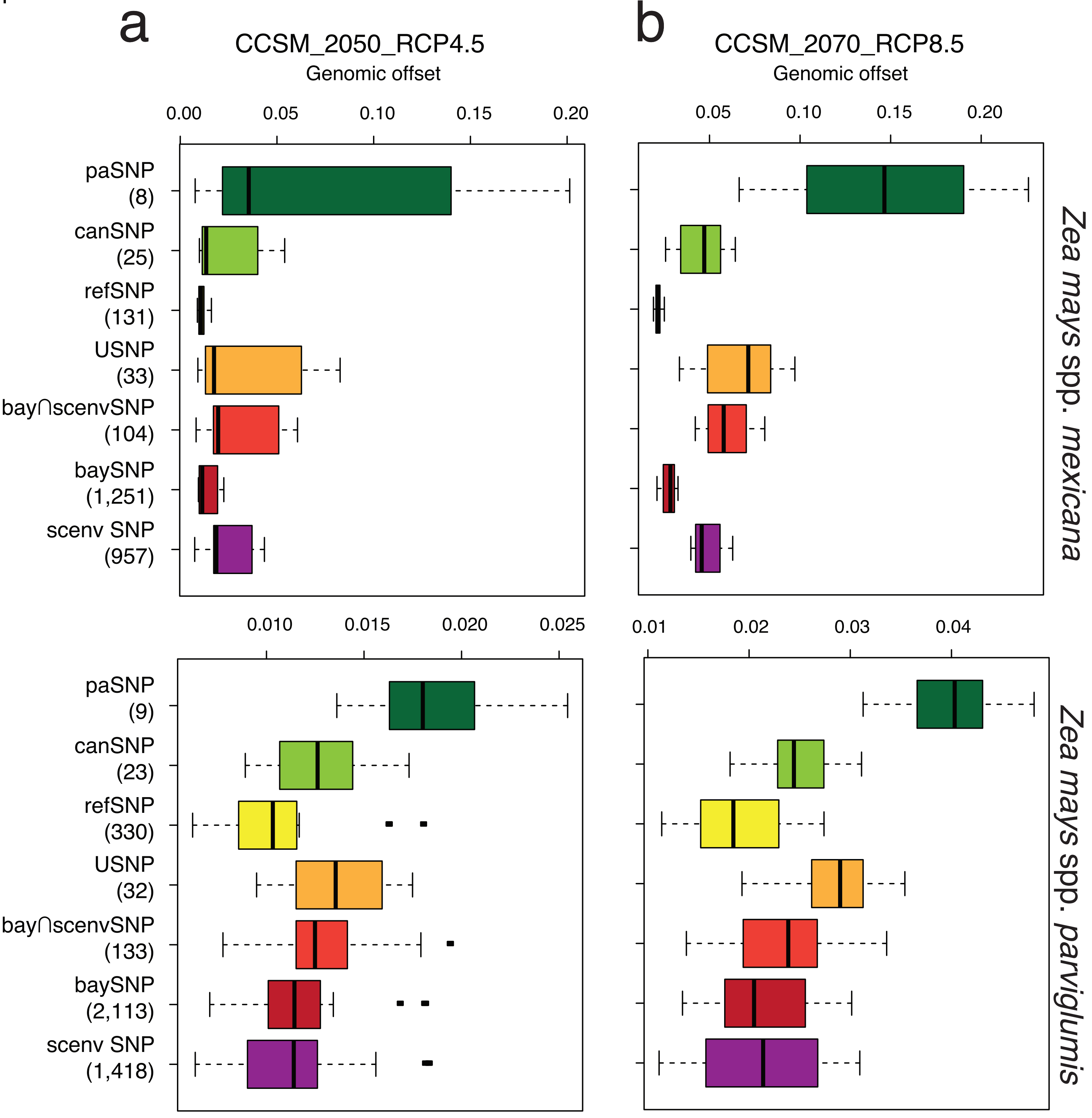

**Supplementary Figure 3.** continued

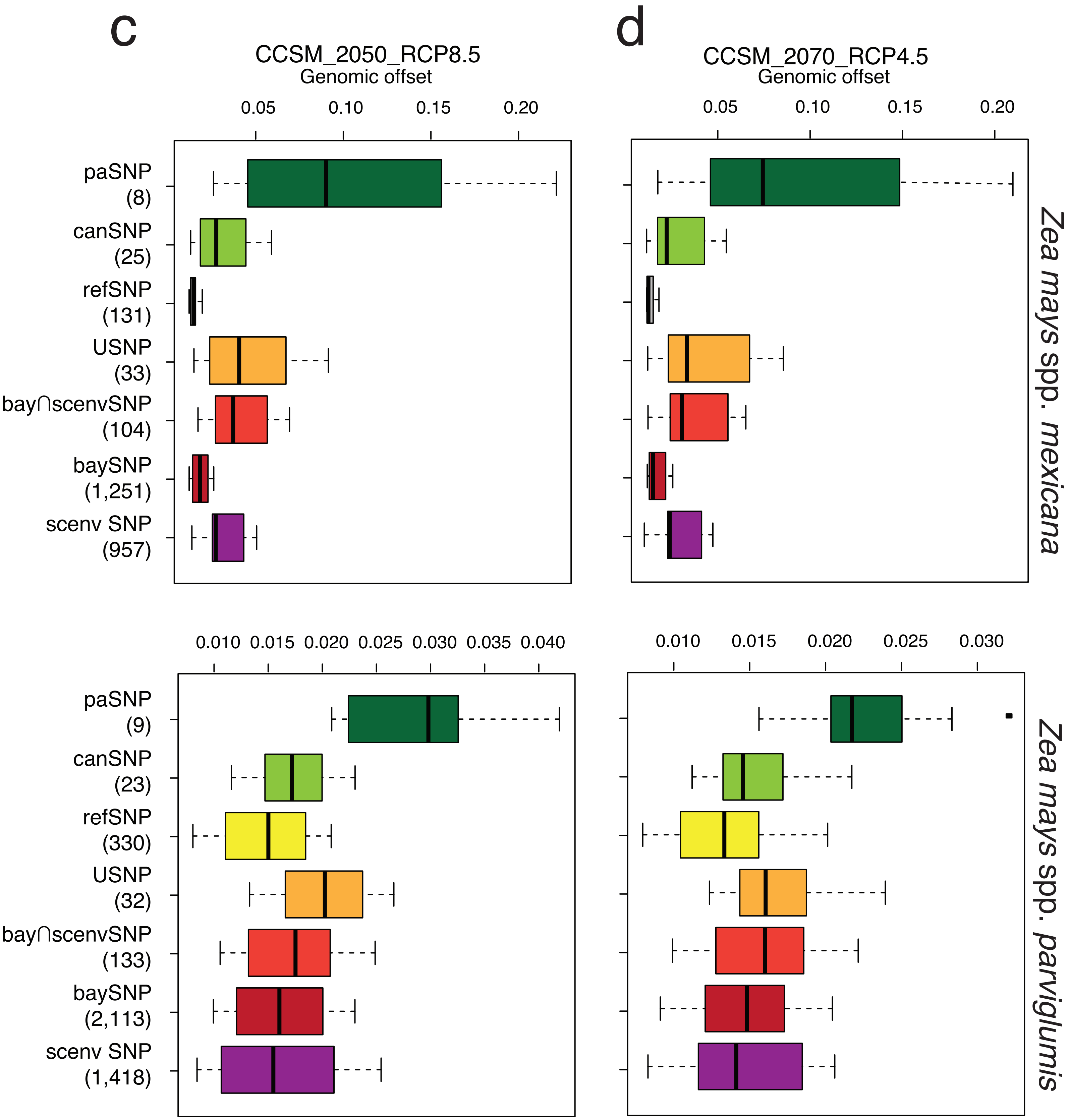

**Supplementary Figure 3.** continued

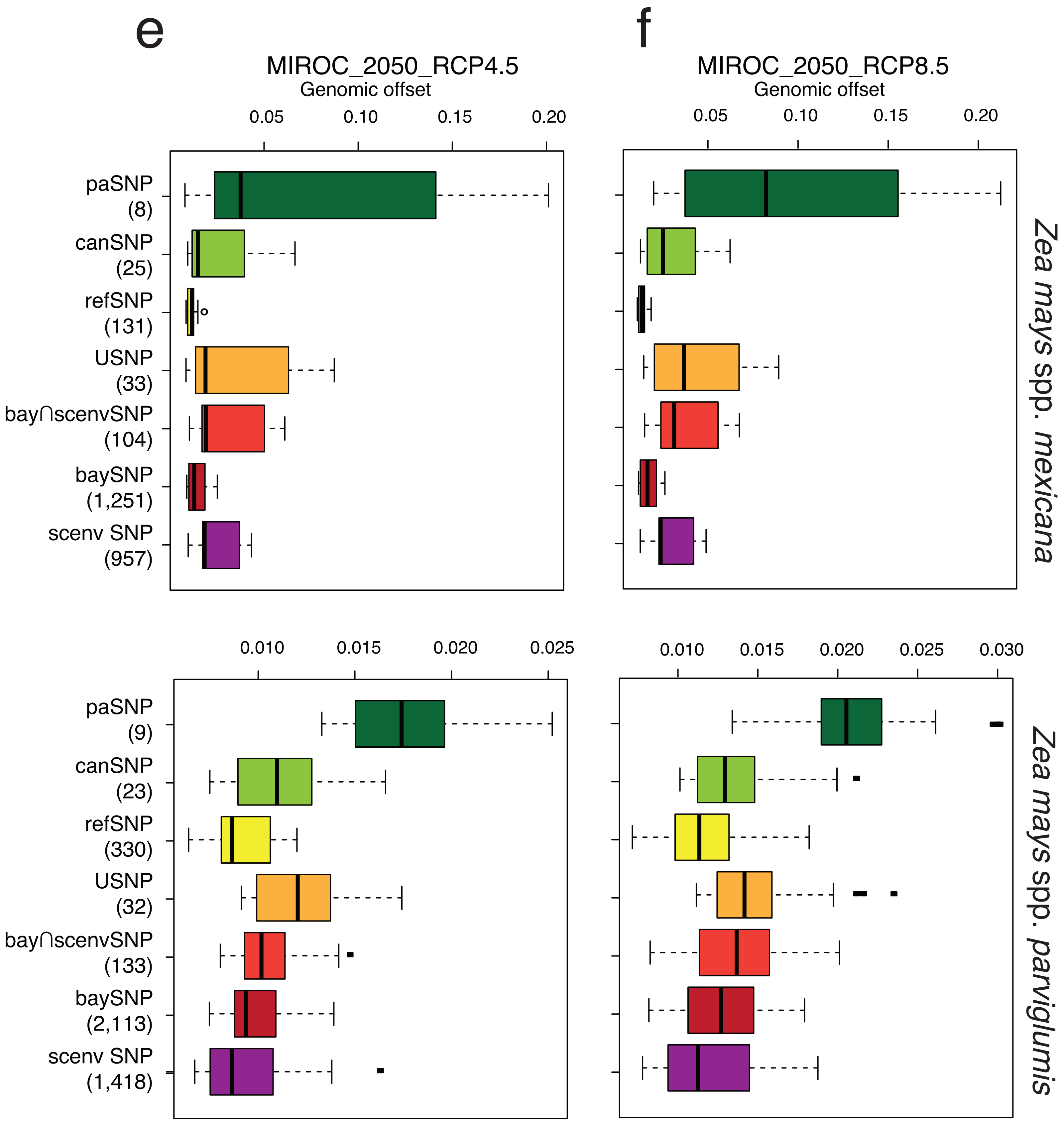

**Supplementary Figure 3.** continued

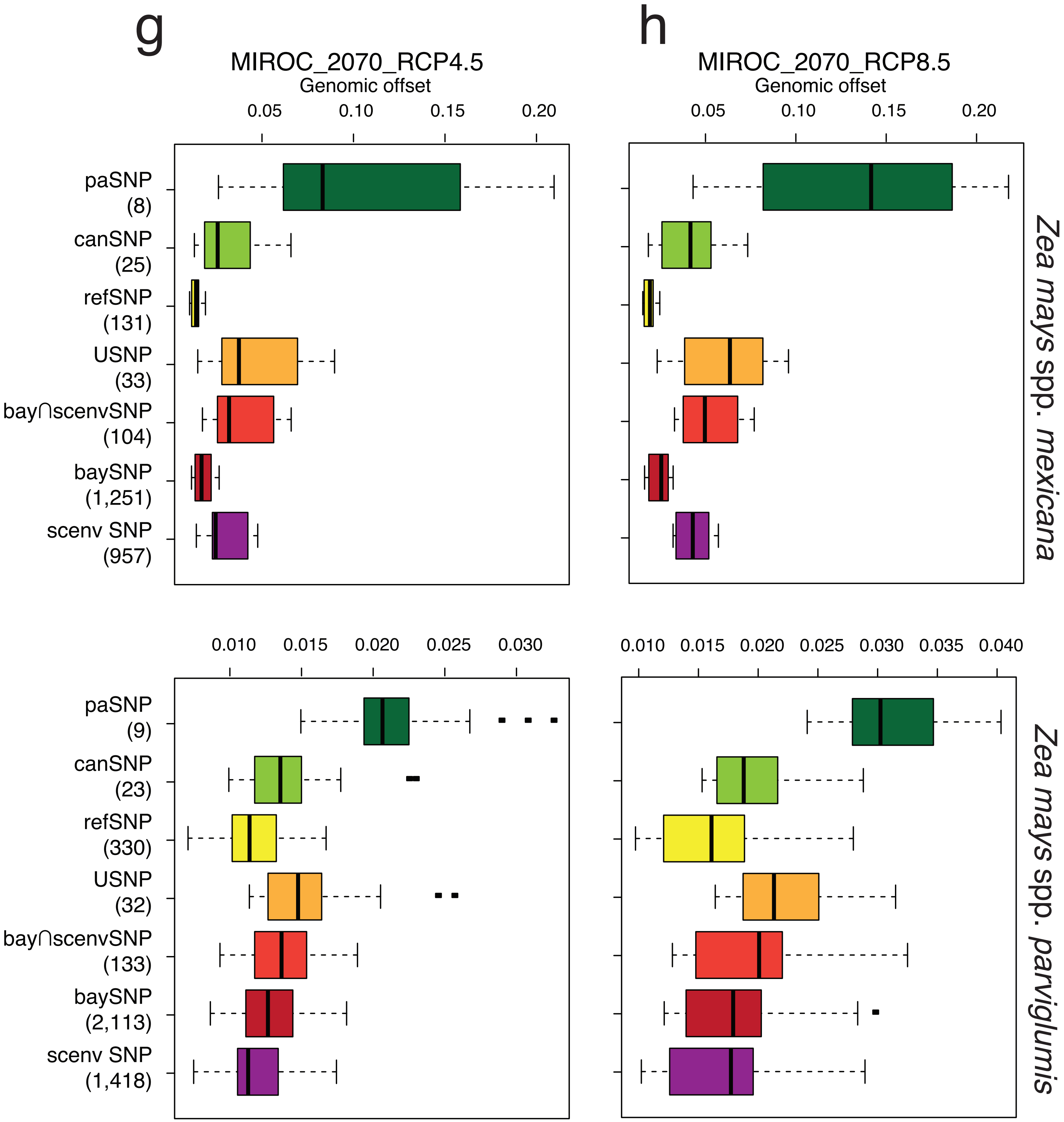

**Supplementary Figure 3.** continued

i

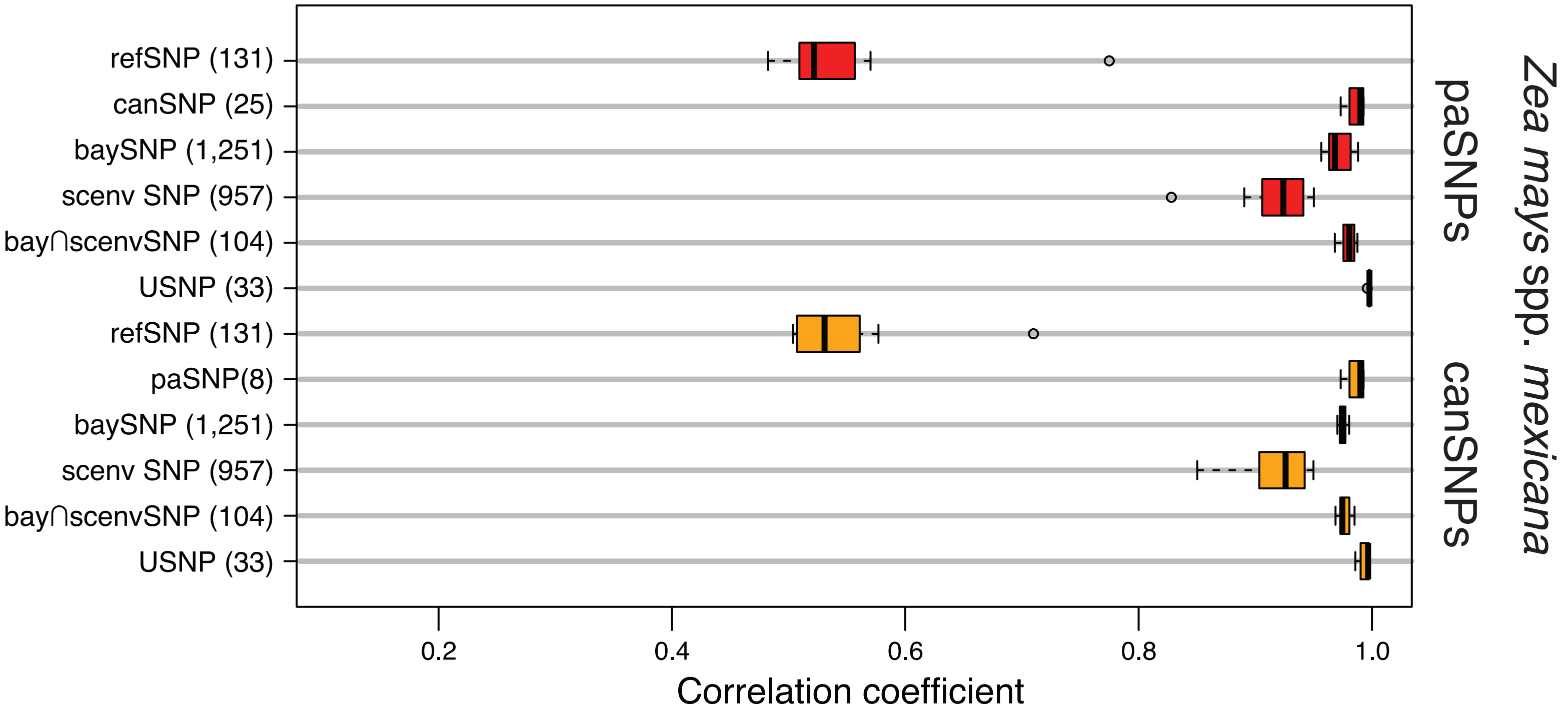


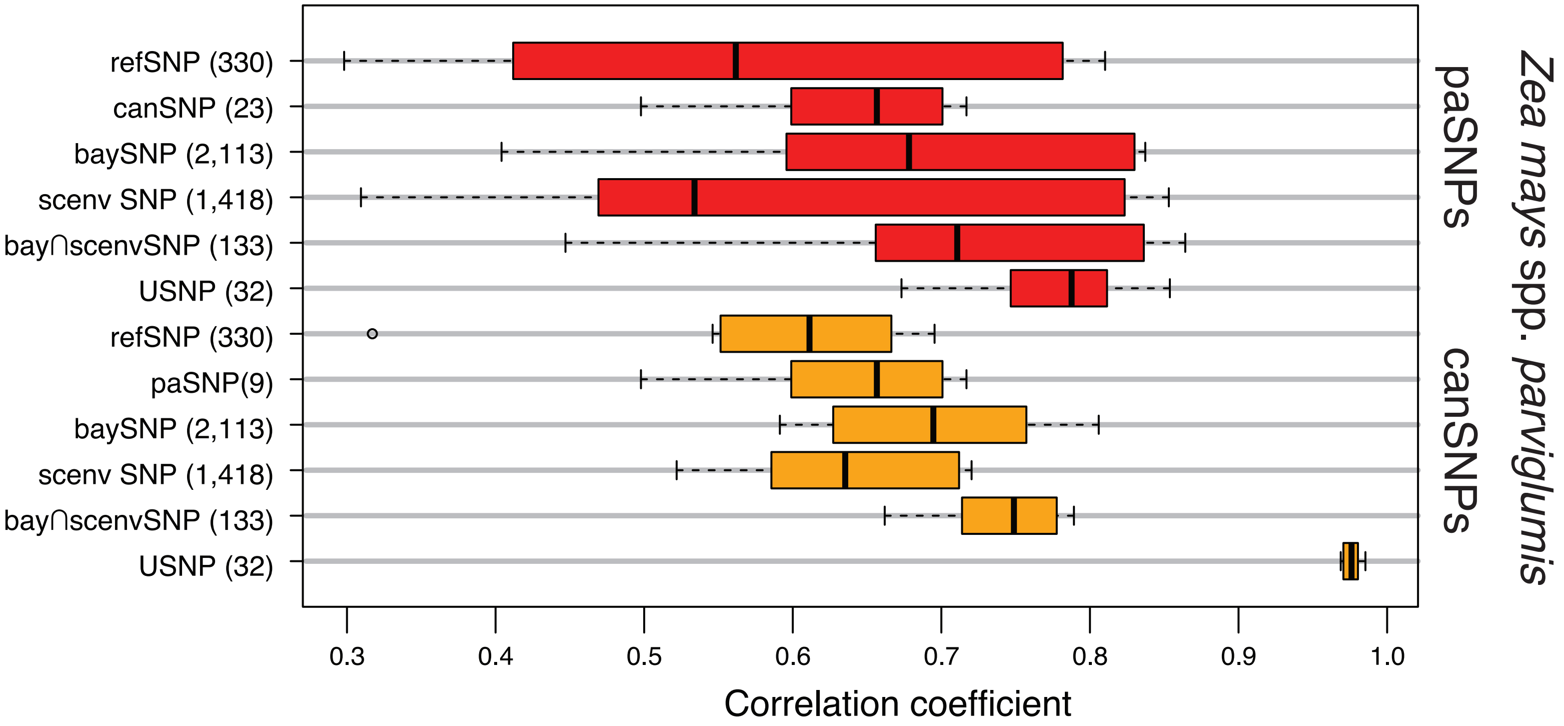

**Supplementary Figure 4.** Contribution of different SNPs to the allele turnover function ($R^2$) constructed with a genome-wide Gradient Forest analyses for sampled populations of two species of teosintes in Mexico: (*Zea mays* spp. *parviglumis* and *Z. mays* spp. *mexicana*). Allele turnover functions were constructed using the complete set of 33,454 SNPs identified in teosintes. Boxplots showing the distribution of $R^2$ among SNPs with varying levels of climate-frequency associations. Gray boxes represent the distribution of $R^2$ for SNPS subsets according to their overall contribution to the allele turnover function. For the definition of SNP categories we used the quartile distribution of $R^2$ (1Q, median, 3Q) estimated for all SNPs. Total number of SNPs in each category are given in parenthesis. The number of refSNPs is less than the 500 sampled because only the SNPs with significant contributions are plotted.

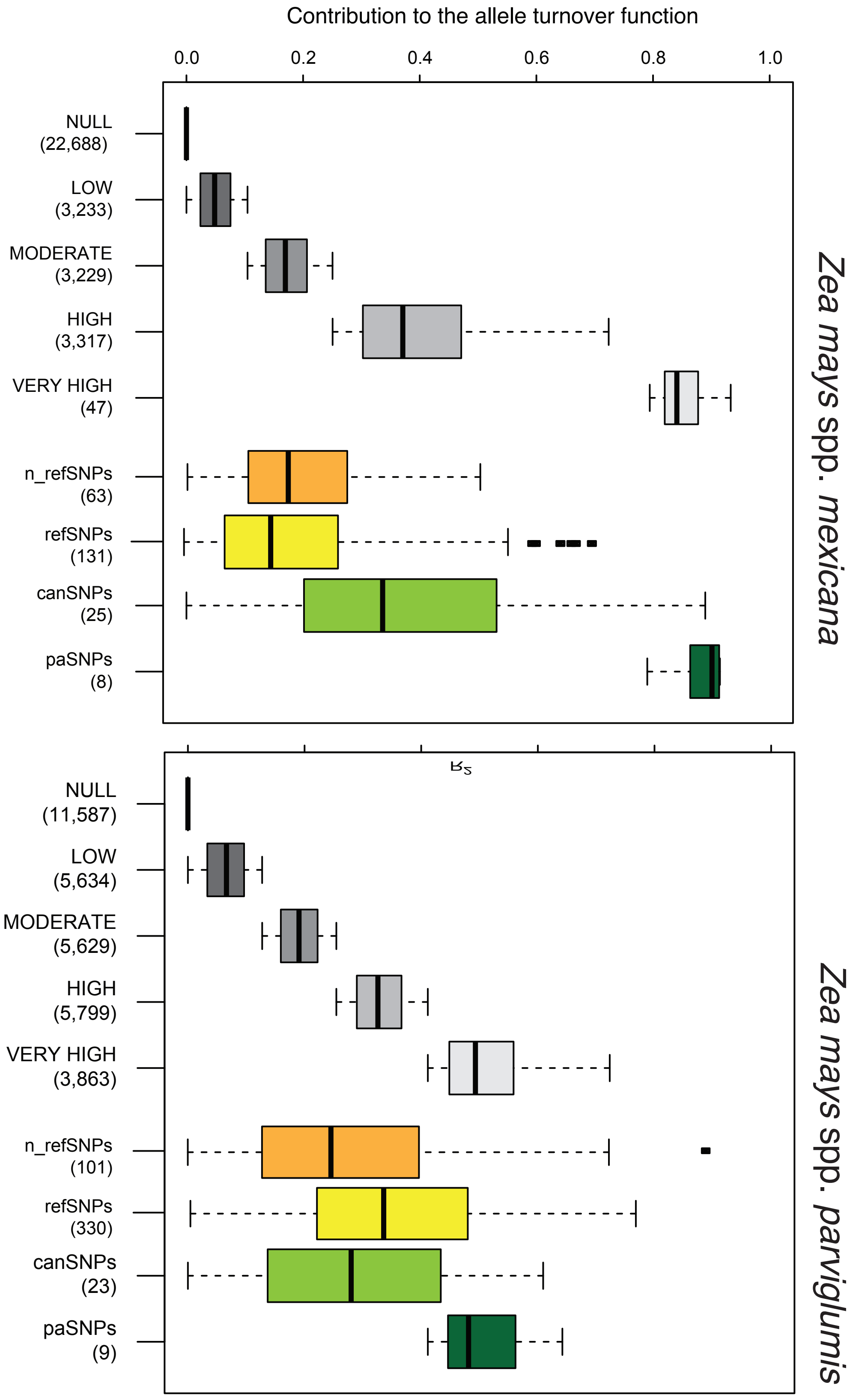

**Supplementary Figure 5.** Linear regressions between the estimated genomic offset under the two most extreme models of climate change (CCSM_2050_RCP4.5 and CCSM_2070_RCP8.5) versus the adaptive scores and the degree of climate change expected for sampled populations of two species of teosintes in Mexico: (*Zea mays* spp. *parviglumis* and *Z. mays* spp. *mexicana*). Open and black circles represent the genomic offset of populations under CCSM_2050_RCP4.5 and CCSM_2070_RCP8.5, respectively. Regression coefficients and statistical significance are given for each linear model.

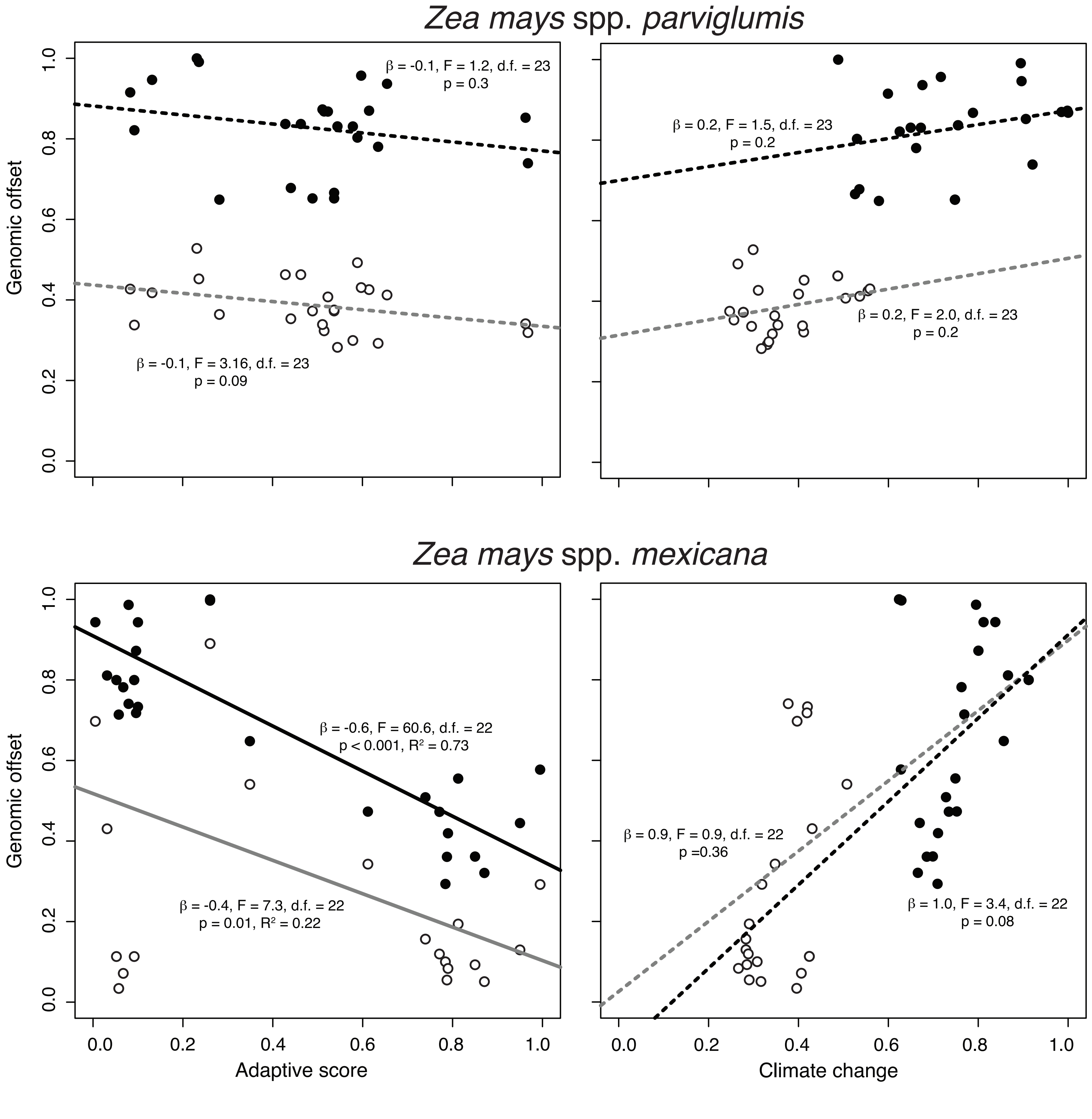

**Supplementary Figure 6.** Genomic offset estimated under the eight models of climate change for areas within the present-day and future allele distribution models for putative adaptive SNPs for two species of teosintes in Mexico: (*Zea mays* spp. *parviglumis* and *Z. mays* spp. *mexicana*). NP: areas outside the present-day and future distribution of paSNPs; P: areas only within the present-day distribution of paSNPs; F: areas only within the future distribution of paSNPs; P+F: areas within the overlapped present-day and future distribution of paSNPs. Missing values for F and P+F signify that predicted areas overlap with the present-day models.

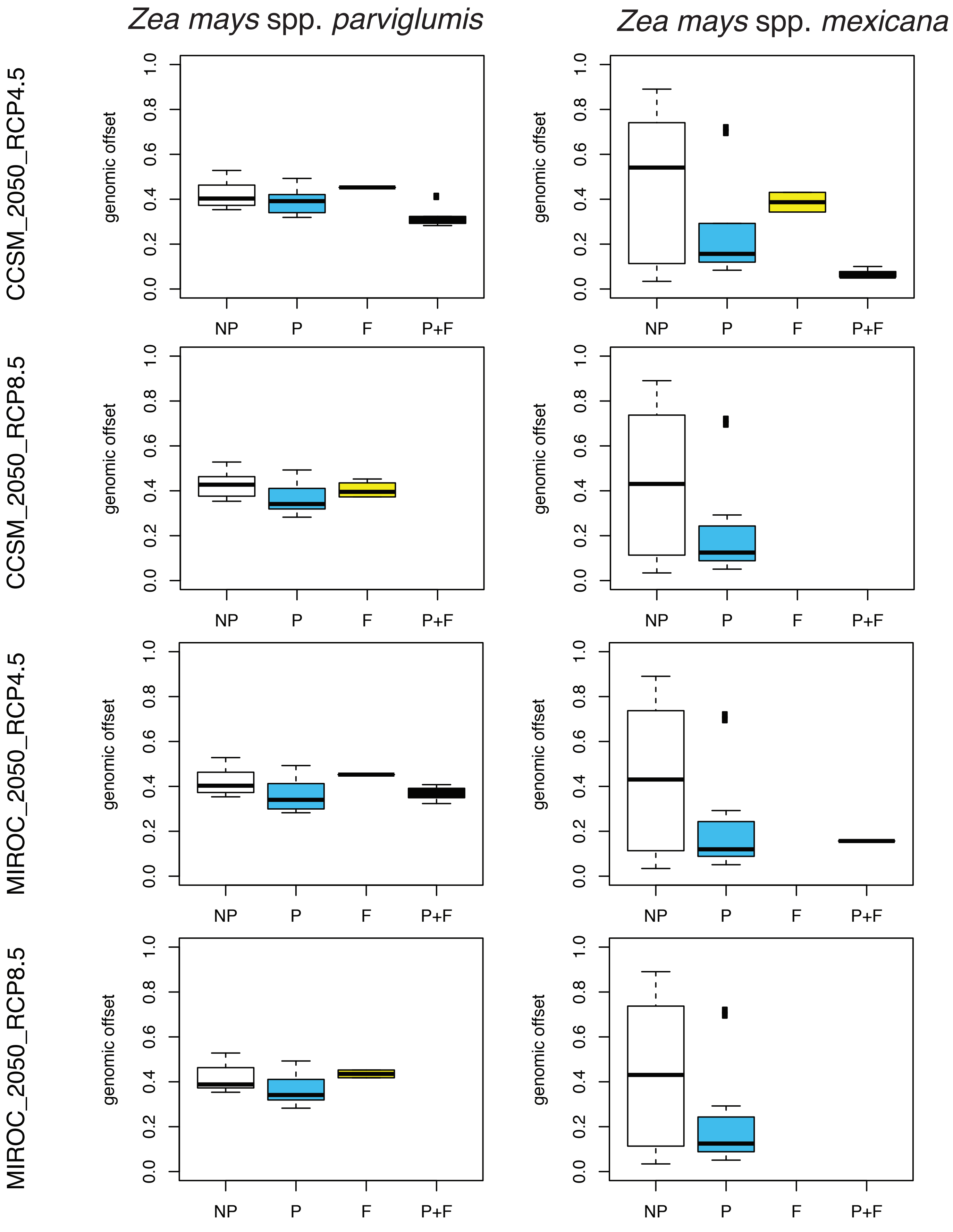

**Supplementary Figure 6.** Continued

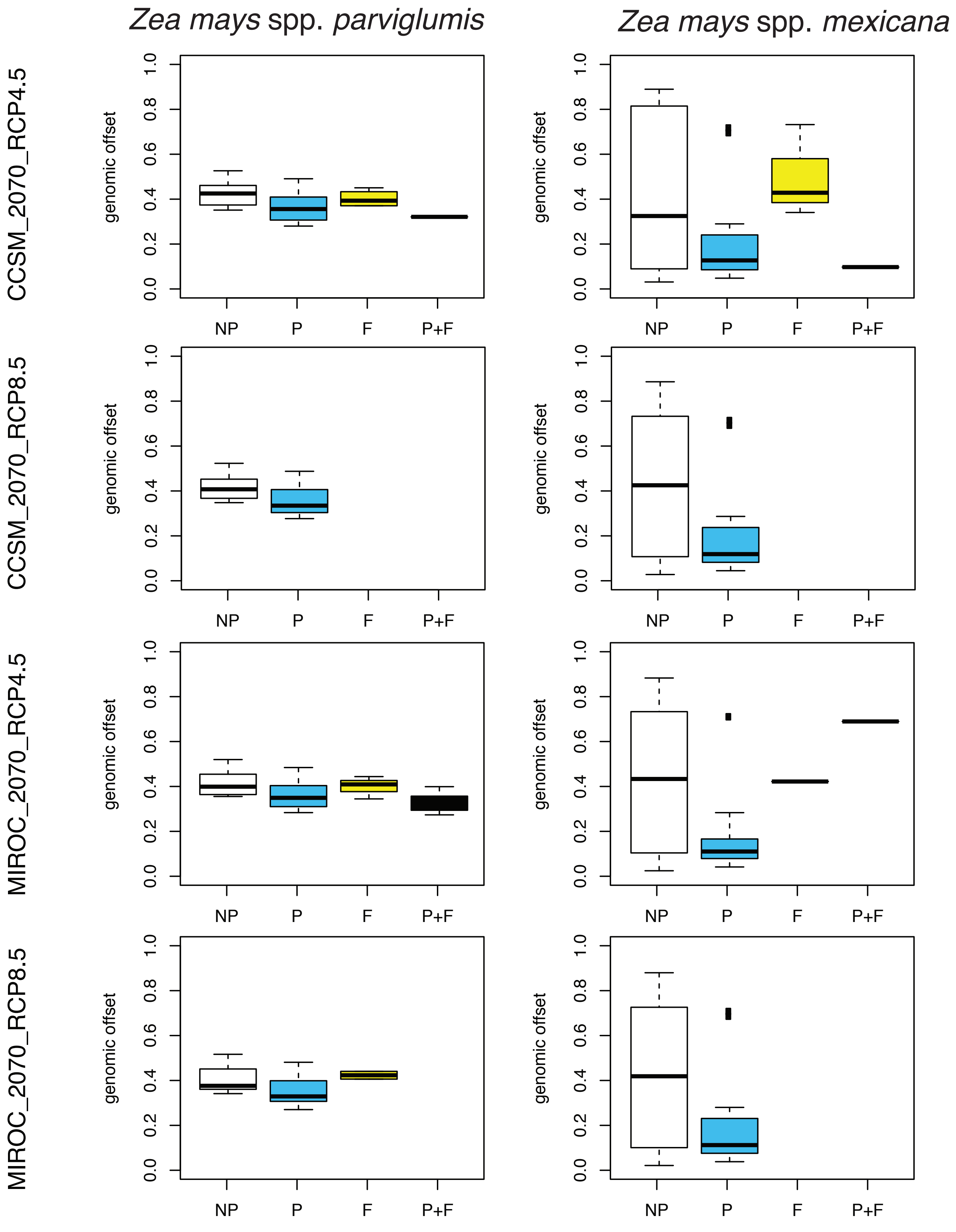

**Supplementary Figure 7.** Linear regressions between the estimated genomic offset under the eight models of climate change versus populations' genetic diversity (Hs) of sampled populations of two species of teosintes in Mexico: (*Zea mays* spp. *parviglumis* and *Z. mays* spp. *mexicana*). Regression coefficients and statistical significance are given for each linear model.

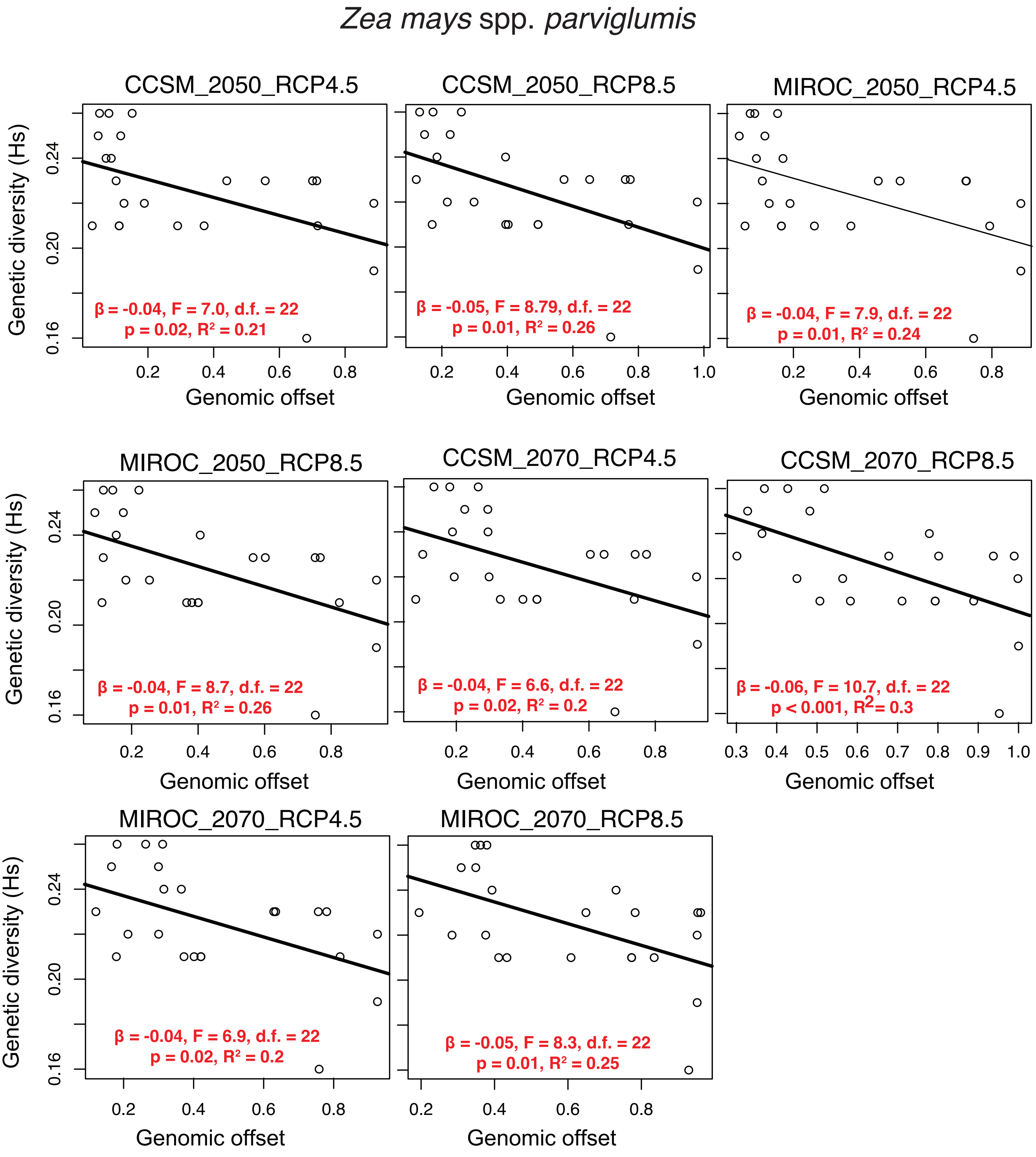

**Supplementary Figure 7.** continued

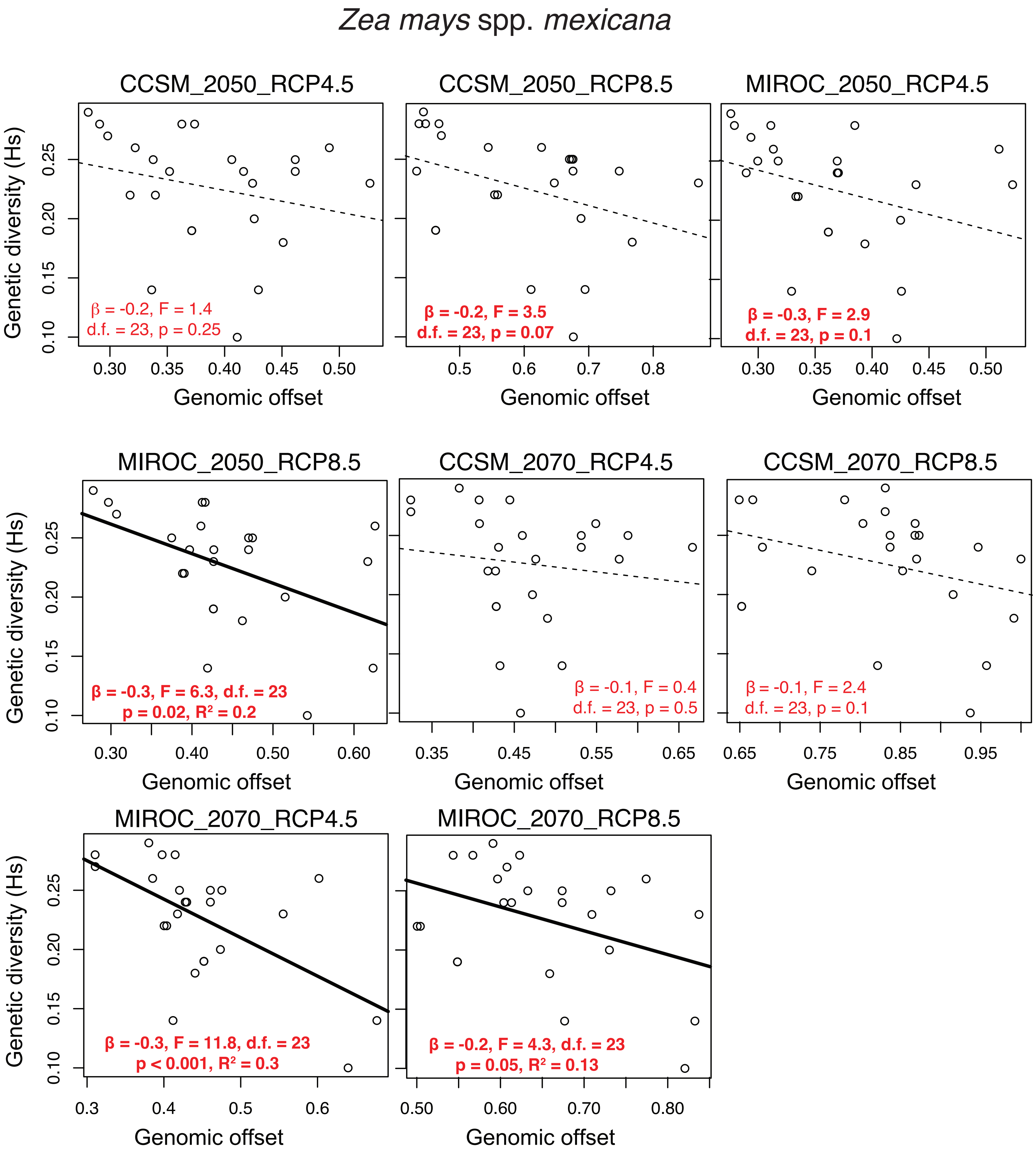

**Supplementary Figure 8.** Adaptive score and genomic offset as a function of migration probabilities estimated using circuit theory and allele distribution models under the two most extreme models of climate change (CCSM_2050_RCP4.5 and CCSM_2070_RCP8.5) for sampled populations of two species of teosintes in Mexico: (*Zea mays* spp. *parviglumis* and *Z. mays* spp. *mexicana*).Genomic offset was estimated using the paSNPs. Except otherwise stated, F-statistics are statistically significant with p-val < 0.01 (d.f. = 21).

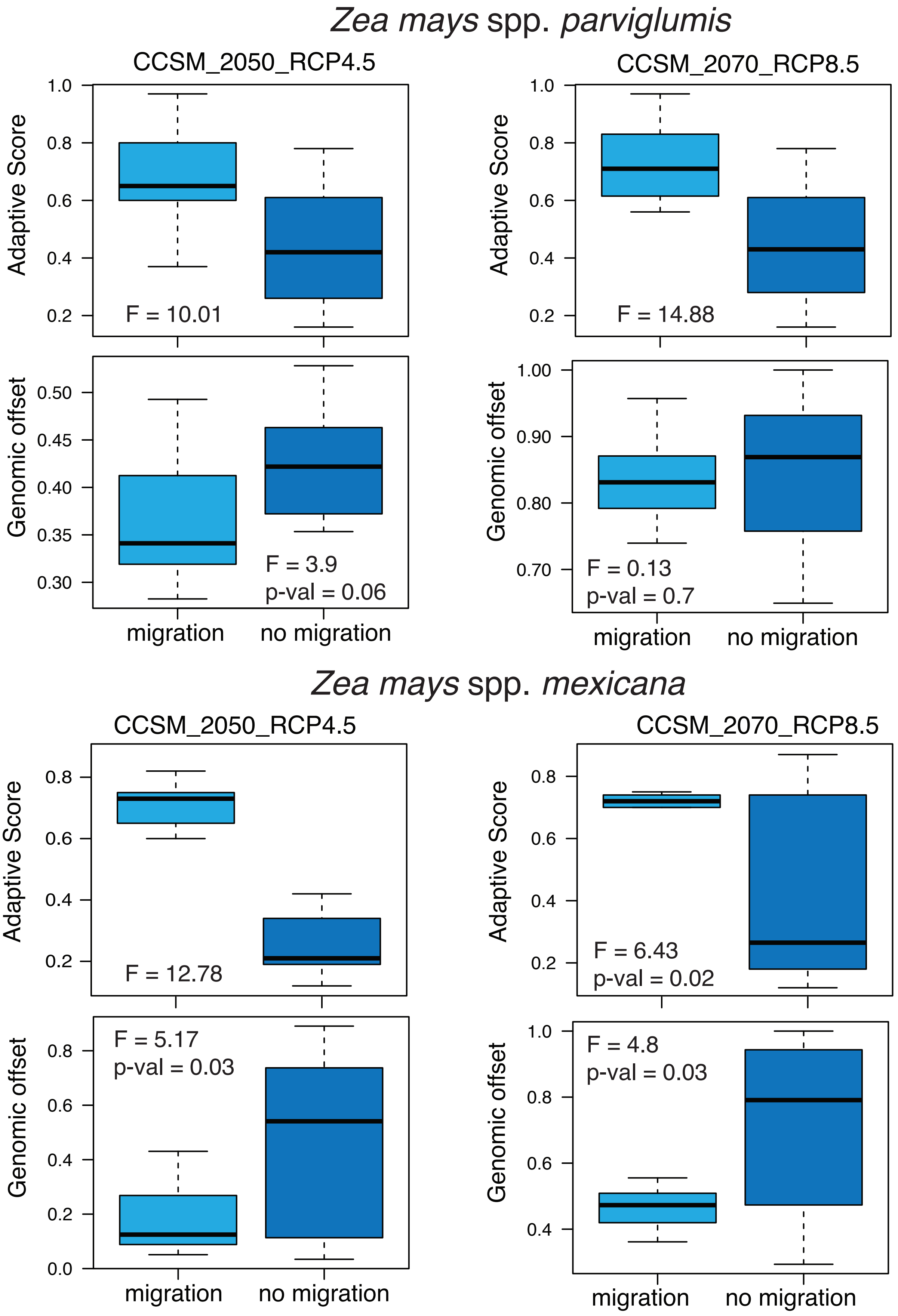

**Supplementary Figure 9.** Linear regressions between the estimated adaptive score versus mean annual temperature, geographic range size, and mean genomic offset for 46 maize landraces in Mexico. a, Association between adaptive scores using paSNPs and mean temperature across known occurrence of landraces (same as in Figure 2d). b, Association between adaptive scores using the paSNPs and geographic range size for landraces (same as in Figure 2c). c-d, Association between adaptive scores using paSNPs and mean genomic offset estimated from known occurrence of landraces under two most extreme models: CCSM_2050_RCP4.5 (same as in Figure 2e) and CCSM_2070_RCP8.5. Frequency histograms show the distribution of regression coefficients and adjusted $R^2$ estimated for 1,000 sub-samples of refSNPs and the red vertical lines represent the estimates using paSNPs.

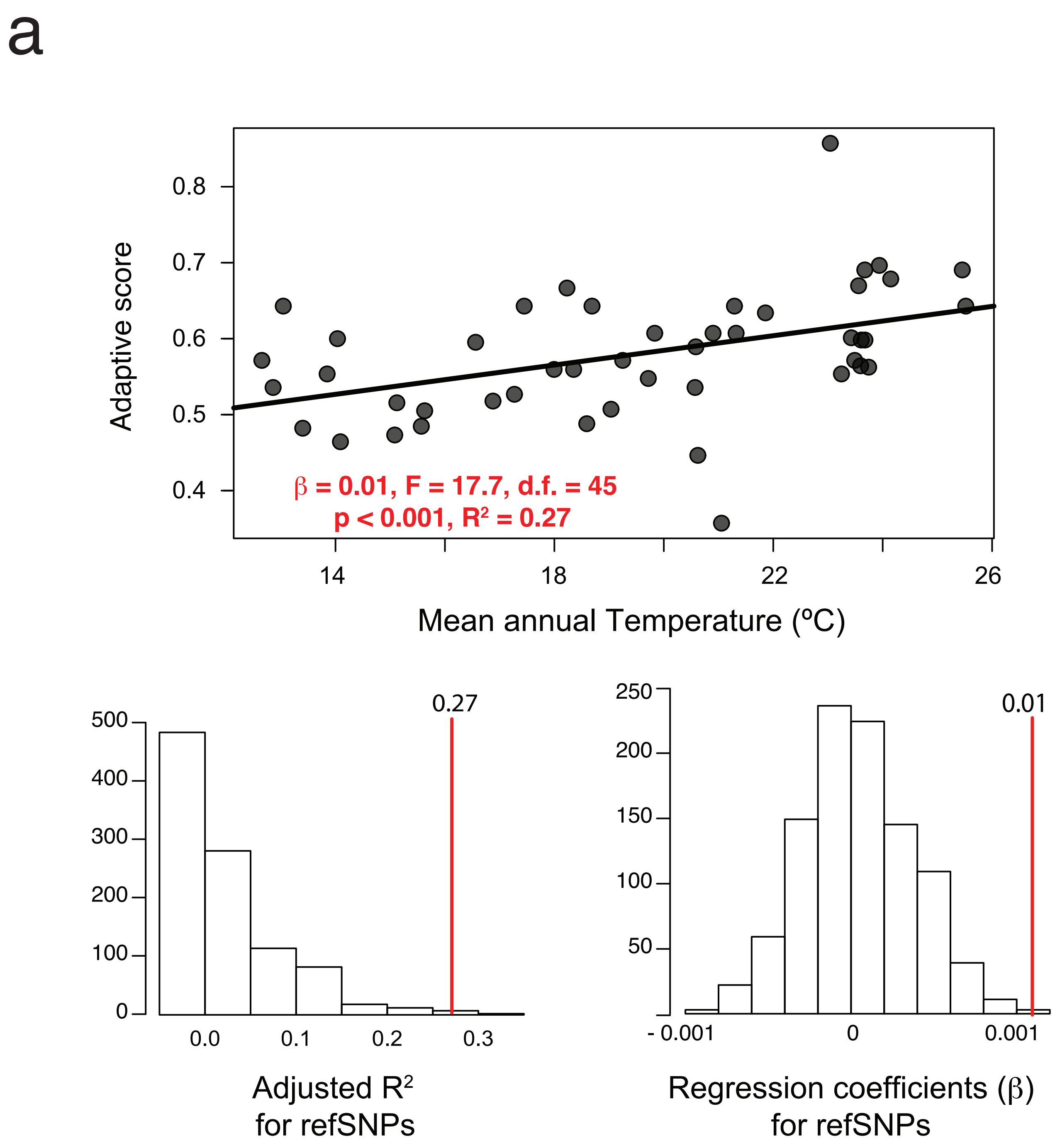

**Supplementary Figure 9.** Continued

b

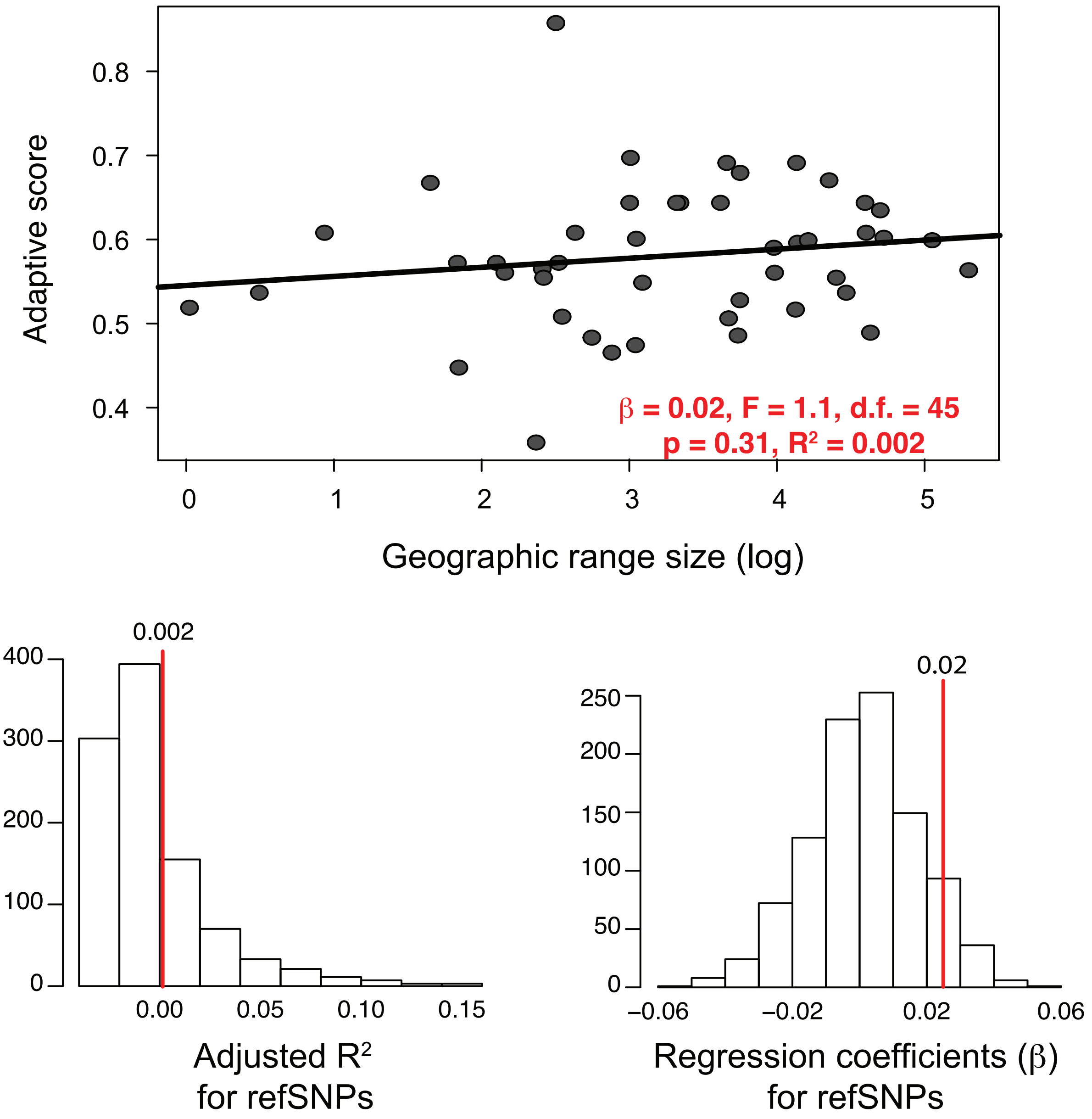

**Supplementary Figure 9.** Continued

C

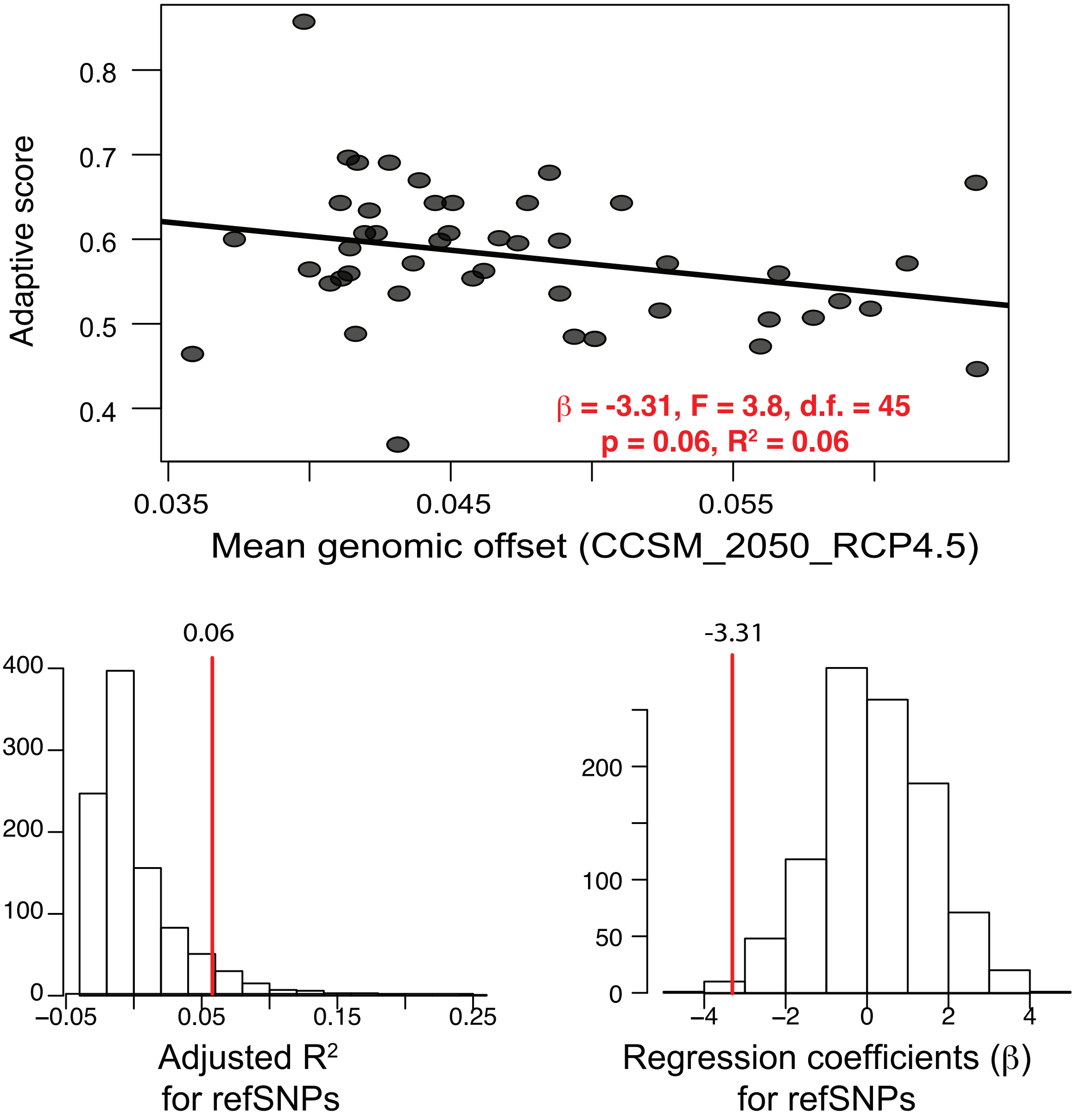

**Supplementary Figure 9.** Continued

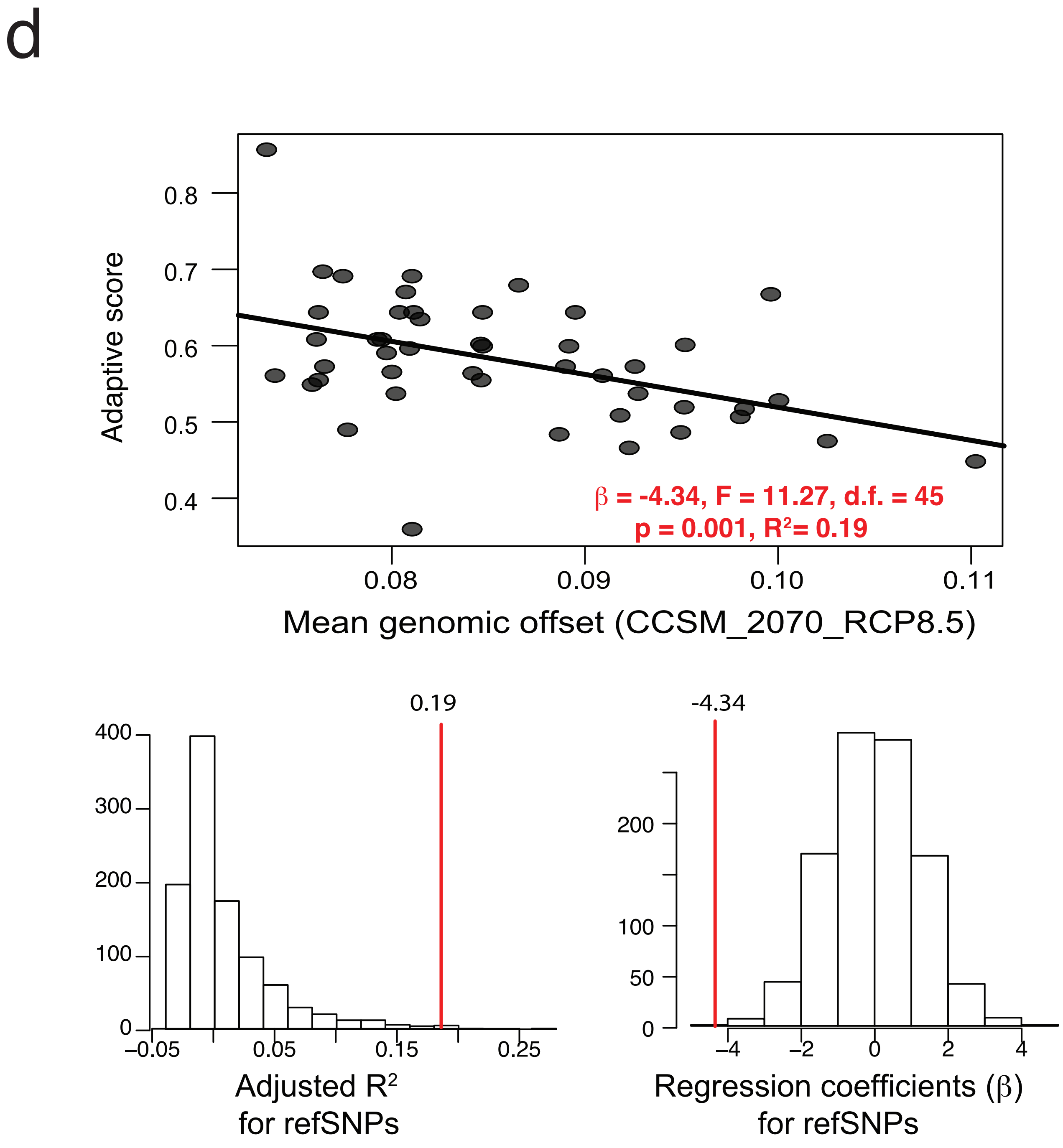

**Supplementary Figure 10.** Correlations between pairwise genetic differentiation ($F_{ST}$) and estimated allele turnover as predicted by Gradient Forest among sampled populations of two species of teosintes in Mexico: (*Zea mays* spp. *parviglumis* and *Z. mays* spp. *mexicana*). Allele turnover and $F_{ST}$ were estimated separately for each set of SNPs. a-b, correlations for reference SNPs. c-d, correaltions for candidate SNPs. e-f, correlations for putative adaptive SNPs. Correlation statistics (r) are given for partial Mantel tests performed controlling for environmental distances among populations. All correlations were sistically significant with p-value < 0.001.

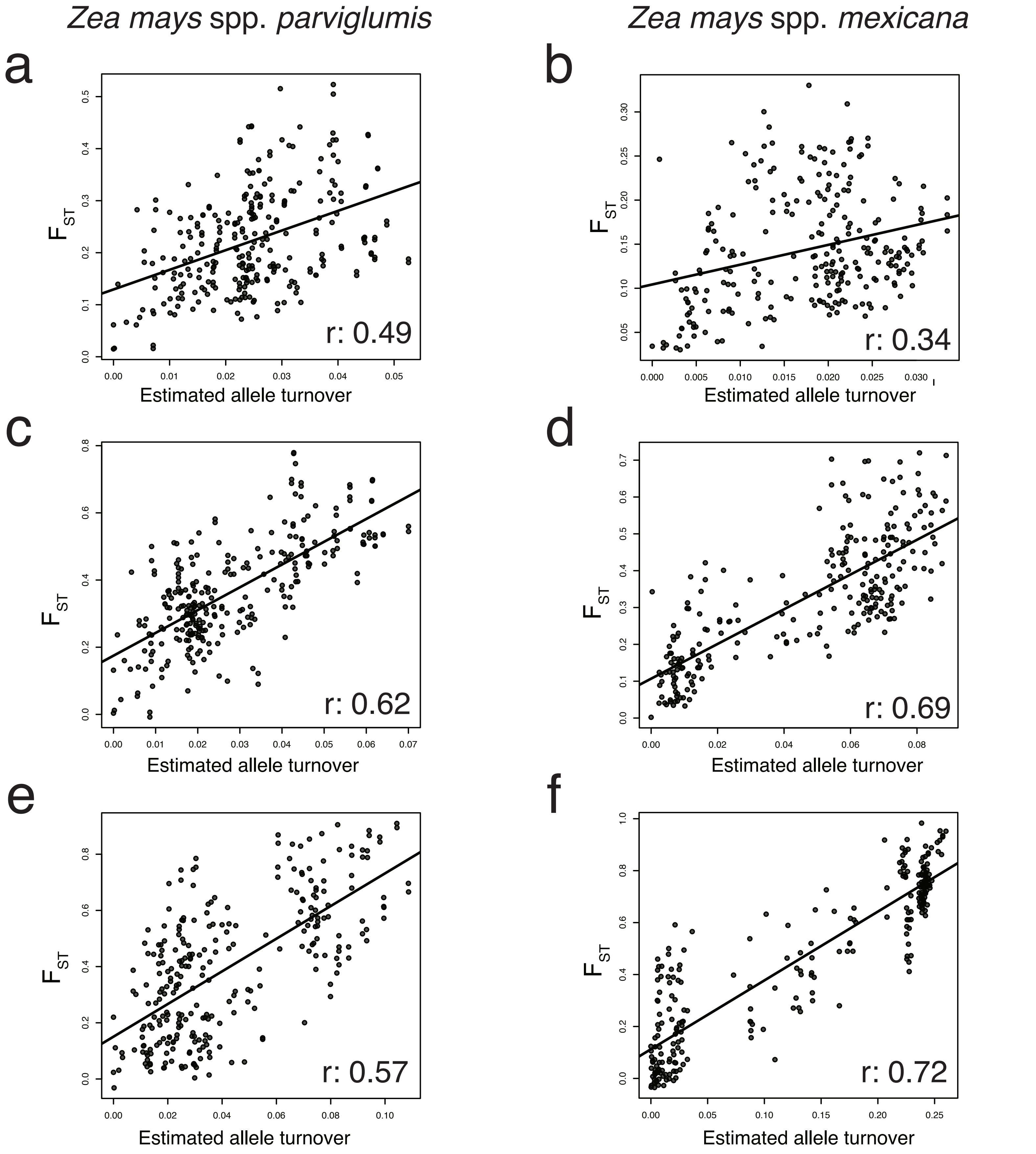

**Supplementary Figure 11.** Linear regressions between climatic suitability values for candidate and putative adaptive SNPs as predicted by ecological niche modeling and allele frequencies for sampled population of two species of teosintes in Mexico: (*Zea mays* spp. *parviglumis* and *Z. mays* spp. *mexicana*). Ecological niche models were constructed for each SNPs separately (models available in ASCII format at https://github.com/spiritu-santi/teosintes) and mean suitability was estimated for candidate SNPs and putative adaptive SNPs separately. All regressions were statistically non-significant with p-value > 0.05.

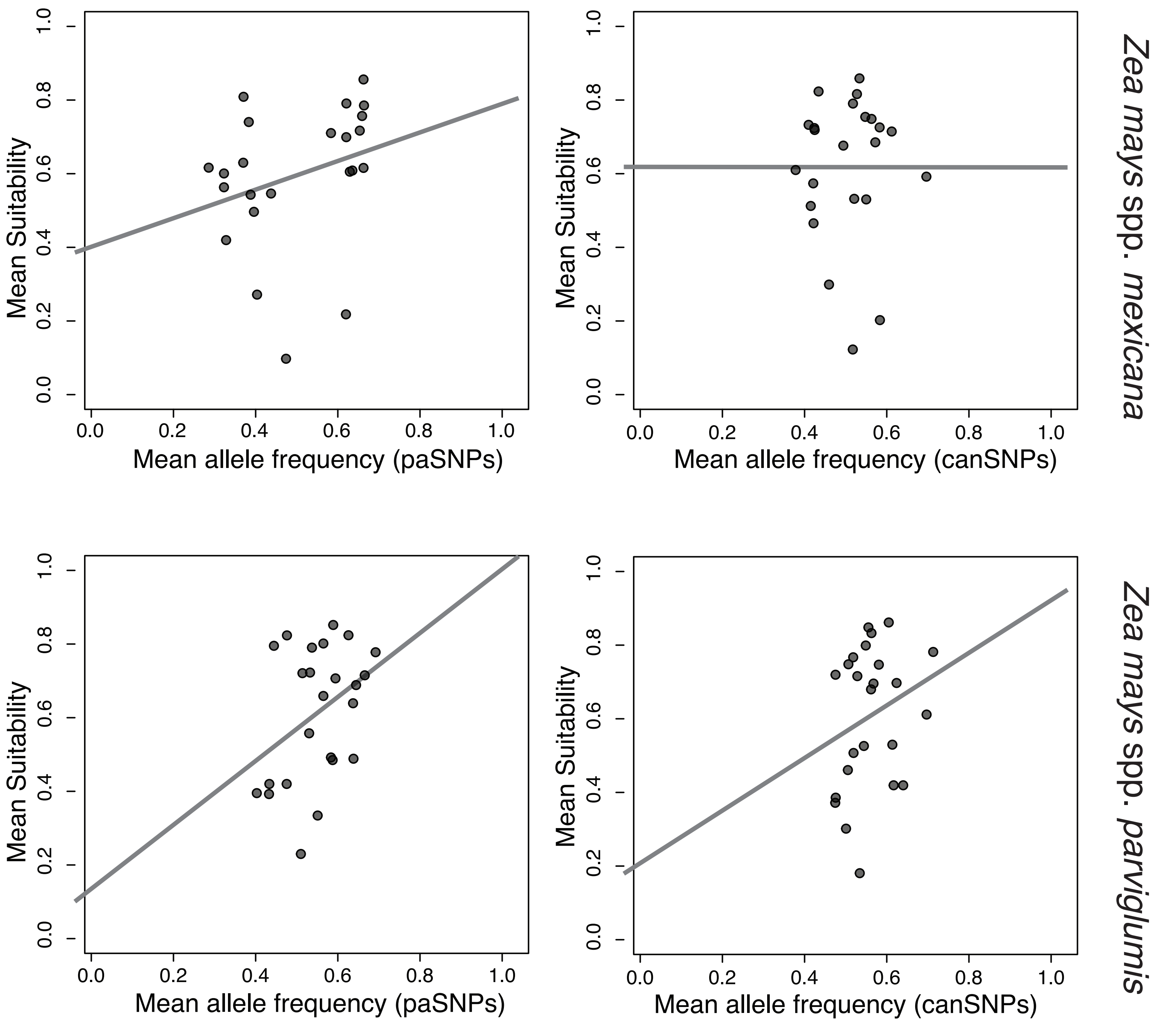

**Supplementary Figure 12.** Minimum geogaphic distances to population with the highest adaptive score as a function of migration probabilities estimated using circuit theory and allele distribution models under the two most extreme models of climate change (CCSM_2050_RCP4.5 and CCSM_2070_RCP8.5) for sampled populations of two species of teosintes in Mexico (*Zea mays* spp. *parviglumis* and *Z. mays* spp. *mexicana*).Geographic distances were measured in kilometers and correspond to the the minimum distances to the population with the highest adaptive score (best population) and to the neighboring population with a higher adaptive than the focal population (best neighbor). Except otherwise stated, F-statistics are statistically significant with p-val < 0.01 (d.f. = 21).

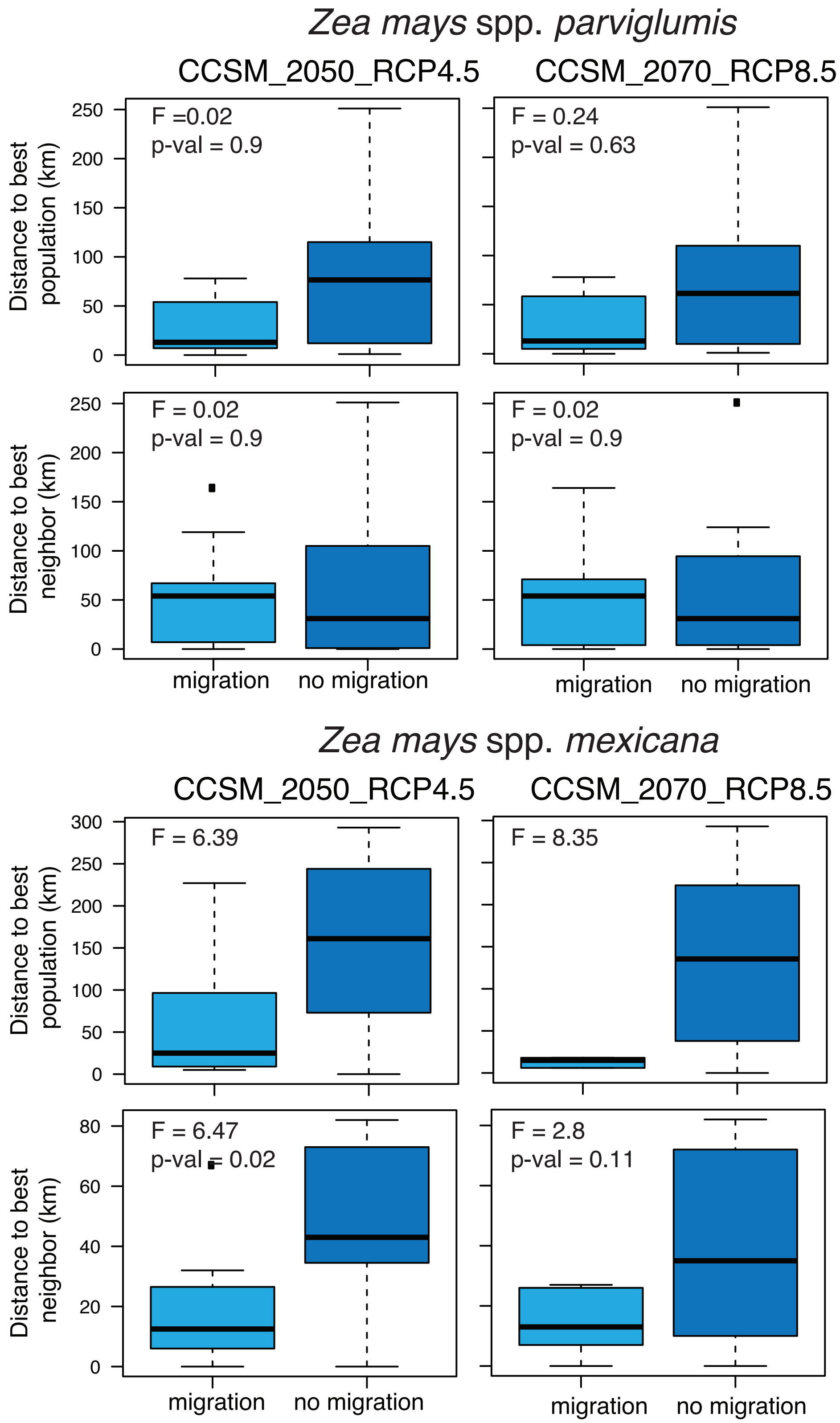

**Supplementary Table 1.** Reduction in the geographical distribution predicted for the two species of teosintes in Mexico under eight different models of climate change: *Zea mays* spp. *mexicana* (*mexicana*) and *Zea mays* spp. *parviglumis* (*parviglumis*). Proportion of known occurrences (ref. 14) and sampled populations within the future predicted distribution for each species. Teosintes occurrences obtained from www.biodiversidad.gob.mx/genes/proyectoMaices.html

| | Year | Climate model | Projected future extent of species relative to the present-day distribution | Proportion of predicted occurrences within the future distribution | Proportion of sampled populations predicted within the future distribution |
|---|---|---|---|---|---|
| *mexicana* | | | | | 23 populations |
| | 2050 | CCSM_4.5 | 83% | 81% | 84% |
| | | CCSM_8.5 | 70% | 67% | 52% |
| | | MIROC_4.5 | 67% | 75% | 72% |
| | | MIROC_8.5 | 60% | 61% | 60% |
| | | **Mean for 2050** | **70%** | **70.9%** | **67%** |
| | 2070 | CCSM_4.5 | 72% | 72% | 68% |
| | | CCSM_8.5 | 57% | 47% | 44% |
| | | MIROC_4.5 | 60% | 53% | 44% |
| | | MIROC_8.5 | 57% | 46% | 40% |
| | | **Mean for 2070** | **61%** | **54.6%** | **49%** |
| *parviglumis* | | | | | 24 populations |
| | 2050 | CCSM_4.5 | 106% | 89% | 88% |
| | | CCSM_8.5 | 102% | 81% | 79% |
| | | MIROC_4.5 | 91% | 88% | 79% |
| | | MIROC_8.5 | 84% | 84% | 83% |
| | | **Mean for 2050** | **96%** | **85.3%** | **82%** |
| | 2070 | CCSM_4.5 | 109% | 85% | 83% |
| | | CCSM_8.5 | 88% | 55% | 50% |
| | | MIROC_4.5 | 92% | 83% | 75% |
| | | MIROC_8.5 | 75% | 62% | 67% |
| | | **Mean for 2070** | **91%** | **71.3%** | **69%** |

**Supplementary Table 2.** Gene annotation for specific putative adaptive SNPs (paSNPs) identified in the two species of teosintes in Mexico: Zea mays spp. mexicana (mexicana) and Zea mays spp. parviglumis (parviglumis). Gene positions were obtained by blasting the chip sequences to the Hapmap2 maize genome reference (Chia et al., 2012). Based on the chip primers, SNPs were identified as being within or near coding regions known for maize. All sequences that were found in a coding regions were annotated against the Phytozome database. Finally, the literature was searched for evidences of local adaptation associated to the putative identified genes (For more details of the methods see ref. 20). According to the annotations, some genes are associated to drought or heat resistance in maize or other plants, while other genes are associated to other stressed related traits. Even if not all SNPs are found to be associated to climate change related traits, we are still finding a strong discriminative power and treating these SNPs as putative adaptive. We believe it is possible that not finding the precise annotations can be explained by the fact that there is a large phylogenetic distance between some species. For instance, an homologous gene of GRMZM2G144985 has been found to be related to phosphorus-impoverished soils response in *Hakea prostrata* (Proteaceae). For a functional validation of these SNPs it will be necessary to perform experimental tests that are outside the scope of the present study.

| | Gene Name | Chip Name | Chromosome | Position | Gene | Associated response | Species | Reference |
|---|---|---|---|---|---|---|---|---|
| mexicana | GRMZM2G141596 | SYN32365 | 9 | 140,846,908 | CONSTANS interacting protein 4 | flowering time | *Zea mays* (Poaceae) | Fjellheim et al. (2014). *Frontiers in plant science*, *5*, 431. |
| mexicana | GRMZM2G144985 | PUT.163a.148928873.366 | 9 | 142,293,132 | long-chain base1 | phosphorus-impoverished soils response | *Hakea prostrata* (Proteaceae) | Kuppusamy et al. (2014). *Plant physiology*, *166*, 1891-1911. |
| mexicana | GRMZM2G145085 | PZE.109096578 | 9 | 142,298,030 | transcription activators;DNA binding;RNA polymerase II transcription factors;catalytics;transcription initiation factors | ABA and drought responses | *Arabidopsis thaliana* (Brassicaceae) | Jin et al. (2011). *New Phytologist*, *190*, 57-74. |
| mexicana | GRMZM2G166780 | SYN27200 | 9 | 139,374,933 | RNA recognition motif in THO complex subunit 4 (THOC4) and similar proteins | abiotic stress | *Oryza sativa* (Poaceae) | Sharma et al (2016). *Frontiers in plant science*, *7*. |
| parviglumis | GRMZM2G063306 | PZE.103059706 | 3 | 112,045,081 | zinc induced facilitator-like 1 | drought tolerance by regulating stomatal closure | *Arabidopsis thaliana* (Brassicaceae) | Remy et al. (2013). *The Plant Cell*, *25*, 901-926. |
| parviglumis | GRMZM2G081928 | PZE.106008139 | 6 | 23,525,249 | Peroxidase superfamily protein | Arsenic stress | *Oryza sativa* (Poaceae) | Yu et al. (2012). *New Phytologist*, *195*, 97-112. |
| parviglumis | GRMZM2G165090 | PZE.101131254 | 1 | 168,430,002 | phytochrome interacting factor 3 | improves drought and salt stress tolerance in rice | *Oryza sativa* (Poaceae) | Gao et al. (2015). *Plant molecular biology*, *87*, 413-428. |

**Supplementary Table 3.** Summary of the genomic offset estimated for species of teosintes in Mexico using different sets of SNPs and eight different models of climate change: Zea mays spp. mexicana (mexicana) and Zea mays spp. parviglumis (parviglumis). For each climatic model we estimated the range and the median of the estimated genomic offset for teosintes occurrences (ref. 14) and sampled populations, separately. Teosintes occurrences obtained from www.biodiversidad.gob.mx/genes/proyectoMaices.html

| SNP dataset | Projected year | Climate model | Range (median) of genomic offset for occurrences | Range (median) of genomic offset for sampled populations | SNP dataset |
|---|---|---|---|---|---|
| *mexicana* | refSNPs | 2050 | CCSM_4.5 | 0.04- 0.07 (0.05) | 0.04-0.08 (0.05) |
| | | | CCSM_8.5 | 0.05- 0.09 (0.06) | 0.05-0.10 (0.06) |
| | | | MIROC_4.5 | 0.04- 0.08 (0.05) | 0.04-0.08 (0.05) |
| | | | MIROC_8.5 | 0.05-0.10 (0.06) | 0.05-0.11 (0.06) |
| | | 2070 | CCSM_4.5 | 0.05-0.08 (0.06) | 0.05-0.09 (0.06) |
| | | | CCSM_8.5 | 0.08-0.13 (0.09) | 0.08-0.13 (0.09) |
| | | | MIROC_4.5 | 0.05-0.12 (0.06) | 0.04-0.12 (0.06) |
| | | | MIROC_8.5 | 0.07- 0.15 (0.08) | 0.07-0.16 (0.09) |
| | canSNPs | 2050 | CCSM_4.5 | 0.04-0.24 (0.06) | 0.03-0.23 (0.08) |
| | | | CCSM_8.5 | 0.06- 0.26 (0.13) | 0.06-0.24 (0.12) |
| | | | MIROC_4.5 | 0.04- 0.29 (0.07) | 0.04-0.25 (0.08) |
| | | | MIROC_8.5 | 0.06- 0.27 (0.11) | 0.05-0.27 (0.11) |
| | | 2070 | CCSM_4.5 | 0.05-0.24 (0.10) | 0.04-0.24 (0.10) |
| | | | CCSM_8.5 | 0.11- 0.28 (0.21) | 0.11-0.30 (0.21) |
| | | | MIROC_4.5 | 0.06- 0.29 (0.11) | 0.05-0.29 (0.11) |
| | | | MIROC_8.5 | 0.08- 0.32 (0.18) | 0.09-0.32 (0.18) |
| | paSNPs | 2050 | CCSM_4.5 | 0.034- 0.88 (0.19) | 0.01-0.90 (0.23) |
| | | | CCSM_8.5 | 0.11- 0.97 (0.39) | 0.10-0.92 (0.37) |
| | | | MIROC_4.5 | 0.035- 0.88 (0.18) | 0.02-0.90 (0.24) |
| | | | MIROC_8.5 | 0.087- 0.93 (0.36) | 0.07-0.93 (0.36) |
| | | 2070 | CCSM_4.5 | 0.077- 0.92 (0.33) | 0.07-0.92 (0.32) |
| | | | CCSM_8.5 | 0.29- 0.99 (0.57) | 0.25-1.00 (0.66) |
| | | | MIROC_4.5 | 0.11- 0.92 (0.36) | 0.07-0.92 (0.36) |
| | | | MIROC_8.5 | 0.19- 0.95 (0.50) | 0.18-0.97 (0.63) |

| | SNP dataset | Projected year | Climate model | Range (median) of genomic offset for occurrences | Range (median) of genomic offset for sampled populations |
|---|---|---|---|---|---|
| *parviglumis* | refSNPs | 2050 | CCSM_4.5 | 0.097- 0.28 (0.16) | 0.09-0.58 (0.14) |
| | | | CCSM_8.5 | 0.13- 0.32 (0.23) | 0.11-0.58 (0.19) |
| | | | MIROC_4.5 | 0.10- 0.19 (0.14) | 0.07-0.50 (0.13) |
| | | | MIROC_8.5 | 0.11-0.28 (0.18) | 0.10-0.62 (0.16) |
| | | 2070 | CCSM_4.5 | 0.12- 0.31 (0.21) | 0.11-0.66 (0.18) |
| | | | CCSM_8.5 | 0.18- 0.43 (0.29) | 0.16-0.59 (0.26) |
| | | | MIROC_4.5 | 0.11- 0.26 (0.18) | 0.10-0.50 (0.17) |
| | | | MIROC_8.5 | 0.15- 0.44 (0.25) | 0.14-0.58 (0.21) |
| | canSNPs | 2050 | CCSM_4.5 | 0.14- 0.27 (0.20) | 0.11-0.44 (0.19) |
| | | | CCSM_8.5 | 0.18-0.36 (0.27) | 0.15-0.57 (0.27) |
| | | | MIROC_4.5 | 0.12- 0.26 (0.17) | 0.11-0.44 (0.18) |
| | | | MIROC_8.5 | 0.16- 0.33 (0.20) | 0.13-0.49 (0.22) |
| | | 2070 | CCSM_4.5 | 0.17-0.34 (0.23) | 0.13-0.54 (0.24) |
| | | | CCSM_8.5 | 0.28- 0.49 (0.38) | 0.24 -0.64 (0.38) |
| | | | MIROC_4.5 | 0.15- 0.36 (0.21) | 0.14-0.45 (0.23) |
| | | | MIROC_8.5 | 0.24- 0.45 (0.29) | 0.18-0.50 (0.33) |
| | paSNPs | 2050 | CCSM_4.5 | 0.21-0.40 (0.28) | 0.16-0.72 (0.30) |
| | | | CCSM_8.5 | 0.33- 0.65 (0.46) | 0.2-0.92 (0.44) |
| | | | MIROC_4.5 | 0.21-0.39 (0.27) | 0.13-0.75 (0.29) |
| | | | MIROC_8.5 | 0.21- 0.47 (0.32) | 0.20-0.69 (0.35) |
| | | 2070 | CCSM_4.5 | 0.24- 0.50 (0.34) | 0.21-0.85 (0.36) |
| | | | CCSM_8.5 | 0.49- 0.75 (0.63) | 0.31- 1.0 (0.6) |
| | | | MIROC_4.5 | 0.23-0.51 (0.32) | 0.19-0.75 (0.35) |
| | | | MIROC_8.5 | 0.38- 0.63 (0.47) | 0.32- 0.77 (0.52) |

**Supplementary Table 4.** Raw values of genomic offset for sampled populations of the two species of teosintes in Mexico using different sets of SNPs and eight different models of climate change**:** Zea mays spp. mexicana (mexicana) and Zea mays spp. parviglumis (parviglumis)

| **Population** | **subespecies** | paSNPs CCSM_4.5_2050 | paSNPs CCSM_8.5_2050 | paSNPs MIROC_4.5_2050 | paSNPs MIROC_8.5_2050 |
|---|---|---|---|---|---|
| Teloloapan_Teloloapan | parviglumis | 0.29 | 0.45 | 0.28 | 0.30 |
| Alcholoa_Teloloapan | parviglumis | 0.28 | 0.44 | 0.28 | 0.28 |
| Teconoapan_Teconoapa | parviglumis | 0.32 | 0.55 | 0.34 | 0.39 |
| Chilpancingo_Chilpancingo | parviglumis | 0.32 | 0.54 | 0.31 | 0.41 |
| Mochitlan_Mochitlan | parviglumis | 0.34 | 0.67 | 0.30 | 0.47 |
| PasoMorelos_Huitzuco | parviglumis | 0.43 | 0.65 | 0.44 | 0.43 |
| Guachinango_Guauchinango | parviglumis | 0.36 | 0.47 | 0.31 | 0.42 |
| Telpitita_VillaPurificacion | parviglumis | 0.43 | 0.69 | 0.43 | 0.62 |
| SMH565_Ejutla | parviglumis | 0.46 | 0.68 | 0.37 | 0.47 |
| SMH577_VilladePurificacion | parviglumis | 0.41 | 0.68 | 0.42 | 0.54 |
| SMH578_Toliman | parviglumis | 0.42 | 0.75 | 0.29 | 0.40 |
| SMHMGCH581_Zitacuaro | parviglumis | 0.35 | 0.43 | 0.37 | 0.43 |
| SanCristobalHonduras_SanJeronim | parviglumis | 0.37 | 0.46 | 0.36 | 0.43 |
| Ahuacatitlan | parviglumis | 0.30 | 0.47 | 0.29 | 0.31 |
| AmatlandeCasas | parviglumis | 0.53 | 0.87 | 0.52 | 0.62 |
| CruceroLagunitas | parviglumis | 0.34 | 0.56 | 0.33 | 0.39 |
| EjutlaA | parviglumis | 0.46 | 0.68 | 0.37 | 0.47 |
| EjutlaB | parviglumis | 0.41 | 0.67 | 0.32 | 0.37 |
| ElRodeo | parviglumis | 0.37 | 0.46 | 0.36 | 0.43 |
| ElSauz | parviglumis | 0.45 | 0.77 | 0.39 | 0.46 |
| LaCadena | parviglumis | 0.49 | 0.63 | 0.51 | 0.63 |
| LaMesa | parviglumis | 0.43 | 0.69 | 0.43 | 0.52 |
| LosGuajes | parviglumis | 0.38 | 0.44 | 0.38 | 0.41 |
| SanLorenzo | parviglumis | 0.34 | 0.61 | 0.33 | 0.42 |
| VillaSeca_Otzolotepec | mexicana | 0.08 | 0.39 | 0.17 | 0.41 |
| Churintzio_Churintzio | mexicana | 0.13 | 0.22 | 0.13 | 0.18 |
| SMH571_Acambaro | mexicana | 0.11 | 0.12 | 0.11 | 0.11 |
| SMH572_Acambaro | mexicana | 0.05 | 0.15 | 0.04 | 0.09 |
| SMH573_Acambaro | mexicana | 0.06 | 0.13 | 0.07 | 0.12 |
| SMH575_SantaAnaMaya | mexicana | 0.09 | 0.19 | 0.09 | 0.15 |
| SMH576_Yuriria | mexicana | 0.19 | 0.30 | 0.19 | 0.25 |
| SMHMGCH579_Puruandiro | mexicana | 0.12 | 0.23 | 0.11 | 0.18 |
| SMHMGCH580_Huandacareo | mexicana | 0.09 | 0.17 | 0.08 | 0.14 |
| SMHMGCH582_JesusMaria | mexicana | 0.38 | 0.40 | 0.37 | 0.38 |
| Cocotitlan_Cocotitlan | mexicana | 0.72 | 0.76 | 0.72 | 0.75 |
| SanNicolas_SnNicolasBuenosAires | mexicana | 0.71 | 0.78 | 0.72 | 0.77 |
| Tenancingo_Tenancingo | mexicana | 0.45 | 0.65 | 0.46 | 0.60 |
| Calpan_Calpan | mexicana | 0.12 | 0.49 | 0.16 | 0.40 |
| Texcoco_Texcoco | mexicana | 0.72 | 0.77 | 0.79 | 0.82 |
| ElPorvenir | mexicana | 0.56 | 0.57 | 0.52 | 0.57 |
| Ixtlan | mexicana | 0.30 | 0.40 | 0.26 | 0.37 |
| Opopeo | mexicana | 0.89 | 0.98 | 0.89 | 0.94 |
| Puruandiro | mexicana | 0.16 | 0.26 | 0.15 | 0.22 |
| SanPedro | mexicana | 0.12 | 0.49 | 0.16 | 0.40 |
| SantaClara | mexicana | 0.89 | 0.98 | 0.89 | 0.94 |
| TenangodelAire | mexicana | 0.04 | 0.17 | 0.05 | 0.11 |
| Xochimilco | mexicana | 0.69 | 0.72 | 0.74 | 0.75 |

| Population | subespecies | canSNPs CCSM_4.5_2050 | canSNPs CCSM_8.5_2050 | canSNPs MIROC_4.5_2050 | canSNPs MIROC_8.5_2050 |
|---|---|---|---|---|---|
| Teloloapan_Teloloapan | parviglumis | 0.22 | 0.29 | 0.22 | 0.23 |
| Alcholoa_Teloloapan | parviglumis | 0.24 | 0.29 | 0.22 | 0.23 |
| Teconoapan_Teconoapa | parviglumis | 0.22 | 0.32 | 0.23 | 0.26 |
| Chilpancingo_Chilpancingo | parviglumis | 0.21 | 0.34 | 0.19 | 0.28 |
| Mochitlan_Mochitlan | parviglumis | 0.32 | 0.48 | 0.26 | 0.38 |
| PasoMorelos_Huitzuco | parviglumis | 0.31 | 0.41 | 0.33 | 0.31 |
| Guachinango_Guauchinango | parviglumis | 0.26 | 0.35 | 0.26 | 0.30 |
| Telpitita_VillaPurificacion | parviglumis | 0.35 | 0.48 | 0.34 | 0.44 |
| SMH565_Ejutla | parviglumis | 0.29 | 0.38 | 0.17 | 0.25 |
| SMH577_VilladePurificacion | parviglumis | 0.34 | 0.48 | 0.34 | 0.41 |
| SMH578_Toliman | parviglumis | 0.28 | 0.48 | 0.21 | 0.23 |
| SMHMGCH581_Zitacuaro | parviglumis | 0.21 | 0.24 | 0.18 | 0.22 |
| SanCristobalHonduras_SanJeronim | parviglumis | 0.19 | 0.26 | 0.17 | 0.21 |
| Ahuacatitlan | parviglumis | 0.23 | 0.30 | 0.22 | 0.25 |
| AmatlandeCasas | parviglumis | 0.27 | 0.41 | 0.26 | 0.29 |
| CruceroLagunitas | parviglumis | 0.24 | 0.33 | 0.23 | 0.26 |
| EjutlaA | parviglumis | 0.29 | 0.38 | 0.17 | 0.25 |
| EjutlaB | parviglumis | 0.36 | 0.46 | 0.25 | 0.31 |
| ElRodeo | parviglumis | 0.19 | 0.26 | 0.17 | 0.21 |
| ElSauz | parviglumis | 0.21 | 0.41 | 0.16 | 0.22 |
| LaCadena | parviglumis | 0.28 | 0.35 | 0.28 | 0.37 |
| LaMesa | parviglumis | 0.22 | 0.36 | 0.20 | 0.27 |
| LosGuajes | parviglumis | 0.26 | 0.31 | 0.27 | 0.28 |
| SanLorenzo | parviglumis | 0.31 | 0.41 | 0.27 | 0.32 |
| VillaSeca_Otzolotepec | mexicana | 0.06 | 0.15 | 0.08 | 0.13 |
| Churintzio_Churintzio | mexicana | 0.06 | 0.08 | 0.07 | 0.07 |
| SMH571_Acambaro | mexicana | 0.05 | 0.06 | 0.04 | 0.06 |
| SMH572_Acambaro | mexicana | 0.05 | 0.07 | 0.04 | 0.06 |
| SMH573_Acambaro | mexicana | 0.04 | 0.06 | 0.04 | 0.06 |
| SMH575_SantaAnaMaya | mexicana | 0.05 | 0.08 | 0.05 | 0.07 |
| SMH576_Yuriria | mexicana | 0.06 | 0.09 | 0.06 | 0.08 |
| SMHMGCH579_Puruandiro | mexicana | 0.06 | 0.09 | 0.05 | 0.07 |
| SMHMGCH580_Huandacareo | mexicana | 0.05 | 0.07 | 0.04 | 0.07 |
| SMHMGCH582_JesusMaria | mexicana | 0.09 | 0.11 | 0.10 | 0.10 |
| Cocotitlan_Cocotitlan | mexicana | 0.19 | 0.21 | 0.20 | 0.21 |
| SanNicolas_SnNicolasBuenosAires | mexicana | 0.19 | 0.21 | 0.20 | 0.21 |
| Tenancingo_Tenancingo | mexicana | 0.13 | 0.19 | 0.14 | 0.18 |
| Calpan_Calpan | mexicana | 0.05 | 0.15 | 0.07 | 0.12 |
| Texcoco_Texcoco | mexicana | 0.20 | 0.21 | 0.21 | 0.22 |
| ElPorvenir | mexicana | 0.17 | 0.20 | 0.15 | 0.17 |
| Ixtlan | mexicana | 0.09 | 0.12 | 0.08 | 0.11 |
| Opopeo | mexicana | 0.24 | 0.26 | 0.29 | 0.27 |
| Puruandiro | mexicana | 0.06 | 0.09 | 0.06 | 0.08 |
| SanPedro | mexicana | 0.05 | 0.15 | 0.07 | 0.12 |
| SantaClara | mexicana | 0.23 | 0.26 | 0.29 | 0.27 |
| TenangodelAire | mexicana | 0.05 | 0.07 | 0.04 | 0.06 |
| Xochimilco | mexicana | 0.18 | 0.19 | 0.19 | 0.20 |

| **Population** | **subespecies** | refSNPs CCSM_4.5_2050 | refSNPs CCSM_8.5_2050 | refSNPs MIROC_4.5_2050 | refSNPs MIROC_8.5_2050 |
|---|---|---|---|---|---|
| Teloloapan_Teloloapan | parviglumis | 0.23 | 0.23 | 0.23 | 0.21 |
| Alcholoa_Teloloapan | parviglumis | 0.16 | 0.18 | 0.16 | 0.15 |
| Teconoapan_Teconoapa | parviglumis | 0.20 | 0.26 | 0.18 | 0.24 |
| Chilpancingo_Chilpancingo | parviglumis | 0.13 | 0.32 | 0.18 | 0.36 |
| Mochitlan_Mochitlan | parviglumis | 0.37 | 0.35 | 0.25 | 0.38 |
| PasoMorelos_Huitzuco | parviglumis | 0.21 | 0.26 | 0.23 | 0.16 |
| Guachinango_Guauchinango | parviglumis | 0.20 | 0.33 | 0.21 | 0.28 |
| Telpitita_VillaPurificacion | parviglumis | 0.34 | 0.42 | 0.25 | 0.32 |
| SMH565_Ejutla | parviglumis | 0.24 | 0.39 | 0.18 | 0.25 |
| SMH577_VilladePurificacion | parviglumis | 0.34 | 0.43 | 0.25 | 0.31 |
| SMH578_Toliman | parviglumis | 0.24 | 0.41 | 0.17 | 0.20 |
| SMHMGCH581_Zitacuaro | parviglumis | 0.20 | 0.23 | 0.18 | 0.22 |
| SanCristobalHonduras_SanJeronim | parviglumis | 0.16 | 0.18 | 0.15 | 0.18 |
| Ahuacatitlan | parviglumis | 0.24 | 0.23 | 0.23 | 0.23 |
| AmatlandeCasas | parviglumis | 0.19 | 0.30 | 0.17 | 0.22 |
| CruceroLagunitas | parviglumis | 0.24 | 0.27 | 0.17 | 0.24 |
| EjutlaA | parviglumis | 0.24 | 0.39 | 0.18 | 0.25 |
| EjutlaB | parviglumis | 0.24 | 0.38 | 0.17 | 0.21 |
| ElRodeo | parviglumis | 0.16 | 0.18 | 0.15 | 0.18 |
| ElSauz | parviglumis | 0.23 | 0.42 | 0.19 | 0.25 |
| LaCadena | parviglumis | 0.14 | 0.19 | 0.14 | 0.37 |
| LaMesa | parviglumis | 0.20 | 0.34 | 0.19 | 0.27 |
| LosGuajes | parviglumis | 0.14 | 0.17 | 0.13 | 0.15 |
| SanLorenzo | parviglumis | 0.21 | 0.33 | 0.18 | 0.23 |
| VillaSeca_Otzolotepec | mexicana | 0.05 | 0.06 | 0.05 | 0.06 |
| Churintzio_Churintzio | mexicana | 0.03 | 0.05 | 0.04 | 0.04 |
| SMH571_Acambaro | mexicana | 0.03 | 0.04 | 0.03 | 0.03 |
| SMH572_Acambaro | mexicana | 0.03 | 0.04 | 0.03 | 0.04 |
| SMH573_Acambaro | mexicana | 0.03 | 0.04 | 0.03 | 0.04 |
| SMH575_SantaAnaMaya | mexicana | 0.03 | 0.04 | 0.03 | 0.04 |
| SMH576_Yuriria | mexicana | 0.04 | 0.05 | 0.04 | 0.04 |
| SMHMGCH579_Puruandiro | mexicana | 0.03 | 0.04 | 0.03 | 0.04 |
| SMHMGCH580_Huandacareo | mexicana | 0.03 | 0.04 | 0.03 | 0.04 |
| SMHMGCH582_JesusMaria | mexicana | 0.03 | 0.04 | 0.04 | 0.04 |
| Cocotitlan_Cocotitlan | mexicana | 0.05 | 0.06 | 0.05 | 0.06 |
| SanNicolas_SnNicolasBuenosAires | mexicana | 0.05 | 0.06 | 0.06 | 0.06 |
| Tenancingo_Tenancingo | mexicana | 0.06 | 0.07 | 0.07 | 0.07 |
| Calpan_Calpan | mexicana | 0.06 | 0.08 | 0.07 | 0.08 |
| Texcoco_Texcoco | mexicana | 0.05 | 0.06 | 0.05 | 0.06 |
| ElPorvenir | mexicana | 0.07 | 0.08 | 0.04 | 0.05 |
| Ixtlan | mexicana | 0.04 | 0.06 | 0.04 | 0.05 |
| Opopeo | mexicana | 0.05 | 0.05 | 0.06 | 0.06 |
| Puruandiro | mexicana | 0.03 | 0.05 | 0.03 | 0.04 |
| SanPedro | mexicana | 0.06 | 0.08 | 0.07 | 0.08 |
| SantaClara | mexicana | 0.04 | 0.05 | 0.06 | 0.06 |
| TenangodelAire | mexicana | 0.06 | 0.07 | 0.07 | 0.07 |
| Xochimilco | mexicana | 0.04 | 0.05 | 0.04 | 0.05 |

| **Population** | **subespecies** | paSNPs CCSM_4.5_2070 | paSNPs CCSM_8.5_2070 | paSNPs MIROC_4.5_2070 | paSNPs MIROC_8.5_2070 |
|---|---|---|---|---|---|
| Teloloapan_Teloloapan | parviglumis | 0.32 | 0.78 | 0.31 | 0.54 |
| Alcholoa_Teloloapan | parviglumis | 0.38 | 0.83 | 0.38 | 0.59 |
| Teconoapan_Teconoapa | parviglumis | 0.42 | 0.74 | 0.40 | 0.50 |
| Chilpancingo_Chilpancingo | parviglumis | 0.41 | 0.87 | 0.39 | 0.60 |
| Mochitlan_Mochitlan | parviglumis | 0.59 | 0.87 | 0.42 | 0.73 |
| PasoMorelos_Huitzuco | parviglumis | 0.48 | 0.87 | 0.42 | 0.71 |
| Guachinango_Guauchinango | parviglumis | 0.44 | 0.65 | 0.41 | 0.57 |
| Telpitita_VillaPurificacion | parviglumis | 0.51 | 0.96 | 0.68 | 0.83 |
| SMH565_Ejutla | parviglumis | 0.53 | 0.84 | 0.46 | 0.67 |
| SMH577_VilladePurificacion | parviglumis | 0.46 | 0.94 | 0.64 | 0.82 |
| SMH578_Toliman | parviglumis | 0.67 | 0.95 | 0.43 | 0.60 |
| SMHMGCH581_Zitacuaro | parviglumis | 0.43 | 0.68 | 0.43 | 0.61 |
| SanCristobalHonduras_SanJeronim | parviglumis | 0.43 | 0.65 | 0.45 | 0.55 |
| Ahuacatitlan | parviglumis | 0.32 | 0.83 | 0.31 | 0.61 |
| AmatlandeCasas | parviglumis | 0.58 | 1.00 | 0.56 | 0.84 |
| CruceroLagunitas | parviglumis | 0.43 | 0.85 | 0.40 | 0.50 |
| EjutlaA | parviglumis | 0.53 | 0.84 | 0.46 | 0.67 |
| EjutlaB | parviglumis | 0.46 | 0.87 | 0.48 | 0.63 |
| ElRodeo | parviglumis | 0.43 | 0.65 | 0.45 | 0.55 |
| ElSauz | parviglumis | 0.49 | 0.99 | 0.44 | 0.66 |
| LaCadena | parviglumis | 0.55 | 0.80 | 0.60 | 0.77 |
| LaMesa | parviglumis | 0.47 | 0.92 | 0.47 | 0.73 |
| LosGuajes | parviglumis | 0.41 | 0.67 | 0.40 | 0.62 |
| SanLorenzo | parviglumis | 0.43 | 0.82 | 0.41 | 0.68 |
| VillaSeca_Otzolotepec | mexicana | 0.30 | 0.78 | 0.36 | 0.73 |
| Churintzio_Churintzio | mexicana | 0.19 | 0.45 | 0.21 | 0.28 |
| SMH571_Acambaro | mexicana | 0.10 | 0.30 | 0.12 | 0.19 |
| SMH572_Acambaro | mexicana | 0.30 | 0.33 | 0.30 | 0.35 |
| SMH573_Acambaro | mexicana | 0.13 | 0.37 | 0.31 | 0.36 |
| SMH575_SantaAnaMaya | mexicana | 0.19 | 0.36 | 0.31 | 0.39 |
| SMH576_Yuriria | mexicana | 0.30 | 0.56 | 0.30 | 0.38 |
| SMHMGCH579_Puruandiro | mexicana | 0.23 | 0.48 | 0.17 | 0.31 |
| SMHMGCH580_Huandacareo | mexicana | 0.18 | 0.43 | 0.18 | 0.38 |
| SMHMGCH582_JesusMaria | mexicana | 0.40 | 0.51 | 0.40 | 0.41 |
| Cocotitlan_Cocotitlan | mexicana | 0.74 | 0.94 | 0.76 | 0.95 |
| SanNicolas_SnNicolasBuenosAires | mexicana | 0.77 | 0.99 | 0.78 | 0.96 |
| Tenancingo_Tenancingo | mexicana | 0.60 | 0.80 | 0.63 | 0.78 |
| Calpan_Calpan | mexicana | 0.44 | 0.79 | 0.42 | 0.77 |
| Texcoco_Texcoco | mexicana | 0.74 | 0.89 | 0.82 | 0.83 |
| ElPorvenir | mexicana | 0.65 | 0.68 | 0.63 | 0.65 |
| Ixtlan | mexicana | 0.33 | 0.58 | 0.37 | 0.43 |
| Opopeo | mexicana | 0.92 | 1.00 | 0.93 | 0.95 |
| Puruandiro | mexicana | 0.27 | 0.52 | 0.26 | 0.35 |
| SanPedro | mexicana | 0.44 | 0.79 | 0.42 | 0.77 |
| SantaClara | mexicana | 0.93 | 1.00 | 0.92 | 0.95 |
| TenangodelAire | mexicana | 0.08 | 0.71 | 0.18 | 0.61 |
| Xochimilco | mexicana | 0.68 | 0.95 | 0.76 | 0.93 |

| **Population** | **subespecies** | canSNPs CCSM_4.5_2070 | canSNPs CCSM_8.5_2070 | canSNPs MIROC_4.5_2070 | canSNPs MIROC_8.5_2070 |
|---|---|---|---|---|---|
| Teloloapan_Teloloapan | parviglumis | 0.26 | 0.48 | 0.24 | 0.35 |
| Alcholoa_Teloloapan | parviglumis | 0.30 | 0.53 | 0.29 | 0.38 |
| Teconoapan_Teconoapa | parviglumis | 0.29 | 0.46 | 0.27 | 0.33 |
| Chilpancingo_Chilpancingo | parviglumis | 0.28 | 0.57 | 0.26 | 0.40 |
| Mochitlan_Mochitlan | parviglumis | 0.44 | 0.61 | 0.35 | 0.50 |
| PasoMorelos_Huitzuco | parviglumis | 0.36 | 0.56 | 0.30 | 0.45 |
| Guachinango_Guauchinango | parviglumis | 0.32 | 0.46 | 0.30 | 0.42 |
| Telpitita_VillaPurificacion | parviglumis | 0.40 | 0.64 | 0.48 | 0.59 |
| SMH565_Ejutla | parviglumis | 0.35 | 0.50 | 0.28 | 0.37 |
| SMH577_VilladePurificacion | parviglumis | 0.38 | 0.65 | 0.47 | 0.60 |
| SMH578_Toliman | parviglumis | 0.45 | 0.58 | 0.24 | 0.36 |
| SMHMGCH581_Zitacuaro | parviglumis | 0.24 | 0.39 | 0.23 | 0.33 |
| SanCristobalHonduras_SanJeronim | parviglumis | 0.23 | 0.38 | 0.21 | 0.33 |
| Ahuacatitlan | parviglumis | 0.25 | 0.51 | 0.25 | 0.37 |
| AmatlandeCasas | parviglumis | 0.30 | 0.53 | 0.29 | 0.46 |
| CruceroLagunitas | parviglumis | 0.29 | 0.49 | 0.27 | 0.33 |
| EjutlaA | parviglumis | 0.35 | 0.50 | 0.28 | 0.37 |
| EjutlaB | parviglumis | 0.40 | 0.61 | 0.37 | 0.45 |
| ElRodeo | parviglumis | 0.23 | 0.38 | 0.21 | 0.33 |
| ElSauz | parviglumis | 0.27 | 0.54 | 0.22 | 0.32 |
| LaCadena | parviglumis | 0.33 | 0.46 | 0.34 | 0.45 |
| LaMesa | parviglumis | 0.28 | 0.51 | 0.25 | 0.40 |
| LosGuajes | parviglumis | 0.29 | 0.49 | 0.28 | 0.45 |
| SanLorenzo | parviglumis | 0.35 | 0.57 | 0.32 | 0.48 |
| VillaSeca_Otzolotepec | mexicana | 0.09 | 0.24 | 0.10 | 0.21 |
| Churintzio_Churintzio | mexicana | 0.07 | 0.15 | 0.09 | 0.11 |
| SMH571_Acambaro | mexicana | 0.06 | 0.11 | 0.06 | 0.08 |
| SMH572_Acambaro | mexicana | 0.08 | 0.13 | 0.09 | 0.11 |
| SMH573_Acambaro | mexicana | 0.06 | 0.13 | 0.09 | 0.11 |
| SMH575_SantaAnaMaya | mexicana | 0.07 | 0.14 | 0.09 | 0.12 |
| SMH576_Yuriria | mexicana | 0.09 | 0.17 | 0.09 | 0.12 |
| SMHMGCH579_Puruandiro | mexicana | 0.08 | 0.16 | 0.07 | 0.11 |
| SMHMGCH580_Huandacareo | mexicana | 0.07 | 0.15 | 0.07 | 0.11 |
| SMHMGCH582_JesusMaria | mexicana | 0.10 | 0.15 | 0.12 | 0.13 |
| Cocotitlan_Cocotitlan | mexicana | 0.21 | 0.26 | 0.21 | 0.26 |
| SanNicolas_SnNicolasBuenosAires | mexicana | 0.21 | 0.27 | 0.21 | 0.27 |
| Tenancingo_Tenancingo | mexicana | 0.17 | 0.23 | 0.19 | 0.22 |
| Calpan_Calpan | mexicana | 0.14 | 0.23 | 0.13 | 0.22 |
| Texcoco_Texcoco | mexicana | 0.20 | 0.26 | 0.22 | 0.25 |
| ElPorvenir | mexicana | 0.20 | 0.24 | 0.18 | 0.22 |
| Ixtlan | mexicana | 0.10 | 0.18 | 0.11 | 0.14 |
| Opopeo | mexicana | 0.24 | 0.28 | 0.29 | 0.32 |
| Puruandiro | mexicana | 0.08 | 0.16 | 0.08 | 0.11 |
| SanPedro | mexicana | 0.14 | 0.23 | 0.13 | 0.22 |
| SantaClara | mexicana | 0.24 | 0.29 | 0.29 | 0.33 |
| TenangodelAire | mexicana | 0.06 | 0.21 | 0.08 | 0.19 |
| Xochimilco | mexicana | 0.18 | 0.26 | 0.20 | 0.25 |

| **Population** | **subespecies** | refSNPs CCSM_4.5_2070 | refSNPs CCSM_8.5_2070 | refSNPs MIROC_4.5_2070 | refSNPs MIROC_8.5_2070 |
|---|---|---|---|---|---|
| Teloloapan_Teloloapan | parviglumis | 0.26 | 0.32 | 0.22 | 0.28 |
| Alcholoa_Teloloapan | parviglumis | 0.20 | 0.29 | 0.19 | 0.25 |
| Teconoapan_Teconoapa | parviglumis | 0.31 | 0.36 | 0.21 | 0.22 |
| Chilpancingo_Chilpancingo | parviglumis | 0.20 | 0.32 | 0.30 | 0.29 |
| Mochitlan_Mochitlan | parviglumis | 0.42 | 0.46 | 0.31 | 0.30 |
| PasoMorelos_Huitzuco | parviglumis | 0.25 | 0.33 | 0.16 | 0.25 |
| Guachinango_Guauchinango | parviglumis | 0.28 | 0.45 | 0.29 | 0.44 |
| Telpitita_VillaPurificacion | parviglumis | 0.38 | 0.55 | 0.32 | 0.55 |
| SMH565_Ejutla | parviglumis | 0.33 | 0.48 | 0.24 | 0.39 |
| SMH577_VilladePurificacion | parviglumis | 0.37 | 0.57 | 0.33 | 0.58 |
| SMH578_Toliman | parviglumis | 0.35 | 0.55 | 0.21 | 0.41 |
| SMHMGCH581_Zitacuaro | parviglumis | 0.25 | 0.32 | 0.22 | 0.26 |
| SanCristobalHonduras_SanJeronim | parviglumis | 0.17 | 0.24 | 0.16 | 0.22 |
| Ahuacatitlan | parviglumis | 0.24 | 0.32 | 0.22 | 0.29 |
| AmatlandeCasas | parviglumis | 0.23 | 0.38 | 0.24 | 0.37 |
| CruceroLagunitas | parviglumis | 0.32 | 0.39 | 0.22 | 0.22 |
| EjutlaA | parviglumis | 0.33 | 0.48 | 0.24 | 0.39 |
| EjutlaB | parviglumis | 0.30 | 0.49 | 0.26 | 0.38 |
| ElRodeo | parviglumis | 0.17 | 0.24 | 0.16 | 0.22 |
| ElSauz | parviglumis | 0.31 | 0.49 | 0.26 | 0.38 |
| LaCadena | parviglumis | 0.17 | 0.26 | 0.35 | 0.39 |
| LaMesa | parviglumis | 0.27 | 0.45 | 0.24 | 0.39 |
| LosGuajes | parviglumis | 0.17 | 0.26 | 0.15 | 0.20 |
| SanLorenzo | parviglumis | 0.29 | 0.45 | 0.25 | 0.40 |
| VillaSeca_Otzolotepec | mexicana | 0.06 | 0.09 | 0.06 | 0.09 |
| Churintzio_Churintzio | mexicana | 0.04 | 0.08 | 0.05 | 0.06 |
| SMH571_Acambaro | mexicana | 0.03 | 0.06 | 0.04 | 0.05 |
| SMH572_Acambaro | mexicana | 0.04 | 0.07 | 0.04 | 0.05 |
| SMH573_Acambaro | mexicana | 0.04 | 0.07 | 0.04 | 0.06 |
| SMH575_SantaAnaMaya | mexicana | 0.04 | 0.07 | 0.04 | 0.06 |
| SMH576_Yuriria | mexicana | 0.05 | 0.08 | 0.05 | 0.06 |
| SMHMGCH579_Puruandiro | mexicana | 0.04 | 0.08 | 0.04 | 0.06 |
| SMHMGCH580_Huandacareo | mexicana | 0.04 | 0.07 | 0.04 | 0.05 |
| SMHMGCH582_JesusMaria | mexicana | 0.04 | 0.07 | 0.05 | 0.06 |
| Cocotitlan_Cocotitlan | mexicana | 0.06 | 0.08 | 0.06 | 0.08 |
| SanNicolas_SnNicolasBuenosAires | mexicana | 0.06 | 0.08 | 0.06 | 0.08 |
| Tenancingo_Tenancingo | mexicana | 0.07 | 0.09 | 0.08 | 0.09 |
| Calpan_Calpan | mexicana | 0.07 | 0.10 | 0.08 | 0.10 |
| Texcoco_Texcoco | mexicana | 0.06 | 0.08 | 0.06 | 0.08 |
| ElPorvenir | mexicana | 0.08 | 0.10 | 0.05 | 0.07 |
| Ixtlan | mexicana | 0.05 | 0.08 | 0.05 | 0.07 |
| Opopeo | mexicana | 0.05 | 0.07 | 0.06 | 0.08 |
| Puruandiro | mexicana | 0.04 | 0.08 | 0.04 | 0.06 |
| SanPedro | mexicana | 0.07 | 0.10 | 0.08 | 0.10 |
| SantaClara | mexicana | 0.05 | 0.07 | 0.06 | 0.08 |
| TenangodelAire | mexicana | 0.07 | 0.10 | 0.07 | 0.09 |
| Xochimilco | mexicana | 0.05 | 0.07 | 0.05 | 0.07 |

**Supplementary Table 5.** Climatic variables associated with the allele frequencies of different sets of SNPs identified in the two species of teosintes in Mexico: Zea mays spp. mexicana (mexicana) and Zea mays spp. parviglumis (parviglumis). Variables with the highest weighted importance for model construction and proportion of SNPs with significant predictive power (range and median of R2 values). refSNPs: references SNPs, canSNPs: candidate SNPs; paSNPs: putative adaptive SNPs. Blioclimatic layer were obtained from www.worldclim.org

| | **SNP dataset** | **Bioclimatic variables*** | **Proportion of SNPs with significant predictive power** | **Range (median) of $R^2$** |
|---|---|---|---|---|
| *mexicana* | refSNPs | 4, 5, 8, 10, 16 | 28% | 0.002-0.6 (0.17) |
| | canSNPs | 9, 10, 13, 12, 16 | 100% | 0.10-0.88 (0.36) |
| | paSNPs | 1, 4, 5, 9, 10 | 100% | 0.07-0.92 (0.87) |
| *mexicana* | refSNPs | 2, 3, 17, 18, 19 | 63% | 0.001-0.87 (0.26) |
| | canSNPs | 4, 12, 16, 18, 19 | 96% | 0.01-0.61 (0.32) |
| | paSNPs | 6, 11, 12, 16, 18 | 100% | 0.12-0.6 (0.46) |

Variable names after ref. 53.

BIO1 Annual Mean Temperature; BIO2 Mean Diurnal Range (Mean of monthly (max temp - min temp)); BIO3 Isothermality (BIO2/BIO7) (* 100); BIO4 Temperature Seasonality (standard deviation *100); BIO5 Max Temperature of Warmest Month; BIO6 Min Temperature of Coldest Month; BIO7 Temperature Annual Range (BIO5-BIO6); BIO8 Mean Temperature of Wettest Quarter; BIO9 Mean Temperature of Driest Quarter; BIO10 Mean Temperature of Warmest Quarter; BIO11 Mean Temperature of Coldest Quarter; BIO12 Annual Precipitation; BIO13 Precipitation of Wettest Month; BIO14 Precipitation of Driest Month; BIO15 Precipitation Seasonality (Coefficient of Variation); BIO16 Precipitation of Wettest Quarter; BIO17 Precipitation of Driest Quarter; BIO18 Precipitation of Warmest Quarter; BIO19 Precipitation of Coldest Quarter

**Supplementary Table 6.** Reduction in the geographical distribution predicted for putative adaptive SNPs (paSNPs) in the two species of teosintes in Mexico under eight different models of climate change: Zea mays spp. mexicana (mexicana) and Zea mays spp. parviglumis (parviglumis). Proportion of known occurrences (ref. 14) and sampled populations within the predicted future distribution for paSNPs. Teosintes occurrences obtained from www.biodiversidad.gob.mx/genes/proyectoMaices.html

| Subspecies | Year | Climate model | Projected future extent of species relative to the present-day [illegible] | Proportion of predicted occurrences within the future distribution | Proportion of sampled populations predicted within the future distribution |
|---|---|---|---|---|---|
| *mexicana* | 2050 | CCSM_4.5 | 108% | 14% | 22% |
| | | CCSM_8.5 | 44% | 2% | 0% |
| | | MIROC_4.5 | 35% | 2% | 0% |
| | | MIROC_8.5 | 33% | 6% | 0% |
| | | **Mean for 2050** | **54.9%** | **6%** | **5%** |
| | 2070 | CCSM_4.5 | 91% | 14% | 17% |
| | | CCSM_8.5 | 15% | 1% | 0% |
| | | MIROC_4.5 | 46% | 6% | 9% |
| | | MIROC_8.5 | 3% | 0% | 0% |
| | | **Mean for 2070** | **38.8%** | **5%** | **6%** |
| *parviglumis* | 2050 | CCSM_4.5 | 73% | 26% | 25% |
| | | CCSM_8.5 | 59% | 14% | 17% |
| | | MIROC_4.5 | 73% | 31% | 17% |
| | | MIROC_8.5 | 52% | 14% | 8% |
| | | **Mean for 2050** | **64.4%** | **21%** | **17%** |
| | 2070 | CCSM_4.5 | 66% | 24% | 21% |
| | | CCSM_8.5 | 36% | 5% | 0% |
| | | MIROC_4.5 | 66% | 26% | 25% |
| | | MIROC_8.5 | 47% | 10% | 8% |
| | | **Mean for 2070** | **53.8%** | **16%** | **14%** |

**Supplementary Table 7.** Migration potential for sampled populations of two species of teosintes in Mexico: *Zea mays* spp. *mexicana* (*mexicana*) and *Zea mays* spp. *parviglumis* (*parviglumis*) under the two most extreme models of climate change: CCSM_2050_RCP4.5 and CCSM_2070_RCP8.5. Minimum geographic distances to the populations with highest frequency of putative adaptive SNPs. Teosintes occurrences obtained from www.biodiversidad.gob.mx/genes/proyectoMaices.html

| Species | Population | Adaptive score | Migration potential CCSM_2050_RCP4.5 | Migration potential CCSM_2070_RCP8.5 | Minimum distance to best population (km) | Adaptive score for best population | Minimum distance to best neighbor (km) | Adaptive score for best neighbor |
|---|---|---|---|---|---|---|---|---|
| mexicana | Calpan_Calpan | 0.21 | no migration | no migration | 266 | 0.79 | 43 | 0.22 |
| mexicana | Churintzio_Churintzio | 0.82 | migration | no migration | 32 | 0.87 | 32 | 0.87 |
| mexicana | Cocotitlan_Cocotitlan | 0.22 | no migration | no migration | 223 | 0.79 | — | Highest adaptive score locally |
| mexicana | ElPorvenir | 0.42 | no migration | no migration | 38 | 0.79 | 38 | 0.79 |
| mexicana | Ixtlan | 0.87 | no migration | no migration | — | Highest adaptive score in species | — | Highest adaptive score locally |
| mexicana | Opopeo | 0.37 | no migration | no migration | 72 | 0.72 | 72 | 0.72 |
| mexicana | Puruandiro | 0.7 | migration | migration | 6 | 0.7 | 13 | 0.75 |
| mexicana | SanNicolas_SnNicolasBuenosAires | 0.2 | no migration | no migration | 293 | 0.79 | 72 | 0.22 |
| mexicana | SanPedro | 0.18 | no migration | no migration | 265 | 0.79 | 42 | 0.22 |
| mexicana | SantaClara | 0.31 | no migration | no migration | 74 | 0.72 | 74 | 0.72 |
| mexicana | SMH571_Acambaro | 0.74 | migration | no migration | 5 | 0.79 | 5 | 0.79 |
| mexicana | SMH572_Acambaro | 0.79 | migration | no migration | 129 | 0.82 | — | Highest adaptive score locally |
| mexicana | SMH573_Acambaro | 0.75 | migration | no migration | 12 | 0.79 | 12 | 0.79 |
| mexicana | SMH575_SantaAnaMaya | 0.74 | migration | migration | 15 | 0.75 | 27 | 0.79 |
| mexicana | SMH576_Yuriria | 0.75 | migration | migration | 47 | 0.75 | — | Highest adaptive score locally |
| mexicana | SMHMGCH579_Puruandiro | 0.7 | migration | migration | 6 | 0.7 | 7 | 0.75 |
| mexicana | SMHMGCH580_Huandacareo | 0.72 | migration | migration | 18 | 0.74 | 25 | 0.75 |
| mexicana | SMHMGCH582_JesusMaria | 0.6 | migration | no migration | 64 | 0.82 | 67 | 0.87 |
| mexicana | Tenancingo_Tenancingo | 0.15 | migration | no migration | 227 | 0.79 | 10 | 0.22 |
| mexicana | TenangodelAire | 0.12 | no migration | no migration | 161 | 0.79 | 77 | 0.22 |

| | | | | | | | | |
|---|---|---|---|---|---|---|---|---|
| mexicana | Texcoco_Texcoco | 0.2 | no migration | no migration | 209 | 0.79 | 31 | 0.22 |
| mexicana | VillaSeca_Otzolotepec | 0.18 | no migration | no migration | 142 | 0.79 | 82 | 0.22 |
| mexicana | Xochimilco | 0.13 | migration | no migration | 200 | 0.79 | 24 | 0.22 |
| parviglumis | Ahuacatitlan | 0.71 | migration | migration | 3 | 0.72 | 64 | 0.78 |
| parviglumis | Alcholoa_Teloloapan | 0.63 | migration | migration | 10 | 0.72 | 75 | 0.78 |
| parviglumis | AmatlandeCasas | 0.3 | no migration | no migration | 105 | 0.61 | 105 | 0.61 |
| parviglumis | Chilpancingo_Chilpancingo | 0.65 | migration | migration | 50 | 0.97 | 50 | 0.97 |
| parviglumis | CruceroLagunitas | 0.96 | migration | migration | 1 | 0.97 | 1 | 0.97 |
| parviglumis | EjutlaA | 0.61 | no migration | no migration | 69 | 0.8 | — | Highest adaptive score locally |
| parviglumis | EjutlaB | 0.6 | migration | no migration | 7 | 0.61 | 7 | 0.61 |
| parviglumis | ElSauz | 0.37 | migration | no migration | 54 | 0.6 | 55 | 0.61 |
| parviglumis | Guachinango_Guauchinango | 0.35 | no migration | no migration | 84 | 0.61 | 84 | 0.61 |
| parviglumis | LaCadena | 0.6 | migration | migration | 78 | 0.6 | 164 | 0.72 |
| parviglumis | LaMesa | 0.16 | no migration | no migration | 8 | 0.6 | 15 | 0.61 |
| parviglumis | LosGuajes | 0.6 | migration | migration | 13 | 0.61 | 119 | 0.72 |
| parviglumis | Mochitlan_Mochitlan | 0.56 | migration | migration | 14 | 0.65 | 54 | 0.97 |
| parviglumis | PasoMorelos_Huitzuco | 0.78 | no migration | no migration | 139 | 0.97 | — | Highest adaptive score locally |
| parviglumis | SanCristobalHonduras_SanJeronimoCoatlan | 0.65 | no migration | no migration | 251 | 0.96 | 251 | 0.97 |
| parviglumis | SanLorenzo | 0.23 | no migration | no migration | 12 | 0.6 | 20 | 0.61 |
| parviglumis | SMH565_Ejutla | 0.49 | no migration | no migration | 1 | 0.61 | 1 | 0.61 |
| parviglumis | SMH577_VilladePurificacion | 0.86 | migration | migration | 664 | 0.97 | — | Highest adaptive score locally |
| parviglumis | SMH578_Toliman | 0.26 | no migration | no migration | 42 | 0.6 | 42 | 0.61 |
| parviglumis | SMHMGCH581_Zitacuaro | 0.61 | no migration | no migration | 115 | 0.63 | 124 | 0.72 |

| | | | | | | | | |
|---|---|---|---|---|---|---|---|---|
| parviglumis | Teconoapan_Teconoapa | 0.97 | migration | migration | — | Highest adaptive score in species | — | Highest adaptive score locally |
| parviglumis | Teloloapan_Teloloapan | 0.72 | migration | migration | 67 | 0.78 | 67 | 0.78 |
| parviglumis | Telpitita_VillaPurificacion | 0.8 | migration | migration | 7 | 0.86 | 7 | 0.86 |

**Supplementary Table 8**. Geographical coordinates, genotype for putative adaptive SNPs, and estimated adaptive scores for maize landraces accessions in Mexico. Detailed information on landrace and known occurrences can be obtained at www.biodiversidad.gob.mx/usos/maices/maiz.html

| Landrace | Longitude | Latitude | PZE.100001195 | PZE.108035280 | PZE.109096578 | SYN27200 | SYN32365 | SYN37470 | PZE.101131254 | PZE.103059706 | PZE.104026211 | PZE.104044827 | PZE.105090166 | PZE.106008139 | PZE.109029456 | PZE.104055811 | Adaptive score |
|---|---|---|---|---|---|---|---|---|---|---|---|---|---|---|---|---|---|
| **Ancho** | -98.77861111 | 18.9975 | GG | AA | AA | CA | GA | GG | AA | AC | CC | GA | GG | GG | AG | GG | 0.4545454545 |
| **Ancho** | -98.80944444 | 18.96805556 | AA | GG | AA | AA | AA | GG | CA | CC | AC | GA | GA | GG | GG | GA | 0.5909090909 |
| **Ancho** | -98.77861111 | 18.9975 | AA | GG | AA | CA | GG | AA | CC | AC | CC | GA | GG | GG | AA | GA | 0.6363636364 |
| **Apachito** | -107.583 | 28.68577778 | GG | AA | AA | AA | AA | AG | AA | AC | CC | AA | GA | GG | AA | AA | 0.6818181818 |
| **Apachito** | -108.0796111 | 29.11311111 | GG | GA | AA | AA | AA | AG | AA | AC | CC | GA | GA | AG | AG | GA | 0.6363636364 |
| **Arrocillo** | -97.98080556 | 19.93408333 | GG | AA | AA | CA | GA | AG | AA | AC | CC | AA | GG | AA | AA | GG | 0.3636363636 |
| **Arrocillo** | -97.97941667 | 19.95577778 | AG | AA | AA | AA | GA | AA | AA | CC | AC | AA | AA | GG | GG | GG | 0.5454545455 |
| **Arrocillo** | -97.98033333 | 19.94305556 | AG | GA | AA | CA | AA | AG | CA | AC | AC | AA | AA | AG | AG | GA | 0.5909090909 |
| **Arrocillo** | -97.97941667 | 19.95577778 | AG | AA | AA | CA | GA | AG | AA | CC | AC | GA | AA | AG | AG | GG | 0.4545454545 |
| **Azul** | -108.0152222 | 28.45538889 | GG | GA | AA | AA | AA | AA | AA | CC | CC | AA | GG | GG | GG | AA | 0.6363636364 |
| **Azul** | -108.0157778 | 28.45730556 | AG | AA | AA | CC | AA | AA | AA | CC | CC | GA | GA | GG | GG | AA | 0.6363636364 |
| **Blando de Sonora** | -108.3833333 | 26.39472222 | GG | GG | AA | AA | AA | AA | AA | AA | CC | GG | AA | AG | AG | AA | 0.8636363636 |
| **Bofo** | -108.0082222 | 27.43477778 | AA | GA | AA | AA | GA | GG | AA | CC | AC | AA | AA | AA | AA | GA | 0.4090909091 |
| **Cacahuacintle** | -99.50888889 | 19.19638889 | AG | GA | AA | AA | GA | GG | AA | AC | CC | GA | GA | AG | AG | AA | 0.5909090909 |
| **Cacahuacintle** | -99.61805556 | 19.1675 | GG | 0 | AA | AA | GA | GG | AA | AC | CC | GG | GA | GG | AG | AA | 0.6818181818 |
| **Cacahuacintle** | -98.05666667 | 19.59972222 | GG | GA | AA | AA | AA | GG | CA | AC | CC | AA | GA | GG | AA | GG | 0.5909090909 |
| **Cacahuacintle** | -98.43111111 | 20.33916667 | GG | GA | AA | AA | AA | AA | CA | CC | CC | GA | AA | AA | GG | AA | 0.7272727273 |
| **Cacahuacintle** | -97.90083333 | 19.22194444 | GG | AA | AA | CC | GA | AG | AA | CC | CC | GA | GA | AG | GG | GG | 0.4090909091 |
| **Celaya** | -101.1188889 | 20.09111111 | GG | AA | AA | CA | AA | AG | AA | AC | CC | GA | GA | AA | AA | AA | 0.5909090909 |
| **Celaya** | -101.4052778 | 21.00861111 | AG | AA | AA | CC | AA | AG | AA | CC | AC | AA | AA | AA | GG | GA | 0.4090909091 |
| **Celaya** | -101.1611111 | 20.37416667 | AG | AA | AA | CA | GA | GG | AA | AC | CC | GA | AA | AG | GG | GG | 0.5 |
| **Chalqueño** | -97.92944444 | 19.29083333 | AA | AA | AA | CC | GG | AG | AA | CC | CC | AA | GG | AG | AA | AA | 0.3636363636 |
| **Chalqueño** | -100.2275 | 18.66027778 | GG | AA | AA | CC | AA | AG | CA | AC | CC | GA | GA | AG | AG | GG | 0.5454545455 |
| **Chalqueño** | -97.98805556 | 19.35916667 | GG | AA | AA | CA | GA | AA | AA | CC | CC | GA | AA | GG | AA | GA | 0.6363636364 |
| **Chalqueño** | -99.59527778 | 19.41808889 | AG | GG | AA | CA | AA | GG | AA | AA | AC | GG | AA | AG | AG | GA | 0.6363636364 |
| **Chalqueño** | -98.33722222 | 20.22138889 | AG | GA | GA | CA | GA | AG | AA | CC | CC | AA | GA | GG | AG | AA | 0.5 |
| **Chalqueño** | -98.28055556 | 20.02972222 | AG | GG | AA | CA | AA | GG | CA | CC | CC | GA | GG | AG | GG | GA | 0.5 |
| **Chalqueño** | -98.80388889 | 19.10361111 | GG | AA | AA | CC | GA | AG | AA | AC | CC | AA | GG | GG | AG | GG | 0.4090909091 |
| **Chapalote** | -109.2996667 | 29.90441667 | GG | AA | AA | AA | GG | AG | AA | AC | CC | AA | AA | AG | AG | GG | 0.5 |
| **Chapalote** | -109.6742778 | 29.80538889 | GG | GG | AA | AA | AA | AG | AA | AC | CC | GA | GG | AG | GG | GG | 0.5454545455 |
| **Cónico** | -98.04833333 | 19.51888889 | GG | AA | AA | AA | AA | AG | AA | CC | CC | AA | GA | GG | AG | AA | 0.6363636364 |
| **Cónico** | -99.50972222 | 19.18472222 | GG | AA | AA | AA | AA | AG | AA | CC | CC | GA | GG | GG | AA | GA | 0.5909090909 |
| **Cónico** | -98.37861111 | 20.33166667 | AG | AA | AA | CC | AA | AG | AA | CC | AC | AA | AA | AA | GG | GA | 0.4090909091 |
| **Cónico** | -97.65222222 | 19.31916667 | GG | GA | AA | AA | AA | AG | AA | AA | CC | AA | GG | AG | AG | GA | 0.5909090909 |
| **Cónico** | -100.0627778 | 19.46055556 | AG | GG | AA | AA | GA | GG | AA | CC | CC | AA | GA | GG | AA | GA | 0.5 |
| **Cónico** | -98.29111111 | 20.24333333 | AA | GA | AA | AA | GA | GG | CA | AC | CC | GA | AA | GG | AG | AA | 0.7272727273 |
| **Cónico** | -98.405 | 20.28666667 | AA | AA | AA | CA | GG | AG | AA | CC | CC | GA | AA | AG | GG | GA | 0.5 |
| **Cónico** | -98.37388889 | 20.35888889 | AG | AA | AA | CA | GA | AA | AA | CC | CC | AA | GA | GG | AG | AA | 0.5909090909 |
| **Cónico** | -98.78166667 | 19.09055556 | GG | AA | AA | AA | GA | AG | AA | CC | CC | AA | GA | GG | AG | GG | 0.5 |
| **Cónico** | -98.64722222 | 19.75944444 | AG | AA | AA | CA | GA | GG | AA | AC | CC | AA | AA | AA | GG | GG | 0.4090909091 |
| **Cónico** | -99.95611111 | 19.65111111 | AG | AA | AA | CC | GA | AG | AA | CC | CC | AA | AA | AG | AG | GA | 0.4545454545 |
| **Cónico** | -98.27472222 | 19.99861111 | GG | AA | AA | AA | AA | GG | AA | CC | CC | GG | AA | AG | AA | AA | 0.6818181818 |
| **Cónico** | -99.79111111 | 19.56944444 | GG | AA | AA | CC | AA | AG | AA | CC | CC | GA | AA | AA | AG | AA | 0.5454545455 |
| **Cónico** | -98.06944444 | 19.39388889 | AG | AA | AA | CA | GA | AG | AA | CC | AC | AA | GA | GG | AA | AA | 0.5 |
| **Cónico** | -100.1675028 | 19.35202222 | GG | GA | AA | CA | GG | AG | AA | AA | CC | GA | GA | AG | AA | GA | 0.5454545455 |
| **Cónico** | -100.0255556 | 19.49083333 | AG | AA | AA | CA | GG | GG | AA | AC | CC | GA | AA | AG | AG | GA | 0.5 |
| **Comiteco** | -91.97702778 | 16.19611111 | AA | GG | AA | AA | AA | AG | CA | AC | CC | GA | GG | AG | AG | GA | 0.6363636364 |
| **Comiteco** | -93.09966667 | 16.62313889 | AG | GA | AA | CC | AA | AG | CC | CC | CC | AA | AA | GG | AG | GG | 0.5909090909 |
| **Comiteco** | -92.02008333 | 16.24877778 | AG | GA | AA | CA | GA | AA | AA | AC | CC | GA | GA | GG | AA | AA | 0.6818181818 |

**Supplementary Table 8**. Geographical coordinates, genotype for putative adaptive SNPs, and estimated adaptive scores for maize landraces accessions in Mexico. Detailed information on landrace and known occurrences can be obtained at www.biodiversidad.gob.mx/usos/maices/maiz.html

| Landrace | Longitude | Latitude | PZE.100001195 | PZE.108035280 | PZE.109096578 | SYN27200 | SYN32365 | SYN37470 | PZE.101131254 | PZE.103059706 | PZE.104026211 | PZE.104044827 | PZE.105090166 | PZE.106008139 | PZE.109029456 | PZE.104055811 | Adaptive score |
|---|---|---|---|---|---|---|---|---|---|---|---|---|---|---|---|---|---|
| **Comiteco** | -91.97488889 | 16.23313889 | AG | GA | AA | CA | AA | GG | AA | CC | CC | GA | AA | GG | AA | GA | 0.5909090909 |
| **Comiteco** | -91.93747222 | 16.19525 | AG | GA | AA | AA | AA | GG | AA | CC | CC | AA | AA | AG | AA | AA | 0.5909090909 |
| **Complejo Serrano de Jalisco** | -103.6666667 | 19.93333333 | AA | GG | AA | AA | GA | GG | CA | AC | CC | GG | GA | GG | AG | GG | 0.6363636364 |
| **Complejo Serrano de Jalisco** | -103.43833 | 19.92833 | GG | GA | AA | CA | GA | AG | CA | CC | AC | AA | AA | AA | AA | GG | 0.4090909091 |
| **Conejo** | -98.74311111 | 17.78013889 | GG | GA | AA | AA | AA | AG | CA | AC | CC | GA | GA | GG | AG | AA | 0.7727272727 |
| **Conejo** | -98.74188889 | 17.77797222 | AG | GA | AA | AA | GA | GG | CA | AC | CC | GA | AA | AG | GG | GA | 0.6363636364 |
| **Conejo** | -98.68102778 | 17.88563889 | AA | GA | AA | AA | GG | GG | CC | AC | CC | GA | GA | AG | AG | AA | 0.6363636364 |
| **Conejo** | -98.65641667 | 17.74525 | GG | GG | AA | AA | AA | GG | CA | AA | CC | AA | GA | GG | AA | AA | 0.7272727273 |
| **Conico_Norteno** | -106.63225 | 28.49672222 | AG | GA | AA | CC | AA | GG | AA | AA | CC | GA | GA | AG | GG | AA | 0.5909090909 |
| **Conico_Norteno** | -106.6291111 | 28.49488889 | AA | GG | AA | CC | GA | AG | CA | AC | CC | GA | GA | AG | GG | AA | 0.5909090909 |
| **Conico_Norteno** | -106.6555 | 28.51872222 | GG | AA | AA | CA | AA | AA | AA | AC | AC | AA | AA | AG | AA | AA | 0.6363636364 |
| **Coscomatepec** | -97.56488889 | 19.96466667 | GG | GG | AA | AA | AA | GG | AA | AC | AC | AA | AA | GG | AG | GA | 0.5909090909 |
| **Coscomatepec** | -96.78333333 | 18.5 | AG | GG | AA | AA | AA | AG | AA | CC | CC | GA | AA | GG | AG | GA | 0.6818181818 |
| **Coscomatepec** | -97.45566667 | 20.12658333 | GG | GA | AA | CA | AA | GG | AA | CC | CC | GG | GA | GG | GG | AA | 0.6363636364 |
| **Cristalino_de_Chihuahua** | -106.8788889 | 28.19322222 | AG | GA | AA | CA | AA | AG | AA | CC | CC | AA | AA | AA | AA | AA | 0.5454545455 |
| **Cristalino_de_Chihuahua** | -107.4748056 | 28.50283333 | AG | GG | AA | CC | GA | AG | AA | AC | AC | AA | AA | AG | AA | GA | 0.4545454545 |
| **Dulce** | -108.5333333 | 28.13333333 | AG | GA | AA | CA | GA | GG | CC | CC | AC | GA | AA | GG | GG | AA | 0.6363636364 |
| **Dulcillo del Noroeste** | -108.9246667 | 28.53702778 | AG | AA | AA | AA | GA | GG | AA | AC | CC | AA | GA | GG | GG | AA | 0.5909090909 |
| **Dulcillo del Noroeste** | -109.106 | 28.613 | AA | GG | AA | AA | GA | AG | AA | CC | CC | AA | AA | GG | GG | AA | 0.6363636364 |
| **Dzit-Bacal** | -92.97972222 | 16.03225 | GG | GG | AA | CA | AA | GG | CC | AC | AC | GG | GG | GG | GG | GA | 0.6363636364 |
| **Dzit-Bacal** | -93.00194444 | 16.36402778 | AG | GG | AA | CA | AA | AG | AA | AC | CC | AA | AA | GG | GG | AA | 0.6818181818 |
| **Dzit-Bacal** | -93.45688889 | 16.75177778 | AA | GG | AA | CA | AA | AG | AA | AC | AC | GG | AA | AG | GG | GA | 0.6363636364 |
| **Elotero de Sinaloa** | -105.5463889 | 22.88972222 | GG | AA | AA | CA | GA | AG | AA | CC | CC | AA | AA | AA | AG | GA | 0.4545454545 |
| **Elotero de Sinaloa** | -105.8916667 | 23.40611111 | AG | GA | AA | AA | GA | GG | AA | CC | CC | GA | GA | GG | AA | AA | 0.5909090909 |
| **Elotero de Sinaloa** | -106.0816667 | 23.43083333 | AA | GG | AA | CA | GA | GG | AA | AC | AC | GA | AA | GG | AA | GG | 0.5 |
| **Elotero de Sinaloa** | -106.4261111 | 23.84611111 | AG | GG | AA | CA | GG | GG | CA | AA | CC | GG | GA | AG | AG | GA | 0.5909090909 |
| **Elotero de Sinaloa** | -105.8308333 | 23.4575 | AG | GG | AA | AA | GG | AG | CA | AC | CC | GG | GA | GG | GG | AA | 0.7272727273 |
| **Elotes Conicos** | -99.79111111 | 19.56944444 | AG | AA | AA | CC | GG | GG | AA | CC | AA | AA | AA | GG | AG | GA | 0.3181818182 |
| **Elotes Conicos** | -97.9725 | 19.32166667 | AG | AA | AA | CC | GA | AG | AA | CC | AC | AA | AA | AG | AG | GG | 0.3636363636 |
| **Elotes Conicos** | -97.92472222 | 19.19055556 | AG | AA | AA | CA | GA | GG | AA | AC | CC | GA | AA | AG | AA | GA | 0.5454545455 |
| **Elotes Conicos** | -100.0255556 | 19.49083333 | AG | GG | AA | AA | AA | AG | CA | AA | AC | GG | GG | GG | GG | GA | 0.7272727273 |
| **Elotes Conicos** | -98.37388889 | 20.35888889 | GG | GG | AA | AA | AA | AG | AA | CC | CC | GG | GA | GG | AG | GA | 0.6818181818 |
| **Elotes Conicos** | -98.37555556 | 20.35472222 | AG | GG | AA | AA | GA | AG | CA | CC | AC | GA | GG | GG | AG | GG | 0.5 |
| **Elotes Conicos** | -98.38333333 | 20.3975 | AG | AA | AA | CA | GA | GG | AA | AC | AA | GG | GG | AG | GG | AA | 0.4545454545 |
| **Elotes Conicos** | -99.05027778 | 19.19166667 | GG | AA | AA | CA | AA | GG | AA | CC | CC | GA | GA | AG | AA | GA | 0.5 |
| **Elotes Conicos** | -98.31555556 | 19.4875 | AG | GA | AA | CA | GA | GG | AA | CC | CC | GA | GA | AG | AG | GA | 0.4545454545 |
| **Elotes Conicos** | -98.37555556 | 20.35472222 | AG | AA | AA | CA | AA | GG | AA | CC | CC | GA | GA | AA | AG | AA | 0.5 |
| **Elotes Conicos** | -98.77233333 | 19.05555556 | GG | GA | AA | CA | GA | GG | AA | CC | AA | AA | GA | GG | AA | GA | 0.3636363636 |
| **Elotes Conicos** | -98.04722222 | 19.33861111 | AG | AA | AA | AA | GG | AG | AA | AC | AC | GA | AA | AA | AG | GG | 0.4545454545 |
| **Elotes Conicos** | -99.92944444 | 19.70611111 | GG | AA | AA | AA | GG | GG | AA | AC | AC | AA | GA | GG | AA | GA | 0.4545454545 |
| **Elotes Conicos** | -98.80666667 | 19.10861111 | AG | AA | AA | CA | AA | AA | AA | CC | CC | AA | GA | GG | AG | GA | 0.5909090909 |
| **Elotes Occidentales** | -100.7658333 | 20.74416667 | GG | AA | AA | CA | AA | AG | AA | AC | CC | AA | GA | GG | AG | AA | 0.6363636364 |
| **Elotes Occidentales** | -100.1083333 | 19.04388889 | AG | GA | AA | AA | AA | AG | CA | CC | CC | GA | GG | GG | AG | AA | 0.6818181818 |
| **Elotes Occidentales** | -100.8147222 | 20.78444444 | GG | GG | AA | AA | GA | AG | CC | AC | AC | AA | GA | GG | GG | GA | 0.6363636364 |
| **Elotes Occidentales** | -100.9355556 | 20.78722222 | AG | GA | AA | AA | GG | AG | AA | CC | CC | GA | GG | AA | AA | GA | 0.4090909091 |
| **Gordo** | -108.7097222 | 29.84027778 | AG | GG | AA | AA | GA | AG | AA | AA | CC | AA | GA | GG | AA | GG | 0.5909090909 |
| **Gordo** | -108.7097222 | 29.84027778 | AG | GA | AA | AA | AA | GG | AA | CC | AC | GA | AA | GG | AA | GA | 0.5909090909 |
| **Jala** | -104.4286944 | 21.10166667 | AG | GG | AA | CA | GA | AA | AA | CC | AC | GA | AA | GG | AG | AA | 0.6363636364 |
| **Jala** | -104.4405556 | 21.1 | AA | GA | AA | AA | AA | GG | AA | CC | CC | GA | AA | GG | AA | GA | 0.6363636364 |

**Supplementary Table 8**. Geographical coordinates, genotype for putative adaptive SNPs, and estimated adaptive scores for maize landraces accessions in Mexico. Detailed information on landrace and known occurrences can be obtained at www.biodiversidad.gob.mx/usos/maices/maiz.html

| Landrace | Longitude | Latitude | PZE.100001195 | PZE.108035280 | PZE.109096578 | SYN27200 | SYN32365 | SYN37470 | PZE.101131254 | PZE.103059706 | PZE.104026211 | PZE.104044827 | PZE.105090166 | PZE.106008139 | PZE.109029456 | PZE.104055811 | Adaptive score |
|---|---|---|---|---|---|---|---|---|---|---|---|---|---|---|---|---|---|
| **Jala** | -104.4286944 | 21.10166667 | GG | GG | AA | AA | GA | AG | AA | AA | AC | GA | GA | AG | AG | GA | 0.5909090909 |
| **Jala** | -104.4399794 | 21.07805 | GG | GA | AA | CA | GA | AG | AA | AC | AC | AA | AA | GG | AG | AA | 0.5909090909 |
| **Mushito** | -99.805 | 21.2775 | AA | AA | AA | CC | AA | AG | AA | AC | CC | GA | AA | AG | GG | GA | 0.5909090909 |
| **Mushito** | -111.9864167 | 26.89047222 | AG | 0 | GA | AA | AA | GG | AA | CC | CC | GA | GG | GG | GG | AA | 0.5454545455 |
| **Mushito** | -100.13722 | 21.3575 | AG | AA | AA | CA | GA | AA | CA | AC | CC | AA | AA | GG | AG | GA | 0.6818181818 |
| **Nal-tel de Altura** | -98.21666667 | 16.3 | GG | GG | AA | AA | AA | GG | CA | AA | CC | GG | 0 | GG | AA | AA | 0.7727272727 |
| **Nal-tel de Altura** | -96.73333333 | 18.01666667 | AG | GA | AA | CA | GA | GG | CA | AC | CC | AA | GG | AA | AA | AA | 0.4545454545 |
| **Nal-tel de Altura** | -91.97688889 | 16.23319444 | GG | AA | AA | CA | GA | GG | CA | AC | AC | GA | GA | AA | AA | GA | 0.4545454545 |
| **Nal-tel de Altura** | -91.91638944 | 16.21563917 | GG | AA | AA | CC | GG | GG | AA | CC | CC | AA | AA | AG | AG | GA | 0.3636363636 |
| **Nal-tel de Altura** | -97.06666667 | 18.85 | AG | GA | AA | AA | GA | GG | AA | CC | CC | AA | AA | GG | AG | GA | 0.5454545455 |
| **Oloton** | -92.3145 | 15.36608333 | GG | GA | GA | CA | GA | AG | AA | AC | CC | AA | AA | GG | AA | AA | 0.5909090909 |
| **Oloton** | -92.29358333 | 15.36011111 | AG | AA | AA | CA | AA | AG | AA | CC | AC | AA | GA | GG | AG | GA | 0.5 |
| **Oloton** | -92.54222222 | 16.63975 | AG | AA | AA | CA | GA | AG | AA | CC | CC | GG | GG | AG | AA | AA | 0.5454545455 |
| **Oloton** | -92.18767778 | 15.29614444 | AG | AA | AA | CC | AA | AA | AA | CC | CC | GA | AA | AG | GG | AA | 0.6363636364 |
| **Olotillo** | -98.43611111 | 21.13333333 | GG | AA | AA | AA | AA | GG | AA | AC | CC | AA | AA | GG | GG | AA | 0.6818181818 |
| **Olotillo** | -98.34638889 | 21.06388889 | GG | GA | AA | AA | GA | AA | AA | AC | CC | AA | GG | GG | AG | GA | 0.5909090909 |
| **Olotillo** | -92.97972222 | 16.03225 | GG | AA | GA | CA | AA | AA | CA | AC | CC | GA | AA | AG | AG | AA | 0.7272727273 |
| **Olotillo** | -93.00194444 | 16.36402778 | AA | AA | AA | CA | AA | AG | CA | CC | CC | GA | GA | GG | AA | GA | 0.6363636364 |
| **Olotillo** | -92.216925 | 14.90176111 | AG | GG | AA | CA | AA | AG | CA | AC | CC | GA | AA | AG | GG | GG | 0.6363636364 |
| **Olotillo** | -92.2045 | 14.87376389 | AA | AA | AA | CC | AA | GG | AA | AC | CC | GA | GA | GG | AG | GA | 0.5454545455 |
| **Onaveño** | -109.68 | 29.80894444 | AA | GA | AA | AA | AA | GG | CA | AA | CC | GG | GG | GG | AG | GG | 0.6818181818 |
| **Onaveño** | -108.8240833 | 27.17294444 | GG | GA | AA | AA | AA | GG | AA | CC | CC | AA | GG | AG | AA | AA | 0.5 |
| **Palomero de Chihuahua** | -106.4436667 | 26.35983333 | AA | GG | AA | CA | GA | AA | AA | CC | CC | GA | GA | AG | GG | GG | 0.5 |
| **Palomero_Toluqueno** | -99.71715556 | 19.79833333 | AG | AA | AA | CC | AA | GG | AA | AC | AC | AA | AA | GG | AG | GA | 0.5 |
| **Pepitilla** | -100.1686111 | 19.0525 | GG | GA | AA | AA | GA | GG | AA | CC | CC | GG | AA | AG | AG | AA | 0.6363636364 |
| **Pepitilla** | -100.1686111 | 19.0525 | GG | GA | AA | AA | GG | GG | AA | CC | CC | GG | GA | GG | AG | AA | 0.5909090909 |
| **Pepitilla** | -100.1083333 | 19.04388889 | GG | GA | AA | CC | GA | AG | AA | AC | CC | AA | GA | AG | AA | GA | 0.4545454545 |
| **Pepitilla** | -100.1083333 | 19.04388889 | AG | GA | AA | CA | AA | GG | CA | CC | CC | AA | GG | AG | AA | GA | 0.4545454545 |
| **Raton** | -100.7042778 | 25.68125 | GG | GG | AA | AA | AA | GG | CA | CC | CC | GA | GA | GG | GG | GG | 0.5909090909 |
| **Raton** | -99.25505556 | 24.12780556 | AG | GG | AA | CA | AA | GG | AA | AC | CC | AA | GA | AG | GG | GG | 0.4545454545 |
| **Raton** | -99.02108333 | 24.29011111 | GG | AA | AA | AA | AA | GG | AA | AA | CC | GA | AA | AG | GG | AA | 0.7272727273 |
| **Reventador** | -108.9119444 | 26.84361111 | AG | GG | AA | AA | GA | GG | AA | AC | CC | GG | AA | GG | AG | GA | 0.6818181818 |
| **Reventador** | -110.2101111 | 29.79355556 | GG | GA | AA | CA | AA | AG | CA | AC | AC | GA | AA | GG | AG | AA | 0.7272727273 |
| **Tablilla de Ocho** | -106.6324722 | 28.32019444 | AG | GG | AA | AA | GA | AG | CC | AA | AC | AA | AA | AA | GG | GA | 0.6363636364 |
| **Tablilla de Ocho** | -106.0163611 | 26.94213889 | AG | GA | AA | CA | AA | AA | AA | AC | AC | AA | AA | 0 | GG | AA | 0.5909090909 |
| **Tabloncillo** | -105.6068056 | 22.94927778 | GG | GG | AA | AA | GA | AG | AA | AC | CC | GA | GA | AG | AG | GA | 0.5909090909 |
| **Tabloncillo** | -109.2448056 | 27.82702778 | GG | GG | AA | AA | GA | AG | AA | AC | CC | AA | AA | AG | AA | AA | 0.6363636364 |
| **Tabloncillo** | -107.5618889 | 25.40125 | GG | GA | AA | AA | GG | AA | AA | AC | AC | GA | GA | AG | AG | GA | 0.5454545455 |
| **Tabloncillo** | -108.9255278 | 28.41063889 | AG | GA | AA | AA | AA | AG | AA | AC | CC | AA | AA | GG | AG | AA | 0.7272727273 |
| **Tabloncillo_Perla** | -105.39025 | 21.99097222 | AA | GG | AA | AA | GA | GG | AA | CC | CC | GG | AA | GG | GG | GA | 0.6363636364 |
| **Tabloncillo_Perla** | -105.2216111 | 21.94602778 | AA | GG | AA | AA | GG | AA | CA | AC | CC | GA | AA | GG | AG | GA | 0.7272727273 |
| **Tabloncillo_Perla** | -105.1493611 | 20.87416667 | AG | GG | AA | AA | AA | GG | AA | AC | CC | GG | AA | AG | GG | AA | 0.7272727273 |
| **Tehua** | -93.206 | 17.19680556 | AA | AA | AA | CA | GA | AG | AA | AC | CC | AA | GA | GG | AA | GG | 0.5 |
| **Tehua** | -93.17527778 | 17.21630556 | AA | GA | AA | CC | GA | AG | AA | CC | CC | AA | AA | GG | AG | AA | 0.5454545455 |
| **Tepecintle** | -92.08888889 | 17.32222222 | GG | AA | AA | CC | AA | GG | AA | AC | CC | GA | GA | AG | AG | GG | 0.4545454545 |
| **Tepecintle** | -92.31925 | 15.36652778 | AA | AA | AA | CA | GA | AG | AA | AC | CC | GA | AA | GG | AA | AA | 0.6818181818 |
| **Tepecintle** | -92.19657222 | 14.82029722 | GG | GG | AA | AA | AA | AG | AA | AC | CC | GG | GG | 0 | GG | GG | 0.5454545455 |
| **Tepecintle** | -92.46687222 | 17.25071111 | AA | GA | AA | AA | AA | AG | CA | AA | CC | GG | AA | GG | GG | GG | 0.8181818182 |
| **Tuxpeno** | -92.81925 | 16.04422222 | GG | GG | AA | CA | AA | GG | AA | AC | AC | GG | AA | AG | GG | GG | 0.5454545455 |

**Supplementary Table 8**. Geographical coordinates, genotype for putative adaptive SNPs, and estimated adaptive scores for maize landraces accessions in Mexico. Detailed information on landrace and known occurrences can be obtained at www.biodiversidad.gob.mx/usos/maices/maiz.html

| Landrace | Longitude | Latitude | PZE.100001195 | PZE.108035280 | PZE.109096578 | SYN27200 | SYN32365 | SYN37470 | PZE.101131254 | PZE.103059706 | PZE.104026211 | PZE.104044827 | PZE.105090166 | PZE.106008139 | PZE.109029456 | PZE.104055811 | Adaptive score |
|---|---|---|---|---|---|---|---|---|---|---|---|---|---|---|---|---|---|
| **Tuxpeno** | -106.1319444 | 23.25388889 | GG | GG | AA | AA | GA | AG | AA | CC | CC | GA | GG | AG | GG | GA | 0.5 |
| **Tuxpeno** | -99.72272222 | 24.75586111 | AG | GG | AA | AA | GA | AG | AA | CC | CC | GA | GA | AA | GG | GG | 0.4545454545 |
| **Tuxpeno** | -92.68894972 | 16.1194325 | AA | GA | AA | CC | AA | AG | AA | AC | AC | AA | GG | GG | AG | AA | 0.5 |
| **Tuxpeno_Norteno** | -99.53447222 | 24.85894444 | AG | AA | AA | AA | AA | AG | AA | AC | CC | GA | GA | GG | GG | GA | 0.6818181818 |
| **Tuxpeno_Norteno** | -100.2275 | 18.66027778 | GG | GA | AA | AA | AA | GG | AA | CC | CC | GA | AA | AG | AG | GG | 0.5454545455 |
| **Vandeno** | -110.2121667 | 29.79602778 | GG | AA | AA | AA | GA | GG | AA | CC | CC | GA | AA | GG | AG | GG | 0.5454545455 |
| **Vandeno** | -92.97783333 | 16.04222222 | AA | GG | AA | CA | GA | AG | CA | AC | CC | GA | AA | GG | GG | GA | 0.6818181818 |
| **Vandeno** | -92.69643389 | 16.12061694 | AG | GG | AA | CC | GG | GG | AA | CC | CC | GA | AA | AG | AG | GA | 0.4090909091 |
| **Vandeno** | -92.46687222 | 17.25071111 | GG | GG | AA | AA | GA | GG | AA | AC | CC | GG | GA | GG | GG | GA | 0.6363636364 |
| **Zamorano Amarillo** | -104.45 | 19.6 | AA | GG | AA | AA | GA | GG | CA | AA | AC | GA | GA | GG | GG | GG | 0.5909090909 |
| **Zamorano Amarillo** | -102.7166667 | 19.96666667 | AG | GA | AA | AA | GG | AG | CA | AC | AC | AA | GA | AG | AG | GA | 0.5 |
| **Zamorano Amarillo** | -104.6333333 | 19.71666667 | GG | GG | AA | CC | AA | AA | AA | CC | AC | GG | GA | AG | AG | GG | 0.5 |
| **Zapalote Grande** | -93.00194444 | 16.36402778 | AG | GG | AA | AA | GA | GG | CC | AC | CC | AA | AA | AG | AG | AA | 0.6818181818 |
| **Zapalote_Chico** | -93.00194444 | 16.36402778 | AG | GA | AA | AA | GA | GG | CC | AC | CC | AA | AA | AG | AG | AA | 0.6818181818 |

**Supplementary Table 9.** Geographic distribution, environmental information and most common use of different maize landraces in Mexico with the potential for crop improvement (ref. 14). Detailed information on landraces can be obtained at www.biodiversidad.gob.mx/usos/maices/maiz.html

| Landrace | Distribution | Enviroment | Use |
|---|---|---|---|
| Azul | Chihuahua | Not reported in ref. 14 | Food source |
| Conejo | Oaxaca, Guerrero and Michoacan | tempered and dry | Not reported in ref. 14 |
| Cónico Norteño | Guanajuato, Chihuahua, Zacatecas, Durango, Aguascalientes, San Luis Potosi, Coahuila and Nuevo Leon | low precipitation and extreme temperatures | Improvement for drought resistance |
| Elotes Occidentales | Nayarit, Jalisco, Michoacan, Guanajuato, Zacatecas, San Luis Potosi, Morelos, Puebla, Guerrero and Oaxaca | Not reported in ref. 14 | Food source |
| Olotillo | Chiapas, Oaxaca y Guerrero, Veracruz, Puebla, Hidalgo and San Luis Potosi | Humid and dry tropics, poor soils | Food source |
| Raton | Tamaulipas, Nuevo León, Coahuila, Chihuahua, Durango, Zacatecas, San Luis Potosí, Veracruz, Guerrero and Morelos | subtropics y deserts, needs low humidity, | Improvement for highland races |
| Tabloncillo | Michoacán, Jalisco, Nayarit y Sinaloa, Sonora, Chihuahua and Baja California Sur | Not reported in ref. 14 | Improvement, food source |
| Tabloncillo perlas | Sinaloa, Nayarit, Jalisco, Colima, Michoacan, Sonora and Baja California Sur | Low lands and dry, thin soils | Food source |
| Tepecintle | Oxaca y Chiapas | Laderas | Food source |
| Tuxpeño Norteño | North, Center of Mexico | Subtropics, season, | Improvement |
| Vandeño | Pacific coast | Seasonaility and low precipitation | Improvement |
